# Co-expression-based models improve eQTL predictions and highlight novel transcriptome-wide genes associated with schizophrenia

**DOI:** 10.64898/2026.02.10.704353

**Authors:** Fabiana Rossi, Leonardo Sportelli, Gianluca C. Kikidis, Giulia Grassi, Fabio Di Camillo, Alessandro Bertolino, Giuseppe Blasi, Christopher Borcuk, Daniela Fusco, Thomas M. Hyde, Joel E. Kleinman, Davide Marnetto, Silvia Pellegrini, Antonio Rampino, Benedetto Vitiello, Stephan Ripke, Alice Braun, Julia Kraft, Sintia Iole Belangero, Paulo R. Menezes, Celso Arango, James T.R. Walters, Michael C. O’Donovan, Michael J. Owen, David Braff, Aiden Corvin, Derek W. Morris, Enrico Domenici, Jim van Os, Esref Atbaşoğlu, Meram C. Saka, Marta Di Forti, Bernhard T. Baune, Carlos N. Pato, Andrew McQuillin, Vera Golimbet, Nikolay Kondratyev, Valentina Escott-Price, Anna Gareeva, Elza Khusnutdinova, Jorge A. Cervilla, Margarita Rivera, Claudine Laurent-Levinson, Alessandro Serretti, Ole A. Andreassen, David St. Clair, Todd Lencz, Anil K. Malhotra, Nina S. McCarthy, Bryan J. Mowry, Dan Rujescu, Ina Giegling, Annette M. Hartmann, Bettina Konte, Markus M. Nöthen, Marcella Rietschel, George Kirov, Patrick F. Sullivan, Tracey L. Petryshen, Thomas Werge, Andrew McIntosh, Tõnu Esko, Erik G. Jönsson, Hannelore Ehrenreich, Brien P. Riley, Douglas F. Levinson, Joseph D. Buxbaum, Elvira Bramon, Roel A. Ophoff, Rolf Adolfsson, Eli A. Stahl, Bart P.F. Rutten, Sinan Guloksuz, Cristina M. Del-Ben, Florence Thibaut, Daniel R. Weinberger, Giulio Pergola

## Abstract

Non-coding genetic variants statistically associated with complex heritability phenotypes are thought to act primarily through transcriptome regulatory mechanisms. Predictions of gene expression in tissue like the human brain traditionally rely primarily on *cis*-eQTLs. Here, we introduce INGENE and MODULE, *trans*-eQTLs models designed to enhance the prediction of gene expression by capturing the collective impact of candidate *trans*-eQTLs acting within co-expression networks. Exploiting RNA-seq data in six post-mortem brain regions (amygdala, caudate nucleus, dorsal/subgenual anterior cingulate cortex, dorsolateral prefrontal cortex, and hippocampus), we validate our models on two testing datasets, demonstrating increased gene predictability compared to both an original *cis*-based model and to EpiXcan, the leading benchmark in *cis*-model performance. Integration of *cis*- and *trans*-predictions significantly improves gene-level expression imputation (MLE α= 0.05) for 18,744 genes across the six brain regions considered. Applying *cis* and *trans* models to PGC wave 3 genotypes identifies 766 SCZ-associated genes across brain regions (pFDR < .01), emphasizing the complementary nature of *cis* and *trans* predictions in trait association discovery. Of these genes, 641 represent novel transcriptome-wide associations with schizophrenia, highlighting the role of *trans*-heritability and genetic interactions underlying risk for this disorder, in addition to further supporting 125 previous candidates.

## Introduction

Schizophrenia (SCZ) is a severe psychiatric disorder with complex heritability estimated between 60% and 80% in twin studies^1,2^. Genome-wide association studies (GWASs) indicate that most common variation contributing to SCZ risk are non-coding single nucleotide polymorphisms (SNPs)^2^. While GWAS provide valuable insight into genetic associations with disease susceptibility, additional functional genomics approaches are required to clarify the molecular mechanisms of illness^3^. In the case of SCZ, larger sample sizes have enhanced the statistical power to identify significant loci^4,5^, but the relative increment in explained variance by newly discovered loci remains modest^6^, reflecting the challenge of resolving the molecular mechanisms linking non-coding variants to disease^7,8^.

As gene expression is intermediate between the DNA sequence and phenotype, mRNA is the proximal functional unit exerting non-coding genetic effects on illness liability^9,10^. Integrating GWAS with gene expression data in postmortem brain studies has therefore become central to functional genomics in psychiatric disease^11–16^. However, most noncoding risk variants do not colocalize with known expression quantitative trait loci (eQTLs)^7,17–19^, limiting the interpretability of GWAS findings. Transcriptome-wide association studies (TWAS)^12^ were designed to address these challenges by imputing genetically regulated gene expression and correlating it with phenotypes. In TWAS, two main approaches are commonly employed: the direct prediction of expression within genotyped samples (gene-level TWAS) and the indirect estimation of associations between predicted expression and traits while considering linkage disequilibrium among SNPs (SNP-level TWAS). Both approaches identify gene-level trait associations while reducing the multiple-testing burden compared to SNP-based GWAS^1314^. Traditional TWAS frameworks, such as PrediXcan^13^ and related methods^14,20,21^, have so far mainly relied on *cis*-eQTLs, i.e., variants within ± 1Mb of a gene, under the assumption that local regulation dominates genetic control of expression. Yet, *cis*-eQTLs account for only a fraction of the heritability of gene expression^2,22,23^, and many genes depleted of *cis*-eQTLs are likely influenced by *trans*-acting mechanisms^24^. Such effects, mediated through transcriptional regulators, chromatin modifiers, or co-expressed genes, are hypothesized to explain up to 70% of the heritable interindividual variability in gene expression^24–28^.

The importance of *trans*-regulation may be particularly evident in SCZ, where disease-associated variants act through coordinated transcriptional programs across brain regions and developmental stages^29,30^. As *trans*-effects are weaker than *cis*-effects, *trans*-eQTL detection per se requires large sample sizes^13^. Despite considerable progress, *trans*-eQTL characterization remains an open challenge^17^. Considering the large fraction of heritability that *trans*-eQTLs are hypothesized to explain, identifying them is a research priority.

Although risk-associated variants are scattered across the genome, their functional correlates may converge^31,32^ at molecular- or system-level processes^30^. For example, several studies have reported the convergence of genes associated with SCZ into co-expressed gene sets^33–39^. Such molecular environments of risk vary dynamically across brain regions and over time^31,32^ and even at the cell type level^37,40,41^. Gene co-expression models may also identify transcription factors and regulatory elements acting upon multiple genes^20,32,42^, and these findings can be validated *in vitro*^39,43^.

Several previous studies have leveraged SNPs proximal to genes in co-expressed sets, named co-expression based eQTL (co-eQTLs), to generate system level polygenic scores to translate co-expression into phenotypic predictions^20,38,44–47^ ^48^ ^20,44,49,50^. The rationale of this approach is that shared variance in the expression of multiple genes may reflect risk biology in a systems context and may also help identifying latent co-regulation mechanisms within gene expression modules, e.g., associated with regulatory elements^39,51,52^. Because it limits the search space to co-expression gene sets, this approach to potential *trans*-eQTLs differs from genome-wide screenings, which often fail due to the burden of multiple comparisons^13^.

Building on this rationale, we developed two predictive frameworks to quantify *trans*-regulatory contributions to gene-level expression: 1) the Imputed Network Gene-Expression *trans*- eQTL (INGENE), and 2) the MODule quantitative trait Loci Eigengene (MODULE). INGENE models the *trans-*effects of *cis*-regulated co-expression partners on a target gene, whereas MODULE identifies SNPs associated with the eigenvalue (first principal component) of the target gene’s co-expression module. In addition to our *trans* algorithms, we also implemented a standard *cis*-based model (CIS) and an EpiXcan model^13,14^.

We assessed the potential of INGENE and MODULE to increase the number of genetically predictable risk genes compared to *cis*-based models and tested whether the integration of *cis* and *trans* scores enhances predictive accuracy. We also conducted a co-expression-based transcriptomic association study (coTWAS) focusing on SCZ diagnosis in the Psychiatric Genomic Consortium (PGC) wave 3^6^. Figure 1 summarizes the key steps of our study.

**Figure 1.**
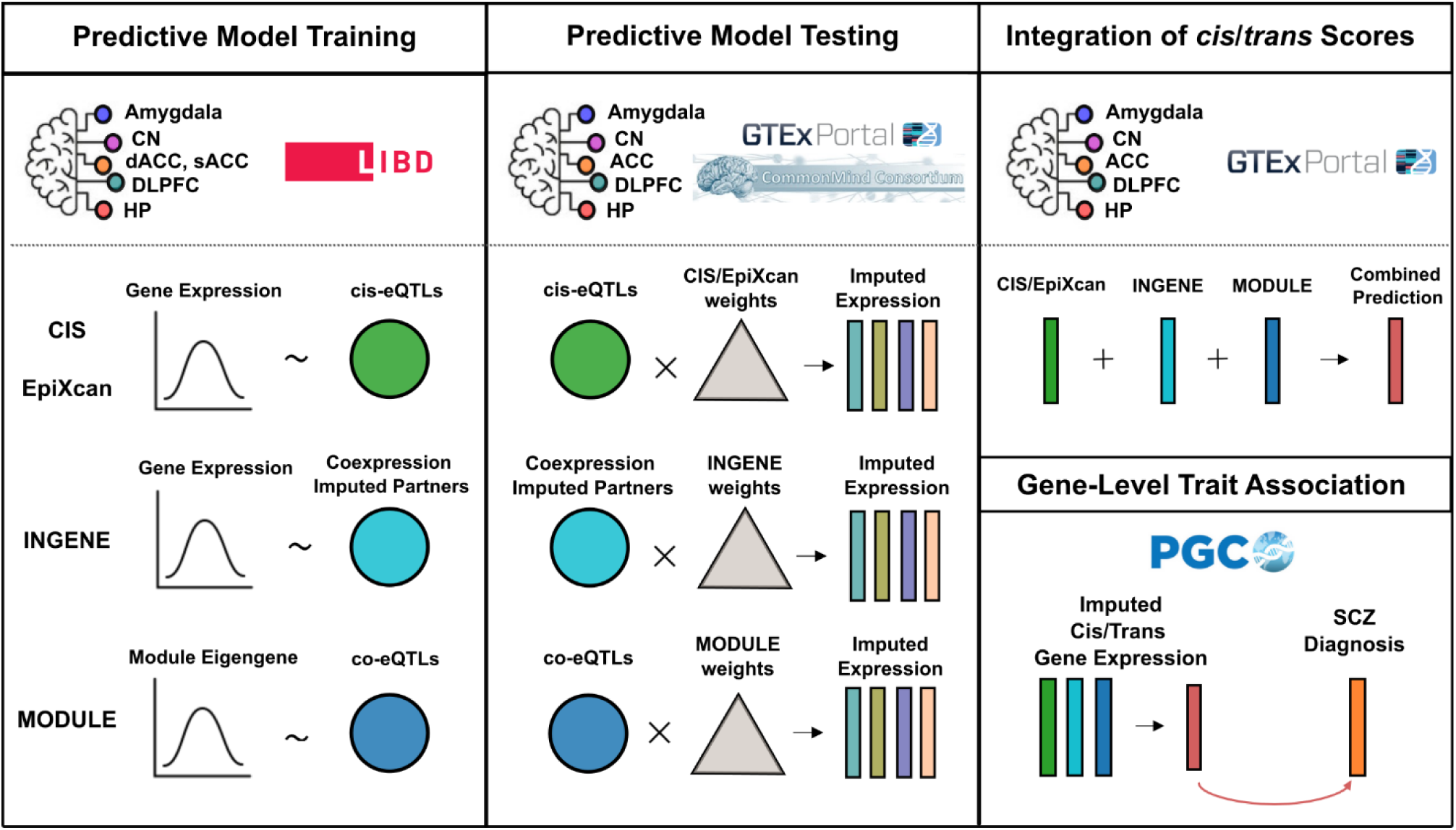
Overview of the analytical pipeline. The pipeline consists of four main stages. **(1) Predictive model training:** Univariate cis-based (CIS, EpiXcan) and trans-based (INGENE, MODULE) gene expression prediction models were trained using Elastic Net regularized regression across six brain regions from the LIBD dataset. **(2) Predictive model testing:** The trained cis- and trans-models were applied to independent genotype datasets (GTEx and CMC) to impute gene-level expression and assess the replicability of cis/trans scores in external cohorts. **(3) Integration of cis/trans scores:** Imputed expression values from CIS/EpiXcan, INGENE, and MODULE models were combined to generate integrated prediction scores. **(4) Gene-level trait association:** The integrated cis/trans-imputed gene expression was tested for association with SCZ diagnosis in the PGC3 cohort using coTWAS.

## Results

### INGENE and MODULE increase the number of predicted genes across brain regions

#### Evaluation of *cis*- and *trans*-model training performance

We used the LIBD brain RNA seq dataset for training as it featured larger sample sizes than GTEx for the brain regions we considered. As *cis*-based benchmarks, we implemented a standard elastic-net model^13^ (CIS) and EpiXcan^14^, which incorporates chromatin-state priors from Roadmap Epigenomics^53^ (https://egg2.wustl.edu/roadmap/web_portal) into elastic-net regression. Across five regions (RoadMap annotations were not available for amygdala), EpiXcan outperformed CIS in both the number of predictable genes (Fig. 2A) and training accuracy (mean adjusted R² of 0.153 *vs.* 0.140; ∼9% relative gain across regions; Fig. S1). Yet the two methods provided complementary coverage: ∼6-8% of genes were uniquely captured by either approach, or CIS outperformed EpiXcan for a subset of shared predictions (Fig. 2B). Given this complementarity and the limited sample sizes available for some regions (Table 2), we retained both models as *cis*-models in downstream analyses to maximize gene representation.

**Figure 2.**
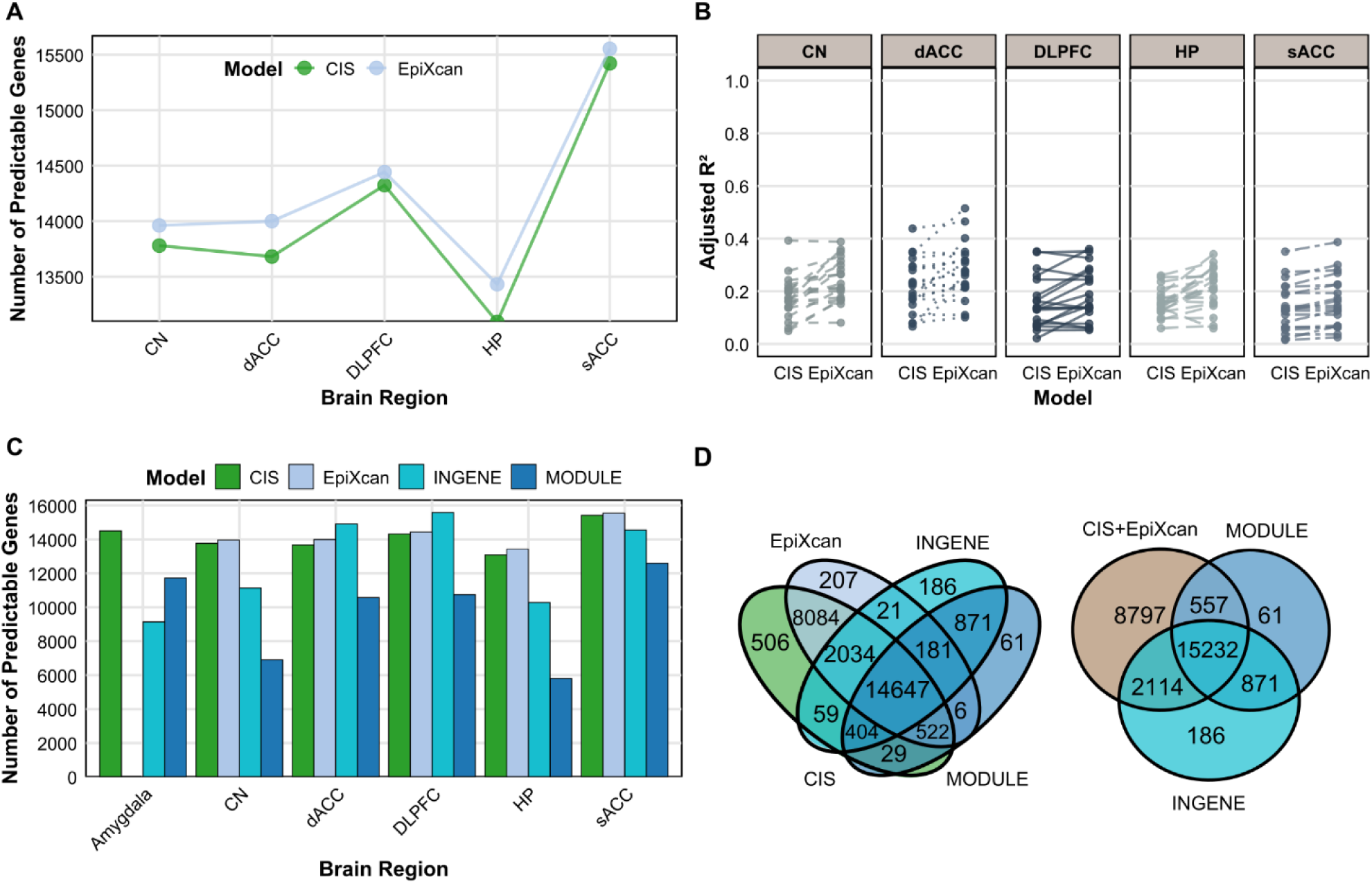
Comparative performance of cis- and trans-prediction models across brain regions. A) Number of predictable genes for CIS (green) and EpiXcan (light blue) across LIBD brain regions. B) Gene-level compar-ison of cross-validation (CV) adjusted R² between CIS and EpiXcan for genes predicted by both models, high-lighting the generally higher predictive accuracy of EpiXcan but also the subset where CIS performs better. C) Number of predictable genes across brain regions for all models: CIS (green), EpiXcan (light blue), IN-GENE (cyan), and MODULE (dark blue). The two trans-aware models substantially expand the pool of im-putable genes relative to cis-only approaches. D) Overlap of predicted genes across models. Left: intersection among CIS, EpiXcan, INGENE, and MODULE. Right: overlap of trans-aware models with the union of cis predictors (CIS+EpiXcan).

We next compared *cis*-predictive models (CIS and EpiXcan) with our *trans*-models, INGENE and MODULE. Both *trans*-models were trained in LIBD and cross-trained in GTEx and the Common Mind Consortium^36^ (CMC) (see Table 2 for individuals’ characteristics), retaining only replicable genes across training datasets (Supplementary Notes: Section 2.1; Fig. S2). This approach reduced false positives and improved stability across cohorts for *trans*-eQTL identification. CIS and EpiXcan were not filtered this way to avoid penalizing *cis*-models, which we showed were sensitive to training sample size (Supplementary Methods: Section 1.5; Supplementary Notes: Section 2.2; Figs. S3-S4).

We quantified model performance during training, bearing in mind that training performance metrics are prone to overfitting and require validation within unseen samples. Within this framework, the models demonstrated different predictive capabilities across regions (Fig. 2C). When pooling data, CIS predicted 26,285 genes, EpiXcan 25,702, INGENE 18,403, and MODULE 16,721 (Fig. 2D left). *Cis*-based models yielded 8,797 unique predictions, (30% of the total), i.e., not shared with *trans*-models, while INGENE and MODULE overlapped considerably, with 16,103 common predictions, representing 96% of MODULE and 88% of INGENE (Fig. 2D right). Figure S5 shows the unfiltered performance comparison. To further explore the regulatory landscape of these *trans-*shared predicted genes, we examined whether they shared regulatory components. Specifically, we considered instances where a SNP acted as a *trans*-eQTL for a MODULE-predicted gene and simultaneously served as a *cis*-eQTL of the co-expression partners that INGENE uses to predict that same gene. We found that 15%-29% of genes predicted by MODULE across the six brain regions exhibited regulatory relationships (Table S1).

We further assessed the cross-validation (CV) performance in terms of adjusted R^2^ on genes commonly predicted by all models across regions. We computed the delta (Δ) *R^2^*CV values between models (Fig. S6) and employed the one-tailed Wilcoxon test to ascertain if these differences were statistically significant. EpiXcan outperformed CIS (mean ΔR² = 0.0103), confirming the expected benefit of incorporating epigenomic priors into *cis*-eQTL modeling. INGENE achieved descriptively higher mean R² than CIS (ΔR² = 0.0241) and EpiXcan (ΔR² = 0.0257), but these differences did not reach statistical significance. In contrast, MODULE exhibited consistently and significantly superior performance, explaining greater variance than all other models, with significant improvements relative to INGENE (ΔR² = 0.0464), CIS (ΔR² = 0.0721), and EpiXcan (ΔR² = 0.0705).

Finally, to contextualize our approach, we compared our models with published *cis*/*trans* frameworks, BGW-TWAS^54^ and MOSTWAS^55^ (see Supplementary Notes: Section 2.3), restricting analyses to the DLPFC, the only common region. In our framework, a gene was classified as *cis* or *trans* whenever the corresponding component contributed to its prediction (Fig. S7A). Notably, ∼59% of the total BGW *cis*/*trans* predictions and ∼96% of the *cis*/*trans* set in our framework were recovered by both approaches (Fig. S7B). This high level of consistency indicates that, even though training- based models are prone to overfitting, our classification of *cis* and *trans* predictions captures regulatory signals robustly aligned with independent frameworks, while extending them to additional genes.

Having established consistency in the training data, we next turned to independent testing datasets from GTEx and CMC to assess the generalizability of our models.

### Evaluation of *cis-* and *trans-*models on a testing dataset

We assessed the replication of all four models—CIS, EpiXcan, INGENE, and MODULE—in the GTEx cohort, which is independent of the LIBD training cohort. This evaluation encompassed the same six brain regions we used for model training. We applied each model to GTEx genotype data to generate predictions, this time ruling out any potential overfitting as they were applied to the testing dataset. Our assessment of predictions was based on two criteria: i) the number of genes imputed with a Pearson’s correlation *r* > 0 and ii) adjusted R^2^>0 between predicted and observed values.

In the testing dataset, CIS predicted a minimum of 4,732 and a maximum of 7,777 genes across different brain regions, while EpiXcan predicted between 4,858 and 7,797 genes. INGENE achieved the broadest coverage, predicting between 5,429 and 10,749 genes. MODULE showed a comparable coverage, with 5,084-10,718 predicted genes across regions (Fig. 3A). Thus, both *trans* models substantially increased the number of predictable genes relative to *cis*-models. For example, INGENE predicted up to 1.8 times more genes than CIS, and up to 1.7 times more than EpiXcan (Fig. S8).

**Figure 3.**
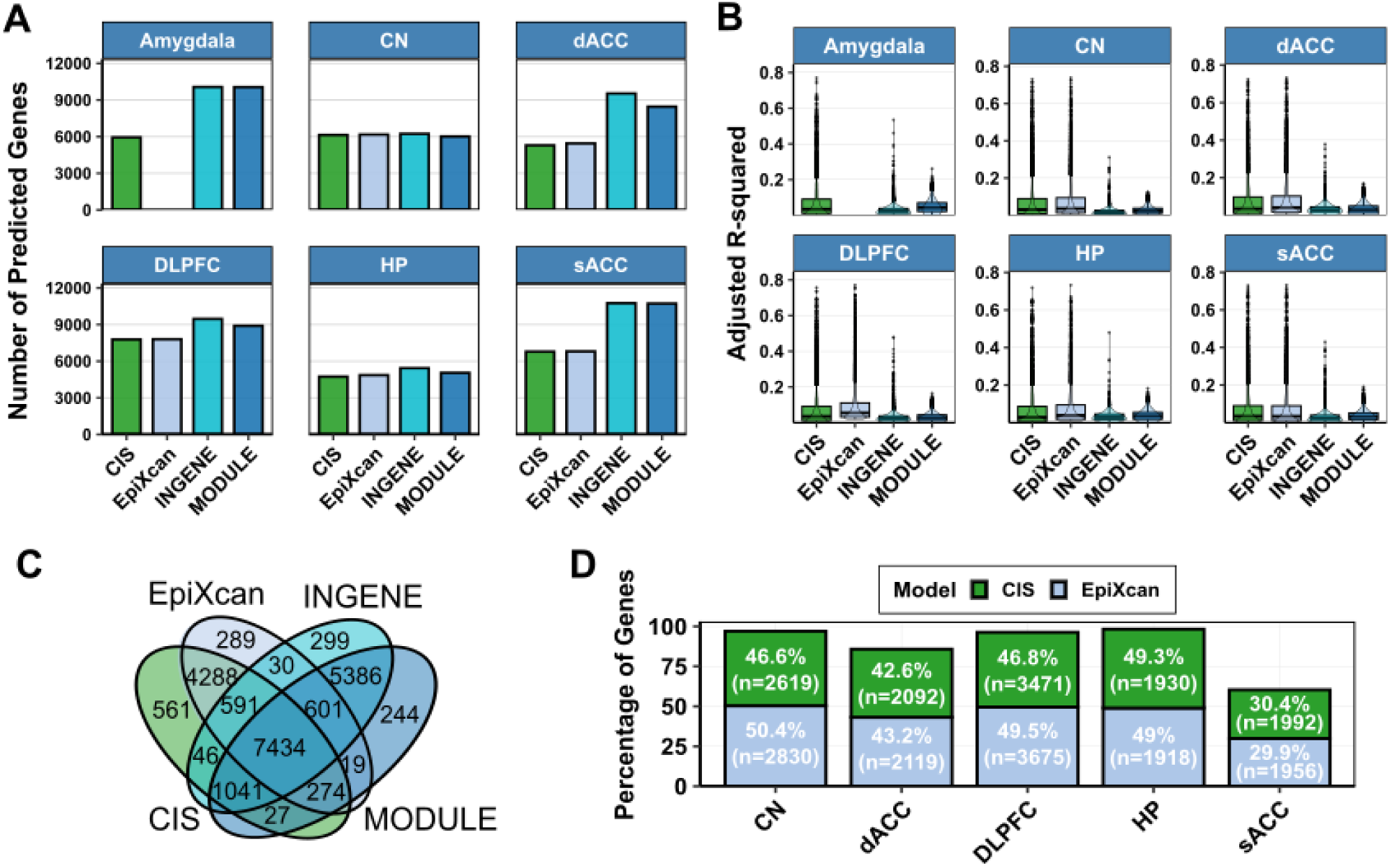
Predictive models replicate across brain regions in GTEx external dataset and predict different genes at different performance. **A)** Barplots showing the number of predicted genes (x axis) in the GTEx dataset by CIS (green), EpiXcan (light blue), INGENE (cyan) and MODULE (dark blue) models. **B)** Box plots of adjusted R^2^ values (x axis) in predicting gene-level expression in GTEx using CIS (green), EpiXcan (light blue), INGENE (cyan) and MODULE (dark blue) predictive models across brain regions. The median is represented by the central line, with the interquartile range (IQR) as the box. Whiskers extend to 1.5×IQR, and outliers are plotted as individual points. **C)** Venn diagram showing the number of total intersecting predicted genes among models. **D)** Bar plots showing the percentage of genes predicted (on the y-axis) within regions commonly identified by CIS (green) and EpiXcan (light blue) models. The percentages indicate the proportion of genes for which one model predicts a higher Adjusted R^2^ value than the other, across a set of “n” commonly analyzed genes.

As expected, due to the generally larger impact of *cis* over *trans* effects on gene expression variance, *cis*-based models explained a higher proportion of variance for the fewer genes they predicted compared to *trans*-informed models. Specifically, the mean adjusted R^2^ values for CIS ranged between 7.1% and 7.8% across regions, while EpiXcan achieved slightly higher values, between 7.2% and 9.4%, aligning with previous findings^14^. In contrast, INGENE explained between 2% and 3.3% of the variance, and MODULE explained between 2.7% and 4.9% (Fig. 3B; the goodness-of-fit for the data is presented in Fig. S9). Across all examined brain regions, Wilcoxon tests comparing adjusted R^2^ distributions between EpiXcan-CIS, EpiXcan-MODULE, and EpiXcan-INGENE consistently demonstrated EpiXcan superior average variance explained (all one-tailed p-values < 2.2e-16). This trend also held for genes commonly predicted by the models, where *cis*- predictions exhibited higher mean R^2^ (Fig. S10). It is also notable how the testing statistics differ from training statistics, underscoring the need to validate the training in independent samples.

In addition to differences in the number of predicted genes and their variance explained, the four frameworks showed both substantial overlap and complementary coverage. The Venn diagram in Fig. 3C illustrates that a large fraction of genes was predicted by multiple models. At the same time, each method contributed unique predictions: CIS identified 561 genes not captured by any other model, EpiXcan 289, INGENE 299, and MODULE 244. These unique subsets underscore the complementarity of *cis*- and *trans*-informed approaches, as well as the added value of retaining all four models within the integrative pipeline.

For genes commonly predicted by CIS and EpiXcan across the four shared tissues, we determined the best-performing *cis*-model based on higher adjusted R^2^ (Fig. 3D). To avoid redundancy and maximize accuracy, all subsequent analyses used the best-performing model between CIS and EpiXcan to represent the *cis*-regulation of gene expression.

To assess the replicability of our *cis-* and *trans*-predictions beyond the GTEx cohort, we extended our analysis by applying our models to postmortem datasets of DLPFC and ACC from the CMC (Table 2). Replicability was quantified both by the correlation of prediction accuracy across cohorts and by the proportion of genes consistently predicted in CMC and GTEx relative to the total number predicted in at least one dataset (Fig. S11). Our findings revealed a consistent pattern of gene predictions across the CMC and GTEx cohorts for commonly imputed genes. Specifically, for *cis*-based models, performance was stable across cohorts: CIS predicted 2,136-4,535 common genes (32-47% of all predicted genes) with cross-cohort correlations of r = 0.43-0.68, and EpiXcan showed similar replicability (2,183-4,565 shared genes, 33-47%; r = 0.43-0.68). By comparison, INGENE yielded 11,057-12,274 shared genes (77-79%) with cross-cohort correlations of r = 0.17-0.52, and MODULE retained 17,594-19,385 shared genes (∼88%) with correlations of r = 0.14-0.17 (Fig. S11). Although absolute correlations for *trans*-based models are lower than for *cis*-only predictors, this pattern is expected given the broader SNP search space and the increased biological complexity of *trans*-regulation. Importantly, the much higher percentage of shared genes highlights the stability achieved by INGENE and MODULE across independent cohorts.

We further benchmarked our LIBD-trained models against BGW^54^ and MOSTWAS^55^ in both CMC and GTEx DLPFC (Supplementary Notes: Section 2.3). Across cohorts, our models predicted the largest number of genes and achieved comparable or superior adjusted R² distributions (Fig. S7C-D). BGW-TWAS and MOSTWAS integrate *cis* and *trans*, explaining their broader performance range, whereas INGENE and MODULE rely exclusively on *trans* signals and therefore showed narrower, more stable distributions. Importantly, the median accuracy of INGENE and MODULE exceeded that of other *trans*-informed models, underscoring the robustness of our co-expression–based approach.

Taken together, these results show that our approach enhances the number of genes exhibiting replicability in independent datasets.

### Functional genetics of co-eQTLs

To further investigate SNPs that exhibit *cis*- and *trans*-regulatory effects among validated predictions, we selected MODULE-derived *trans*-SNPs that were also significant *cis*-eQTLs across 49 tissues in the GTEx QTL collection^56^ (Methods; Table S3). Depending on the brain region, 25-44% of *trans*-SNPs overlapped with GTEx *cis*-eQTLs, and these variants were associated with 5,821 to 19,276 *cis-*regulated eGenes (Table S3). This finding indicates that a substantial portion of the *trans* signals detected by MODULE arises from SNPs that also exert *cis* effects on nearby genes. Gene ontology (GO) enrichment analyses on GTEx *cis*-eGenes genes showed significant overrepresentation of ATP-dependent and catalytic activities, electron-transferase functions, and binding activities of proteins such as MHC class II and cadherin binding at FDR < 0.05 (Fig. S12).

To assess the enrichment of transcription factors (TFs), we conducted a regulomic analysis (Methods). We identified 252 TFs that were significantly overrepresented among the GTEx *cis*-eGenes associated with MODULE *trans*-SNPs (Bonferroni *p* < 0.05; see Fig. S13A). Among all enriched TFs within brain regions, we observed *GATAD2A*, *RERE*, and *IRF3*, all PGC3 prioritized genes along with *SP4*^6^ (Fig. S13B).

### Functional enrichment of SCZ risk variant-associated genes in predictive models

We analyzed genetic associations and functional implications of SNPs regulated by *trans*-eQTL and *cis*-eQTL models in psychiatric disorders, focusing on the potential enrichment for SCZ GWAS-significant SNPs. We hypothesized that SCZ would show relatively high enrichment for *trans*-eQTLs, as evidence supports the role of co-expression networks in channeling genetic risk^24,29,38^.

We correlated log (Odds Ratio) of GWAS-significant^6^ SNPs (*p* < 0.05) with CIS, EpiXcan and MODULE model’s SNP weight, computed as the mean absolute weight across all SNP-regulated genes. We did not perform the same analysis for INGENE as its predictions are based on the EpiXcan and CIS *cis*-regulatory SNPs of co-expressed partner genes rather than direct SNP-to-gene associations, thus the SNPs underlying INGENE reflect partner-gene *cis*-effects rather than the target gene itself. Across all brain regions, both *cis-* and *trans-*based models showed significant positive correlations between QTL weights and the effect sizes of SCZ-associated SNPs (Table S4; Fig. S14), indicating that stronger regulatory SNPs tend to carry higher disease risk. CIS and EpiXcan showed comparable correlations across regions (*r* = 0.14–0.18), whereas MODULE consistently achieved markedly stronger correlations (*r* = 0.28-0.42; p < 0.001) (Table S4). To assess whether these differences were significant, correlations were compared using Fisher’s Z-test (Methods), confirming that MODULE outperformed CIS and EpiXcan (*p* < 2e-16; Fisher’s Z test) across regions (Methods; Fig. S15). These results suggest a stronger association of *trans*- than *cis*-eQTLs with common variant pathogenicity. A similar trend was also found in other disorders, such as bipolar and major depressive disorders, with significant differences in correlation coefficients between predictive models (Fig. S14 middle and bottom panels).

We further tested whether *trans*-predicted genes enriched for SCZ risk SNPs, quantified by the PGC-weight ratio (Supplementary Methods: Section 1.6), exhibited high network connectivity^29^ with PGC3-prioritized genes^6^ across regions. When genes were ranked by PGC-weight ratio and divided into quintiles, mean connectivity increased monotonically across quintiles, as confirmed by linear regression and Spearman’s correlation (Fig. S16; Table S5). Permutation testing (n = 1,000; Supplementary Methods: Section 1.6) established the significance of these trends, revealing robust enrichment for MODULE in DLPFC, CN, HP, and sACC (all p < 0.001), and a weaker effect for CIS in amygdala (p = 0.037). Instead, EpiXcan showed no significant enrichment. This pattern supports the hypothesis that SCZ risk variants are preferentially associated with genes embedded within coordinated co-expression networks with risk genes^29^.

Taken together, these results underscore the biological relevance of interactions among genes predicted by *trans*-eQTLs and the functional role of the SNPs identified, potentially implicating pathways and mechanisms relevant to SCZ etiopathology that act via the mediation of gene co-expression.

### Combination of *cis*/*trans*-scores enhances gene expression prediction in GTEx

We hypothesized that combining *cis* and *trans* predictions should offer the highest number of predictable genes along with the largest explained variance in gene expression. To test this hypothesis, we conducted significance evaluations of gene predictions in GTEx through Maximum Likelihood Estimation (MLE α = 0.05; Methods). We compared a model incorporating covariates and *cis/trans* predictions as explanatory variables against a model with only *cis* and covariates. Whenever the inclusion of *trans* predictions led to a significant increase in adjusted R^2^ compared to the only *cis* model, we considered the gene to be enhanced by *trans*-prediction. We observed a substantial enhancement in the predictability of genes by integrating *cis*, *cis-trans*, and *trans*-genetic predictors within brain regions (Fig. 4A). For *cis*-*trans* predictions, the integration of the *trans*-component consistently yielded an increase in variance explained compared to only-*cis* predictions (one-tailed Wilcoxon p-values < 0.001; Fig. 4B). When consolidating predictions from all brain regions, a total of 18,744 significant genes were predicted by all models (Fig. 4A).

**Figure 4.**
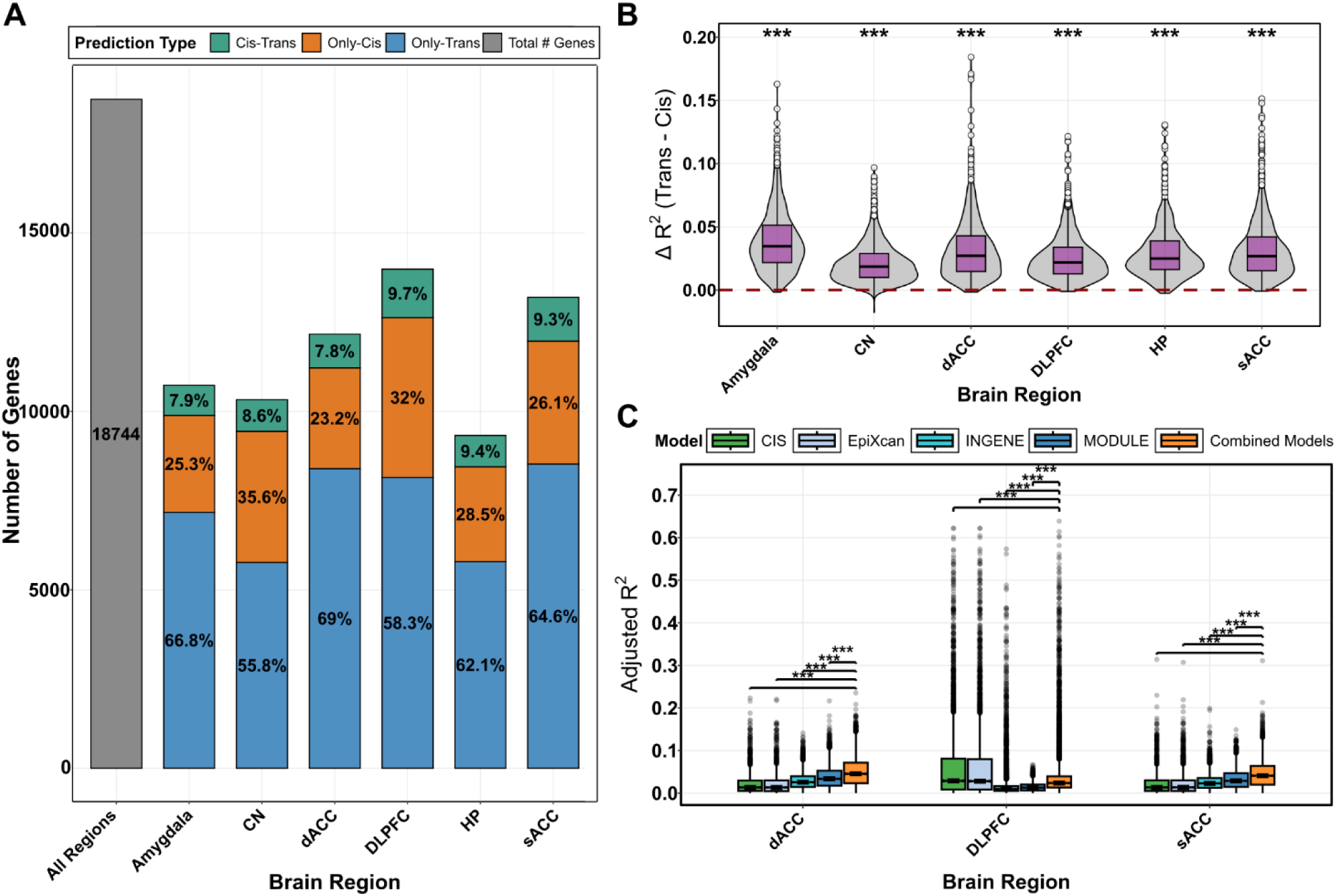
INGENE and MODULE trans-scores improve gene prediction across brain regions. **A)** Barplot showing the number of genes (y axis) significantly predicted across brain regions when combining only-cis (orange), only-trans (light blue) and cis-trans (green) predictions. The grey bar represents unique significant predictions when pooling from all brain regions. B) Distribution of delta adjusted R^2^ values between predictions with at least one trans component and only cis-trans predictors in GTEx data (ΔR^2^(trans-cis)). Asterisks (***) denote a one-tailed Wilcoxon p-value of ≤ 0.001. **C)** Distribution of adjusted R2 values obtained from the combination of cis-trans (orange), CIS (green), EpiXcan (light blue), INGENE (cyan), and MODULE (dark blue) predictions in the dACC, DLPFC, and sACC from the CMC dataset. The median is represented by the central line, with the interquartile range (IQR) as the box. Whiskers extend to 1.5×IQR, and outliers are plotted as individual points. Asterisks (***) denote a Mann–Whitney rank-sum tests p-value of ≤ 0.001.

Having established that the *trans* component enhances the *cis* prediction of gene expression for some genes, we then evaluated whether combining *cis* and *trans* predictions could further improve gene-level prediction. Our goal was to generate a unified prediction model to take advantage of the strengths of each type of prediction. Note that *cis* SNPs were discarded when generating *trans* predictions; therefore, these sets of predictions were independent. For genes with at least two models, we developed a linear model using the GTEx dataset (our testing sample). In this model, true expression values were used as the dependent variable, while a combination of CIS, EpiXcan, INGENE, and MODULE predictors was used as the independent variable. We evaluated the performance of the combined *cis* and *trans* prediction model using the CMC dataset, focusing on the dACC, DLPFC, and sACC regions.

Combining *cis*- and *trans*-based predictions into a single model (*“Combined Models”* in Fig. 4C) improved gene expression prediction across all examined regions, explaining on average 4.1-5.0% of the variance in 7,640-8,600 genes. In comparison, the CIS and EpiXcan models accounted for 2.3–6.4% of the variance in 1,602–3,935 genes, while INGENE and MODULE explained 1.8-2.9% and 1.4-3.7% of the variance across 6,663-8,543 genes, respectively. Mann–Whitney rank-sum tests showed that, in terms of mean adjusted R^2^, the Combined Models significantly outperformed every single-modality predictive model in every region (all p ≤ 0.001; Fig. 4C). We used this combined prediction model for subsequent analyses, including trait association analyses.

Taken together, these findings further highlight the importance of integrating *trans*-effects, along with *cis* scores, to broaden the pool of genes suitable for testing in a gene-mapping setting and provide a more comprehensive prediction of genes associated with complex traits.

### coTWAS identifies trait associations in PGC3 cohorts

To identify genetic associations with SCZ through our integrative approach, we applied *cis-*and *trans*-models to impute gene expression and perform trait mapping using 62 independent genotype cohorts from the PGC3^6^, including 102,613 individuals (Table S6).

Within each cohort and brain region, we selectively retained model-specific genes meeting GTEx selection criteria (see *Evaluation of cis- and trans-models on a testing dataset*; Fig. S17).

We applied logistic regression to associate predicted gene expression with SCZ diagnosis (coTWAS) in each PGC3 cohort separately. We then used the β coefficients from these logistic regressions to perform a meta-analysis of effect sizes across cohorts, with the resulting meta-analytic *p*-values representing the combined evidence of association across cohorts (Methods). To identify independent signals while accounting for LD, we performed conditional analysis following Huckins et al.^15^, across 315 genomic regions based on the distance between the tissue-gene pairs previously meta-analyzed (1Mb window). Genes were added iteratively to joint models based on their meta-analytic *p*-value, and results were FDR-corrected at α = 0.01 to account for non-independence across brain regions due to overlapping subjects (Methods).

Out of 96,535 total tests performed across all brain regions, we identified 1,162 (975 non-MHC) significant associations, corresponding to 766 (693 non-MHC) unique independent genes (Table 1; Table S7). The direction of gene effect size was derived from the adjusted β coefficient of the logistic regression (Methods). Across brain regions, we identified a total of 381 genes that were genetically up-regulated and 414 genes that were down-regulated in SCZ patients compared to controls (Fig. 5).

**Figure 5.**
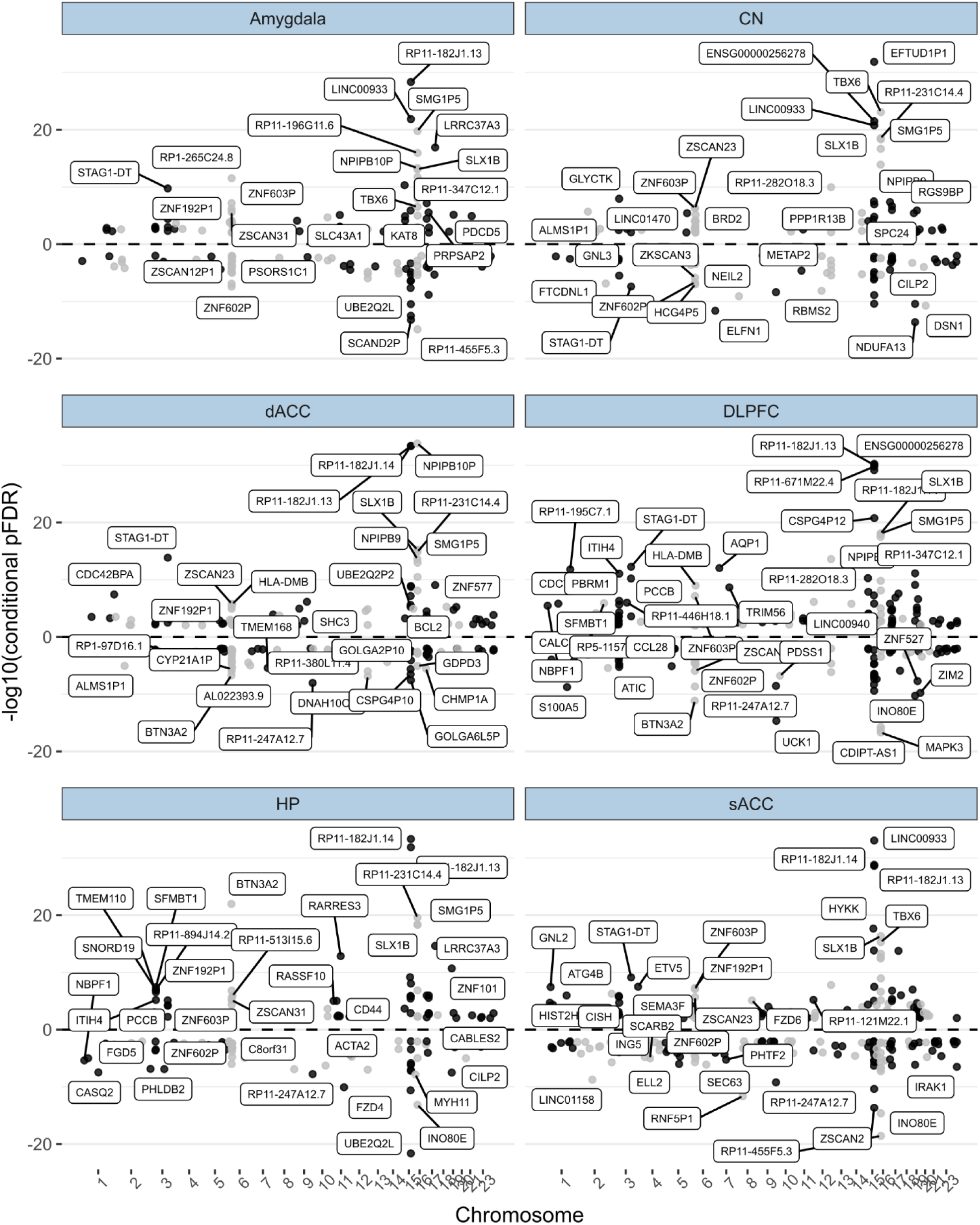
coTWAS significant genes across brain tissues. y axis shows -log10 (conditional p_[FDR]_ < .01), while the x-axis shows chromosomes. Due to clutter, a maximum of 50 gene names per region are displayed. Genes above the dashed line represent positive effects (upregulated in cases), and those below the line represent negative effects (downregulated in cases). Abbreviations: CN: caudate nucleus data; dACC: dorsal anterior cingulate cortex; DLPFC: dorsolateral prefrontal cortex; HP: hippocampus; sACC: subgenual cingulate cortex.

**Table 1.**
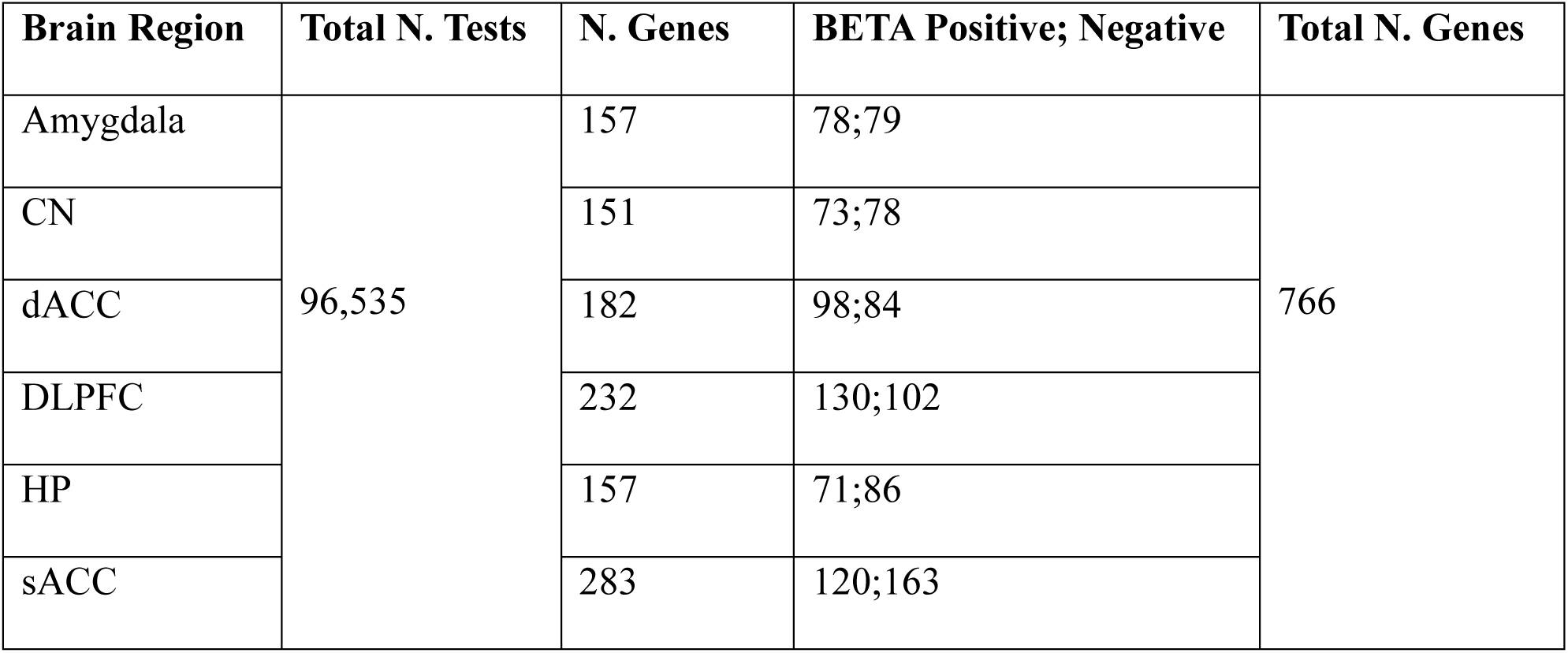
Number of genes tested in the coTWAS framework and number of significant associations at FDR α=0.01 across and within brain regions. Abbreviations: CN: caudate nucleus; DLPFC: dorsolateral prefrontal cortex; dACC: dorsal anterior cingulate cortex; HP: hippocampus; sACC: subgenual anterior cingulate cortex.

| Brain Region | Total N. Tests | N. Genes | BETA Positive; Negative | Total N. Genes |
| --- | --- | --- | --- | --- |
| Amygdala | 96,535 | 157 | 78;79 | 766 |
| CN |  | 151 | 73;78 |  |
| dACC |  | 182 | 98;84 |  |
| DLPFC |  | 232 | 130;102 |  |
| HP |  | 157 | 71;86 |  |
| sACC |  | 283 | 120;163 |  |

To assess the robustness of these associations, we conducted a leave-site-out replicability analysis across the 62 PGC cohorts (Methods). Briefly, for each gene-tissue pair (conditional *p*[FDR] < 0.01), we evaluated the directional concordance and the correlation of effect sizes across held-out sites. Genes exceeding the empirical 95^th^-percentile thresholds for both metrics were classified as replicable. This analysis identified 556 replicable unique genes (empirical one-tailed *p <* 0.05; Fig. S18, Table S7). Across these genes, the mean concordance rate was 69.4% ± 5.1% (SD), and the mean cross-site effect-size correlation was R^2^ = 0.34 ± 0.11. These results indicate that, on average, about 70% of site-specific effect directions were consistent across leave-out iterations, with moderate-to-strong alignment of effect magnitudes. Collectively, these results support the stability and reproducibility of coTWAS-identified gene–trait associations across independent PGC cohorts.

We performed GO enrichment analyses for the 766 coTWAS-significant genes (Fig. 6A) to explore enrichments in SCZ-related biological terms. In the DLPFC, enriched terms considering all β directions converged on Synapse Organization, particularly AMPA glutamate receptor clustering, and on Membrane and Vesicle Transport pathways such as clathrin-coated endocytic vesicle, late endosome, and lysosomal membrane, indicating altered receptor trafficking and turnover. Antigen Processing and MHC complex pathways were significantly enriched and upregulated (β > 0) in both the DLPFC and dACC, and when considering all β directions in sACC and across all regions combined, indicating a broad involvement of neuroimmune signaling in cortical and limbic circuits in individuals with SCZ. These findings delineate a regionally stratified molecular landscape in which excitatory synaptic remodeling predominates in the prefrontal cortex, whereas immune-related biological processes are widespread across regions.

**Figure 6.**
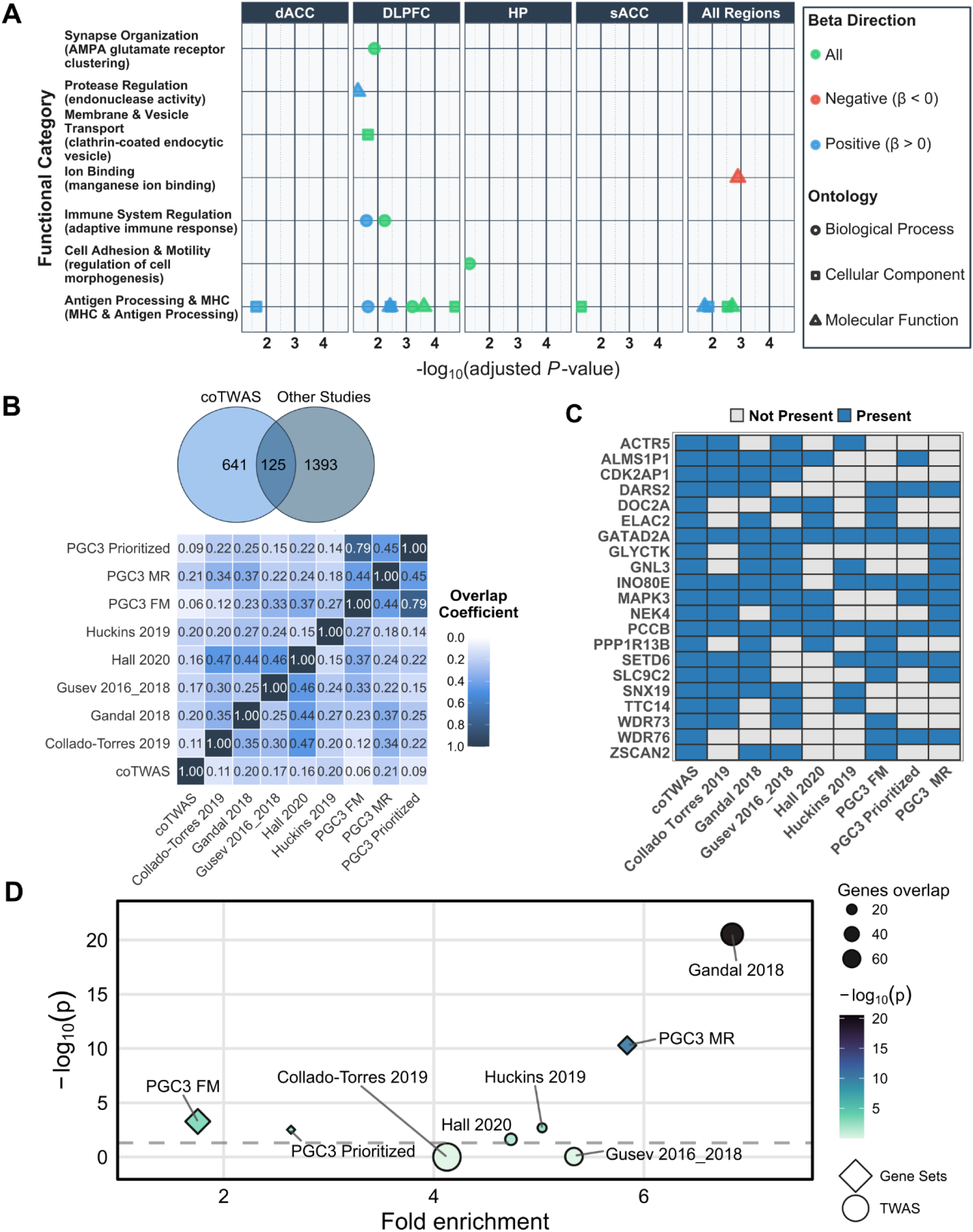
Integrative Analysis of coTWAS-Identified Genes and SCZ-Associated Gene Sets. **A)** Gene ontology enrichment analysis of coTWAS results. The x-axis represents significance as -log10 of adjusted FDR p-values for the top enriched term, while the y-axis shows functional categories grouped by GO domains. **B)** Venn Diagram (top) shows the overlap between coTWAS hits (blue circle) and key SCZ gene sets from previously published studies, including Huckins^15^, SCHEMA^68^, PGC3 Mendelian Randomization, PGC3 fine-mapping, and PGC3 prioritized genes^6^, Boruck^30^, Gandal^69^, Gusev^12^, Hall^70^, and Collado-Torres^71^. Szymkiewicz Simpson coefficient matrix (bottom) quantifies pairwise overlaps among SCZ associations. Darker blues signify greater proportional overlap. **C)** A presence/absence heatmap showing genes on the y-axis and each study on the x-axis. Blue squares indicate that the given gene appears in the referenced study. D) Fold-enrichment (x-axis) versus −log10(p) (y-axis) for the overlap between coTWAS and each indicated gene set, with point size denoting the number of overlapping genes. The dashed horizontal line indicates a p-value threshold of 0.05.

Of the coTWAS significant genes, 354 were predicted by a single model when considering all regions independently, with CIS predicting 21%, EpiXcan 20%, INGENE 40%, and MODULE 19% of these. Across individual regions, the *cis*-component accounted for 26.5-54.3% tissue-specific predictions, *cis*-*trans* combined predictions represented 17.2-30%, and the remaining predictions, ranging from 18.5-44.5%, were attributed solely to *trans*-effects (Fig. S19).

To further understand the link with SCZ risk, we employed the MAGMA Z-score (Supplementary Methods: Section 1.7) to correlate *cis* and *trans*-predictions (Fig. S20). We found a positive correlation for the *cis*-predicted genes (Pearson’s *r* = 0.10, *p* = 0.003). In contrast, the lack of a significant correlation for exclusive *trans*-predictions (Pearson’s *r* = -0.02, p = 0.78) aligns with MAGMA’s underlying design, which focuses on SNPs located proximal to genes. This indicates that MAGMA provides validation for *cis*-predictors, whereas our *trans*-based approaches capture distal regulatory contributions beyond MAGMA’s scope. Supplementary Figure 21 illustrates this complementarity by highlighting top 50 genes with strong coTWAS evidence (log10 adjusted p > 4), and relatively weak MAGMA associations (|Z| < 4).

For coTWAS-significant genes, we further conducted cell specificity enrichment analyses using a human atlas^57^ (Fig. S22). The analyses uncovered a regionally distinct pattern of transcriptional perturbation. In the amygdala, upregulated genes (β>0) were overrepresented in excitatory DG-like neurons (exDG). In the dACC, neuronal stem cells (NSCs) showed enrichment for upregulated genes, while exDG-like neurons were enriched in downregulated genes (β<0). Finally, in the DLPFC, pyramidal neurons (exCA3) and exDG-like neurons displayed gene upregulation, whereas GABAergic interneurons (GABA2) showed downregulation. These results suggest a region- and cell type-specific pattern of alteration in the SCZ, specifically involving neuronal dysfunction.

The coTWAS identified several previously reported SCZ-related genes, including *SNX19*^64^, downregulated across all regions*, GATAD2A*, upregulated in the DLPFC, and *CYP21A1P*^66^ and *C4A*^67^, both downregulated across all regions. All these genes exhibited a predominant *cis* component in their predictions, along with contributions from *trans* models that varied depending on the brain region. To further validate our findings, we cross-referenced our coTWAS significant genes with previously identified SCZ-associated gene-sets. Specifically, we compared our results with SCHEMA genes^69^, the 120 genes prioritized by the PGC3 study^6^, as well as those identified through Summary-based Mendelian Randomization (MR) and Fine-Mapping (FM)^6^ (Fig. 6B; Fig. S23-25). Additionally, we included genes identified by Borcuk et al.^29^ as network neighbors of SCZ risk genes, and previous SCZ TWAS^12,15,70–72^ (Fig. 6B).

Our analysis replicated many well-established SCZ associations (Fig. S26), validating these predictions, while also identifying novel candidates (Fig. 6B, Venn diagram). Notably, coTWAS is the only method, aside from PGC approaches, to show a high overlap Szymkiewicz Simpson coefficient with MR genes^6^ (Figure 6B bottom). While this overlap does not validate the *trans*-based associations, it provides complementary evidence that coTWAS signals are consistent with large-scale causal inference frameworks based on *cis*-eQTL instruments.

The coTWAS set included 125 genes previously implicated in other studies, alongside 641 uniquely identified genes, corresponding to an 83.7% novel discovery rate (Fig. S27A). When stratifying associations by prediction type (*cis*-only, *cis* + *trans*, *trans*-only; Fig. S27B), most previously reported genes were recovered by *cis* and *cis* + *trans* models (28.8-56% of known genes). In contrast, novel associations (51%) were strongly enriched among *trans*-only predictions, indicating that distal regulatory effects are the predominant source of novel gene-level discoveries in our framework. These results emphasize the strength of *trans*-eQTL-driven modeling in capturing regulatory effects missed by traditional *cis*-only methods.

Among the overlapping genes, 21 genes were consistently detected in at least four independent studies (Figure 6C). Although this set of overlapping genes was not significantly enriched for any gene ontology, key risk loci emerged across independent studies, including *MAPK3* and *INO80E*, both residing within the 16p11.2 neurodevelopmental risk locus^73^, as well as *GATAD2A*, a component of the NuRD chromatin-remodeling complex, and *GNL3*, which regulates neural progenitor maintenance^74^.

Enrichment analysis (Fig. 6D) revealed significant overlap with the Gandal gene set^70^ (p = 2.98 × 10⁻²¹) and, to a lesser extent, with the Huckins^15^ (p = 0.002) and Hall^71^ (p = 0.023) gene sets, highlighting a significant overrepresentation of recurrently identified genes across independent studies. To further assess overlap with known SCZ loci, we mapped coTWAS-significant genes to genome-wide significant PGC3 regions^6^. Overall, 158 genes (27.4% of all tissue-specific associations) were within PGC3 loci, while 608 (72.6%) lay outside (Fig. S28A). *Cis* and *cis*–*trans* predictions accounted for most within-locus associations, whereas *trans*-only signals predominantly arose beyond known intervals (Fig. S28A). This pattern was consistent across regions (Fig. S28B), with *trans*-only predictions averaging 2.0% within PGC3 loci versus 28.2% outside. The chromosomal distribution (Fig. S28C) revealed that several of the most significant associations (- log₁₀ FDR > 20) clustered outside established GWAS peaks, highlighting that coTWAS uncovers distal regulatory effects extending beyond known SCZ loci.

In conclusion, our coTWAS analysis identified 766 significant gene associations with SCZ, of which 641 represent novel TWAS associations across all examined brain regions. These results underscore the utility of integrating gene co-expression data implicating gene interactions into the TWAS framework.

## Discussion

In this study, we introduced gene expression prediction models for six brain regions (amygdala, CN, dACC, DLPFC, HP, and sACC) leveraging data from the LIBD postmortem repository. We developed predictive models based on co-expression to identify and assess *trans*-eQTLs and established a pipeline to integrate *trans* and *cis*-scores. We demonstrated that *cis*/*trans* score integration enhances prediction accuracy at the gene level. Given that predictive performance correlates positively with the power to detect trait associations^75^, we applied *cis* and *trans* predictive models to PGC3 genotypes to obtain a gene-level TWAS^6^. We found significant SCZ associations for 766 genes across brain regions.

### *Cis*- *trans*-scores integration

We implemented a cross-dataset training and evaluation strategy to stabilize *trans*-predictive models and reduce overfitting. This cross-dataset framework prioritized genes showing consistent *trans*-regulatory effects across independent resources while maintaining or expanding the pool of predictable genes relative to classical *cis* approaches (CIS, EpiXcan). When applying *cis*- and *trans*-predictive models in the GTEx-independent cohort, we observed an increase in the number of predicted genes compared to using *cis*-models alone (Fig. 3A).

While our *trans*-models successfully predicted a greater number of genes, the performance metrics favored *cis-*models in terms of prediction effect within the testing dataset (Fig. 3B; Fig. S10). This finding is expected, given that the effect size of *cis-*variants is larger compared with *trans-*eQTLs. However, EpiXcan and CIS effectively predicted about ∼15,000 genes, compared with ∼16,000 of *trans*-models (Fig. 3C). The increase in predictable genes associated with *trans*-variants, which replicates in two independent datasets (GTEx and CMC, Fig. S11), illustrates that distal regulatory variants, although individually weaker, contribute cumulative predictive information that complements *cis* regulation and enhances the quantitative fidelity of genetically imputed expression in the human brain.

Accordingly, in GTEx predictions, the integration of *cis*- and *trans*-scores significantly en-hanced gene prediction across brain regions (Fig. 4A-B). Notably, approximately half of the genes were significantly predicted by solely *trans* models, confirming that the synergistic integration of *cis* and *trans* scores significantly boosts discovery power. Identifying specific genes related to complex diseases is challenging because many susceptibility genes interact and contribute to more than one specific pathway, ultimately causing complex biological dysfunction^30,32,39,76^.

### Biological mechanisms of *trans*-predictive models

The architecture of *trans* prediction highlights how distal effects are embedded in functional networks. INGENE models *cis*-mediated *trans*-effects^27^, while MODULE captures shared variance in the expression of multiple genes^30,38^, two complementary mechanisms that together capture the hierarchical organization of transcriptional control. It is indeed conceivable that both *trans*-models might be capturing similar information. We found that up to 29% of MODULE-predictable genes have at least one *cis*-SNP regulating one of the co-expression partners acting as predictors in INGENE (Table S1). This means that a proportion of the genes predictable via MODULE are associated with *cis*-mediation effects in genes co-expressed with the target gene.

We also found that approximately 40% of MODULE *trans*-eQTLs were significant *cis*-eQTLs for at least one eGene across 49 tissues, based on data from the GTEx QTL resource (Methods; Table S3). Interestingly, we found overrepresentation in TFs (Fig. S13A). This finding suggests a poten-tially important regulatory role for TFs in mediating the effects of *trans*-eQTLs on gene expres-sion^77,78^. Among these genes, several have been previously associated with biochemical pathways relevant to SCZ^6^. Notably, *GATAD2A*,^5^ is involved in gene silencing, *RERE*^69^ acts as a transcriptional repressor during development, *SP4* is linked to both SCZ^79^ and bipolar disorder^80^, and *IRF3* is known to play a key role in immune response and inflammatory processes^81^. Additionally, *ESRRA* has been identified as a regulator of *DRD2* co-expression networks associated with schizophrenia risk^43^, while the dysregulation of *HDAC1* has been widely documented in the context of the disorder^82,83^. These observations suggest that these SCZ-associated genes, regulated by *cis*-eQTLs with *trans*-eQTL ef-fects, contribute to the disease via regulatory networks.

It is important to note, however, that most MODULE-predicted gene regulation did not yield evidence of mediation by *cis*-indirect effects. Among multiple non-exclusive processes potentially underlying the remaining 70% of gene expression predictions are the influence of SNPs in *cis* to master regulators that is not mediated by transcription, and the potential failure to identify *cis*-eQTLs due to brain region, cell type, or age stage specificity of *cis*-effects, and feedback regulation from expressed genes^17,84–86^. Inevitable false positives may leave a portion of gene expression unexplained by known processes. Perhaps the simplest explanation of how MODULE predictions originate is that the RNAseq quantification of each single gene at the time of death is subject to uncertainty, and the simultaneous expression of other genes typically co-expressed with the target gene reduces such un-certainty. In other words, co-expression data enhances the detection of functional eQTLs and thus contributes to reducing false negatives.

### Co-expression based model advantage

Gene-level association methods^13,14^ or isoform-level TWAS^87^ represent a notable advantage by pinpointing genes within loci that fall short of genome-wide significance in GWASs. These studies identified SNPs often exhibiting GWAS p-values in the range between 5×10^−8^ and 10^−3^, suggesting borderline associations that could attain significance in larger studies. Our approach deviates from traditional methods by employing co-expression-based models, which re-annotate genetic variants to local (*cis*) and distal (*trans*) genes based on statistical association and not gene proximity. This approach captures the collective impact of variants within functionally relevant networks, moving beyond individual SNP analysis. In our study, *trans*-eQTLs identified by MODULE showed a stronger positive correlation between SNP gene-expression prediction weights and the log(OR) from PGC3 SCZ summary statistics compared to *cis*-eQTLs (Figs. S14-S15), indicating a stronger association with risk.

### Diagnosis association

Applying the integrated *cis*–*trans* framework to 102,613 cases and controls from the PGC3 collection (Table S6) enabled the identification of gene-trait associations reflecting contributions from both regulatory layers. A large proportion of these associations were reproducible in cross-validation (Table S7; Fig. S18), indicating stable performance of the framework across independent cohorts within the PGC3 collection.

The robust positive correlation between the coTWAS association strength and the MAGMA Z-score for *cis*-derived predictions indicates direct and localized control by *cis*-regulatory elements near the genes they regulate. Conversely, the lack of significant correlation for exclusive *trans*-predictions (Fig. S20) highlights the additional insight available when using *trans*-eQTLs. In this scenario, integrating both *cis* and *trans* models provides a more comprehensive understanding of genetic influences on SCZ.

GO enrichment (Fig. 6A) and cell specificity analyses (Fig. S22) of coTWAS genes delineate two convergent axes of SCZ pathophysiology: dysregulation of AMPA receptor trafficking within excitatory neurons and immune-endosomal activation processes. Consistently with recent large-scale studies, our findings support the view that SCZ arises from both neuronal and non-neuronal perturbations across multiple brain regions, with associations varying by both region and cell state^88^. Large-scale single-cell transcriptomic findings have shown that excitatory neurons exhibit pronounced disease-associated expression changes^70^, paralleling our results. In the DLPFC, vesicular and endocytic genes enriched in excitatory neurons were associated with AMPAR trafficking and receptor internalization dynamics, consistent with previous reports of altered receptor trafficking in SCZ^89,90^. Genes enriched in GABAergic interneurons showed relative downregulation in the same region, in line with prior evidence of excitation–inhibition imbalance in SCZ^91,92^. The enrichment of antigen-processing and MHC pathways across prefrontal and cingulate regions is consistent with evidence that neuroinflammatory activity exacerbates inhibitory neuron down-regulation in SCZ^93^. Although classically immune-related^67^, MHC class I molecules also modulate NMDA and AMPA receptor trafficking and regulate synapse density during development^94^, suggesting that the glia component may not involve neuroimmune biology. Although certain genes are preferentially expressed in certain cell types, the eQTL effects we measure may also depend on their expression in other cell types, hence entangling attempts to parse out cell specificity purely based on computational approaches.

The intersection of coTWAS hits with previous SCZ studies^6,12,15,29,59,69–72^ (Fig. 6B-D; Fig. S26) revealed both validation and novelty. While 125 of these genes were already reported, over 80% were newly identified (Fig. 6B; Fig. S27A), with *trans*-only models accounting for most of these discoveries (Fig. S27B). The significant overlap between genes identified through coTWAS analysis and Mendelian Randomization (MR)^6^ (Fig. 6B bottom) indicates that a subset of our coTWAS associations are also supported by large-scale causal inference analyses. We note, however, that the PGC3 MR study relies exclusively on genome-wide significant *cis*-eQTLs as instrumental variables, and therefore cannot be used to directly validate the *trans*-regulated signals identified by coTWAS. Although TWAS and MR are conceptually related, they rely on distinct statistical frameworks and assumptions, and the convergence of results with this external large-scale analysis is reassuring that our coTWAS prioritizes biologically meaningful associations in SCZ. Several coTWAS genes replicated across at least four independent SCZ studies (Fig. 6C), converging on synaptic signaling, chromatin remodeling, and neurodevelopmental control. The MAPK3 cascade, regulated by glutamate receptor activity^95^, links excitatory transmission to transcriptional programs for synaptic plasticity. Consistently, INO80E, GATAD2A, and GNL3, reportedly altered in postmortem SCZ cortex^73,74^, implicate the coordinated disruption of transcriptional regulation and synaptic function.

### Limitations

Our study is limited by the relatively small post-mortem brain sample size for the purpose of genetic associations. Training sample size is pivotal to developing prediction models^13,96^. However, our design involved the use of additional testing datasets to estimate the generalizability of our conclusions. Second, our study is based on bulk tissue data, hence not capturing cell-specific effects. Moving from bulk data to a more refined cell-specific approach may substantially improve the ability to identify specific and *trans*-eQTL regulatory pathways^41,97,98^, but current cell-specific datasets are limited in scale. Third, our study does not account for the influence of sex on gene expression. Sex is known to contribute to the heterogeneity observed in complex diseases like SCZ^99,100^. However, accounting for sex would have halved the samples available to develop sex-specific predictors, ultimately decreasing the signal-to-noise ratio. Fourth, we used European ancestry genotypes because most subjects in PGC3 have this ancestry. Lastly, concerning our SCZ association analysis, relying on genotype imputation may influence the detection of significant associations and the downstream functional analyses.

## Conclusion

Collectively, our findings highlight the effectiveness of integrating co-expression as prior information in predictive models and underscore the importance of combining *cis*- and *trans*-scores to study disorders, like SCZ, showing potential *trans*-heritability. This approach yields gene expression predictions that are independent of GWAS statistics, thus complementing traditional polygenic risk scores and providing a comprehensive tool for understanding individual genetic backgrounds. Furthermore, our study underscores the value of integrating *cis* and *trans* prediction models for identifying genetic factors associated with SCZ. The novel associations discovered reveal the direction of genetic effects associated with SCZ for each gene with brain region granularity, hence informing physiological mechanisms of illness. In conclusion, our findings suggest that gene-gene interactions in biological networks are key to understanding genetic risk for SCZ.

## Methods

### Subjects

The training dataset for this study consisted of genotype and brain tissue obtained from individuals of European ancestry (see the Genotype post-imputation processing section). The samples included neurotypical controls (NC), individuals with schizophrenia (SCZ), bipolar disorder (BP), and major depressive disorder (MDD). These samples were sourced from the Lieber Institute for Brain Development (LIBD) collection. Characteristics of individuals are reported in Table 2.

**Table 2.**
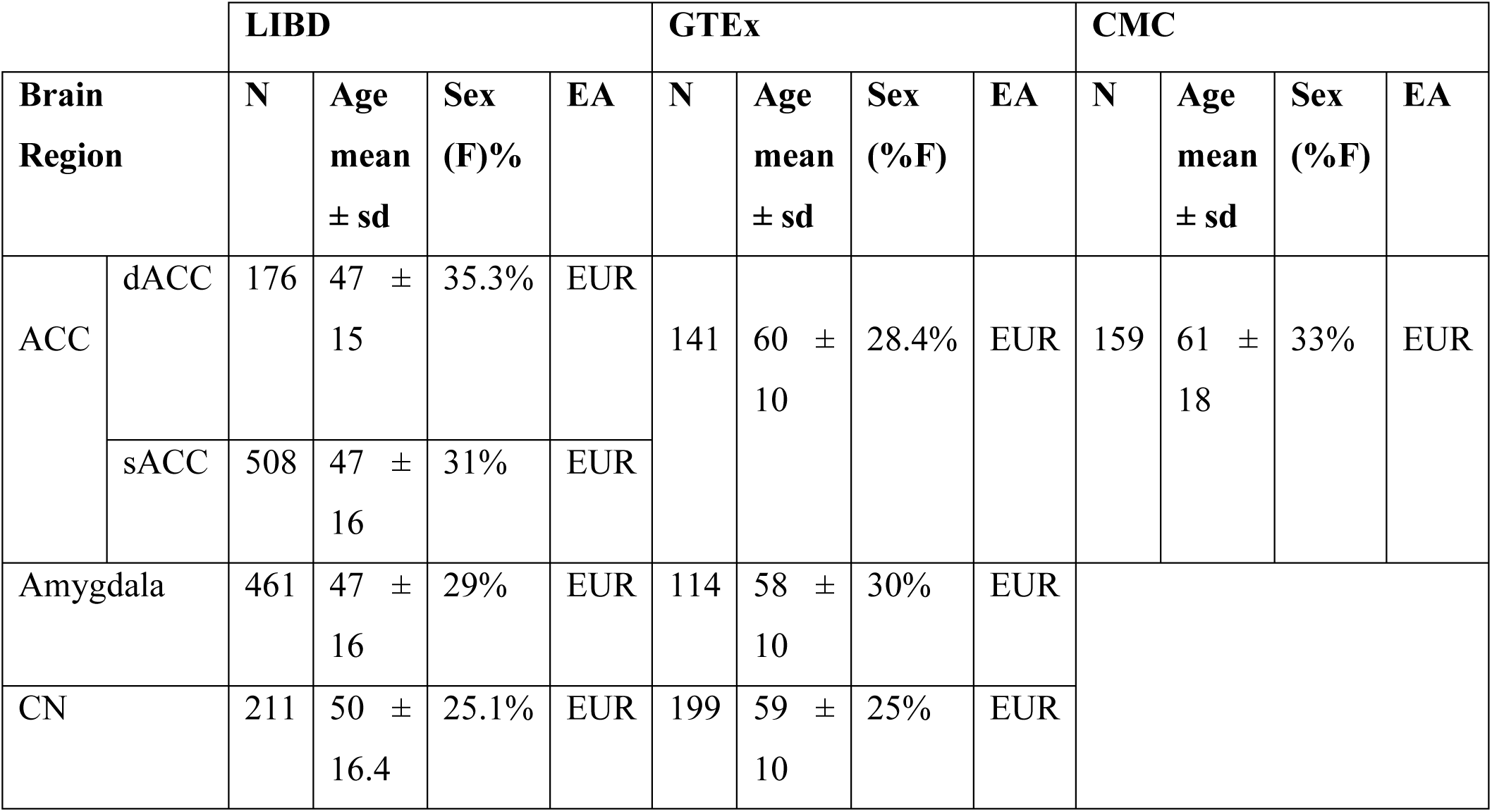
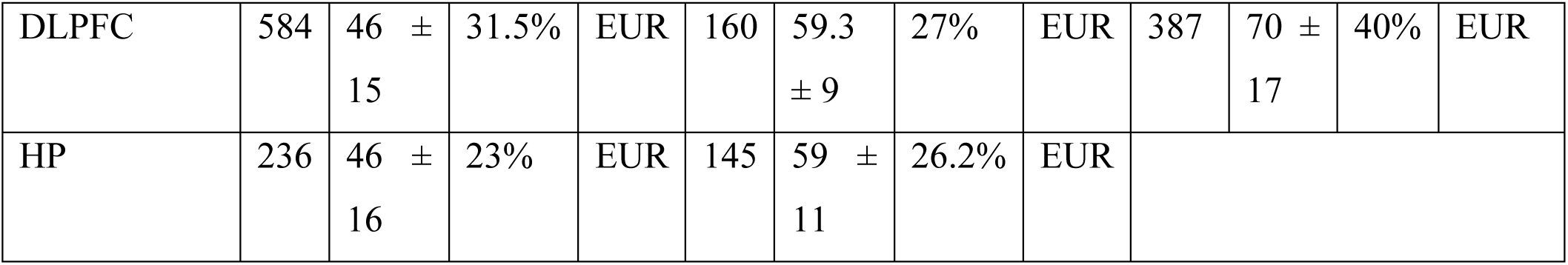
Postmortem data demographics. Abbreviations: N: sample size; F: female; EUR: European; dACC: dorsal anterior cingulate cortex; sACC: subgenual anterior cingulate cortex; CN: caudate nucleus bulk tissue data; DLPFC: dorsolateral prefrontal cortex bulk tissue data; HP: hippocampus bulk tissue data.

The replication studies were carried out using the (i) Genotype-Tissue Expression (GTEx) post-mortem data; here, we selected European Ancestry NC individuals ranging in age from 20 to 79 years (Table 2); (ii) CommonMind Consortium (CMC) post-mortem data from donors with SCZ, bipolar disorder, and NC. We selected individuals of European Ancestry and with ages ranging between 17 and 90 years (Table 2); (iii) PGC3 genotype samples from 102,613 individuals (Table S6) of European Ancestry (SCZ = 42%).

## Materials

### Selection criteria

The discovery dataset of this study included post-mortem human brain specimens from both healthy individuals and individuals diagnosed with SCZ, BP, or MDD. To maintain consistency with our primary post-mortem replication dataset (GTEx) and enhance the statistical power of our predictive post-mortem pipeline, we included only individuals of European ancestry aged 17 years or older. This approach also ensured robust validation in post-mortem samples before applying the predictions to PGC3 cohorts in the SCZ diagnosis association study. Ethical approvals for each dataset used in this study are detailed in Supplementary Methods Section 1.1.

### Lieber Institute for Brain Development (LIBD) post-mortem RNA-seq data

This study used post-mortem gene expression data from tissue homogenate RNA-sequencing. Brain samples were obtained from various sources, each with specific collection and processing protocols. Details of each of the samples are given in the Supplementary Methods (Section1.2). In our analysis, we included samples from the dorsolateral prefrontal cortex (DLPFC)^72,101,102^, hippocampus (HP)^72,103^, caudate nucleus (CN)^104^, amygdala^102,105^, dorsal anterior cingulate cortex (dACC)^102^, and subgenual anterior cingulate cortex (sACC)^105^. Genotype–RNA-seq concordance checks were performed to verify sample identity following the SPEAQeasy pipeline (https://research.libd.org/SPEAQeasy-exam-ple/swap_speaqeasy.htm).

### Post-mortem RNA-seq data – replication studies

For GTEx, we used publicly available and qual-ity-controlled gene expression data from the GTEx consortium v8 (dbGaP Accession phs000424.v8.p2) (http://www.gtexportal.org). The data included samples from the ACC, amygdala, CN, DLPFC (Frontal Cortex BA9), and HP.

We downloaded the CMC study RNA-seq data from the CommonMind Consortium Knowledge Portal (https://doi.org/10.7303/syn2759792). We included samples from DLPFC (syn18097439, release 3.0) and ACC (syn29442240, version 6.0).

### Genotype data

For LIBD brain collections, genotype data were processed and imputed as previously described^101,106^. Genotyping involved Illumina BeadChips, with subsequent imputation through the Trans-Omics for Precision Medicine (TOPMed) imputation server^107^ and the Haplotype Reference Consortium^108^ (HRC) reference panels. GTEx data followed the GTEx pilot analysis^109^, with imputation through IMPUTE2^110^ and the 1000 Genomes Phase 1 freeze as the reference panel^111^.

CMC genotyping was performed using DNA isolated from the DLPFC using the Illumina Infinium HumanOmniExpressExome platform and imputed to HRC.

### Data preprocessing

All analyses have been performed primarily using the R statistical software (version 4.0.0+).

#### Genotype post-imputation processing

Genotype datasets (LIBD, GTEx and CMC) were uniformly processed for post imputation quality control (QC) using the PLINK toolkit version 1.07106. This procedure involved the removal of variants with duplicates, deviations from Hardy-Weinberg equilibrium (HWE) with P < 10^−6^, minor allele frequency (MAF) < 0.01, or missingness rate exceeding 0.05. Additionally, we excluded subjects in the dataset whose information was incomplete for more than 2% of SNPs. The degree of recent shared ancestry, i.e., the identity by descendent (IBD), was estimated within the cohorts to define the relatedness of all pairs on individuals through the PLINK function ‘--genome’. We considered 0.125 to represent the relatedness of 3rd degree that was used as a cut-off to exclude the possible influence of relatedness within cohorts on the dependency between observations. To control for population stratification, we performed a cross-validation algorithm for a lasso and elastic-net regularized generalized linear model using HapMap3107 as a reference panel. We calculated eigenvalues and superpopulations were assigned based on the overlap of principal components between the sample and reference. Individuals with less than 90% genetic overlap for European ancestry were excluded108. The final number of genotypes after processing was as follows: LIBD = 7,521,829; GTEx =8,623,182; CMC = 5,859,752. Additionally, we subset LIBD genotypes with overlapping SNPs in the GTEx cohort to maximize the power of further analyses, resulting in a final number of 6,819,569 SNPs.

### PGC3 cohorts genotype preprocessing

We analyzed 62 cohorts of European ancestry for which genotype information was available from the PGC3 (Table S6). Genotype data acquisition, imputation, and processing as well as calculation of genomic eigenvariates (GEs) for population stratification were performed in each cohort separately as previously reported by the PGC.

We selected for each cohort separately SNPs with high imputation quality (INFO >0.9), low missingness (<1%), minor allele frequency > 0.01 and in Hardy-Weinberg equilibrium (HWE: P > 10^-6^) using the PLINK v1.09 software.

### Analysis Pipeline

#### Quantification and Statistical Analysis

We used the “recount3” R package^112^ to quantify gene-level mRNA expression by converting processed gene counts and gene lengths into reads per kilobase per million mapped reads (RPKMs). We selected genes above median RPKM ≥ 0.1 and free of floor effects (maximum 20% of zeroes per gene). We log-transformed RPKM with an offset of 1, i.e., log2(RPKM+1), and we used inter-array distance to identify outlier subjects deviating more than 3 SDs from the mean ^38^. After removing mitochondrial genes, the final datasets had varying numbers of genes for different regions (see Table S8). We next converted LIBD and GTEx RPKM values into transcripts per kilobase million (TPM).

In the LIBD dataset, we conducted separate regression analyses for each brain region, accounting for various confounding variables. These included diagnosis, sex, age, RNA Integrity Number (RIN), mitochondrial mapping rate, ribosomal RNA (rRNA) rate, gene mapping rate, ancestry estimated through the first five ancestry components (see the Genotype post-imputation processing section), and the first three gene expression principal components (PCs). We found a significant correlation between individual neuronal proportion (”neu”), calculated using the R package BRETIGEA^32^, and the principal components PC1 to PC3. The Pearson’s correlation coefficients for each brain region were: 0.85 (Amygdala), 0.64 (CN), 0.76 (dACC), 0.70 (DLPFC), 0.90 (sACC), and 0.83 (HP), all with p-values < 0.05. As a result, we decided not to include “neu” as a regressor to avoid collinearity issues. The PCs already account for this factor, having been used as surrogate variables to clean the expression datasets, making the additional inclusion of “neu” unnecessary.

Similarly, in the GTEx dataset, log-transformed RPKM was residualized by sex, mean age, RIN, rRNA, postmortem interval (PMI), the first five ancestry components, and the first three principal components (PCs). Here, we identified a correlation between “neu” and the PCs with absolute Pearson’s correlation coefficients of 0.87 (Amygdala), 0.92 (ACC), 0.84 (CN), 0.91 (DLPFC), and 0.73 (HP), leading us to exclude this association from consideration.

The CMC log-transformed RPKM data was residualized by diagnosis, sex, age, postmortem interval, RIN, rRNA_Rate, the ratio of exon-mapped reads to the total reads sequenced, intronic and intergenic rate, the first five ancestry components, and the first three PCs (Pearson’s *r*: 0.75 - ACC and 0.62 - DLPFC).

For all gene-expression datasets, we next rank-normalized residuals using the Blom formula^20^ to limit the impact of deviation from normality in expression data for subsequent analysis. Table S8 shows the number of genes surviving after preprocessing.

#### cis-eQTL discovery

*Cis*-eQTLs were defined as SNPs located within ±1 Mb of each gene’s TSS. For each gene, the TSS was taken as the 5′ end of the annotated gene: the start coordinate for genes on the positive strand and the end coordinate for genes on the negative strand. Associations with gene expression were tested in a 4-fold cross-validation framework using robust linear models (*robustbase* R package). Statistical significance was assessed with F-tests from the *sfsmisc* package. Results across folds were combined using fixed-effect inverse variance meta-analysis, implemented in the *stats* package, yielding a single meta-analyzed effect size, standard error, and p-value for each SNP–gene pair. SNPs were then ranked by p-value, and LD pruning was performed by calculating pairwise *R²* within ±250 kb of the top-ranked SNP. Independent signals were defined as *R²* < 0.1, and lower-ranked SNPs were iteratively removed to retain the strongest associations^38^.

#### Training and Application of *cis*-eQTLs based model: CIS

To develop our CIS training model, we employed the approach originally established by the PrediXcan^13^ method. We used matched genotype and gene expression data to identify a specific set of genetic variants within ±1 Mb of each gene, which significantly impacts gene expression within brain regions. The influence of these variants was quantified using regression analysis, implemented within a 4-fold CV framework.

To assess the accuracy of our predictions, we compared the predicted gene expressions with actual measurements across CV folds. Only models exhibiting adjusted R^2^ ≥ 0.01 (*p* < 0.05), were selected for inclusion in our final predictor database.

We applied the tissue-specific CIS-models to the testing genotypes using the MetaXcan^113^ “Predict.py” function (https://github.com/hakyimlab/MetaXcan).

To extend the *cis*-based framework with functional priors, we trained EpiXcan^14^ which incorporates epigenomic annotations into the elastic-net penalty scheme. We retrieved annotations from the Roadmap Epigenomics^53^ portal (https://egg2.wustl.edu/roadmap/web_portal/) and trained tissue-specific models following the publicly available source code in the EpiXcan Bitbucket repository (https://bitbucket.org/roussoslab/epixcan/src/master/). Whenever methodological details were explicitly specified by the authors, we applied the same parameters to maximize comparability. Unspecified parameters and procedures were aligned with our CIS pipeline to ensure internal consistency in our work.

### Training and application of co-expression-based predictive models: INGENE and MODULE Co-expression networks

We leveraged previously published networks derived from Weighted Correlation Network Analysis (WGCNA)^114^ across various studies^32,33,35–38,70,86,115,116^, obtaining 48 co-expression networks. The gray module within each network was excluded from subsequent analyses as it is comprised of non-clustered genes.

### INGENE

#### Models and assumptions

The INGENE model predicts gene-level expression from genetically *cis*-predicted co-expression partners. The model assumes gene G with P co-expression partners and uses an additive genetic model to estimate the gene expression trait Yg:

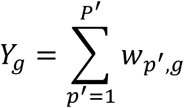

where Yg is the expression trait of gene g, and wp’,g is the effect size of the p’ partners of gene g estimated.

#### INGENE training and application to testing dataset

We first generated CIS and EpiXcan-derived predictions in the LIBD training genotype. Both *cis* models were used for INGENE training: they provided the initial predictions for co-expression partners, which serve as gene-level predictors in INGENE. We applied linear regression to compute gene-level prediction performance metrics for each model. To ensure the most accurate and robust predictions within the INGENE framework, we selected for each gene the best-performing model. Specifically, when both EpiXcan and CIS models predicted the same co-expression partner, we chose the predictor from the model that exhibited superior performance based on a strictly positive adjusted R², thus optimizing the predictive accuracy of the INGENE-imputed co-expression matrices.

Next, we adapted the Elastic-Net linear regression pipeline implemented in PrediXcan^13^ to train the model for each brain region and for each module in the 48 co-expression networks. We fine-tuned the Elastic-Net penalty parameter lambda using nested 4-fold cross-validation (CV) through the “cv.glmnet”^116^ function and weigh co-expressed predictors. We extracted non-zero predictor weights and summary statistics to generate a prediction database. We retained only significant models with adjusted R^2^ ≥ 0.01, p-value < 0.05.

To apply INGENE to the testing data, we first imputed *cis*-mediated predictions using CIS and EpiXcan. These predictions were used to infer co-expression partners, to which predictive weights were applied. For genes predicted by multiple networks, we averaged predictions and selected the *cis*-derived estimate with the highest adjusted R² between predicted and observed expression for inclusion in the final INGENE output.

### MODULE

#### Model and assumptions

In the MODULE model, gene-level expression is predicted based on candidate *trans*-eQTLs identified using the co-expression matrix module eigengene (ME). For each module within the respective network and in each brain region independently, we computed the ME, representing the first principal component (PC1) of the brain region-specific co-expression matrix. The MODULE model assumes gene G with P co-expression partners and a set of markers K associated with the ME. The *trans* component of gene expression under this model is given by:

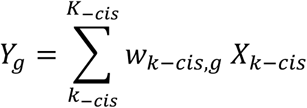

where Yg is the expression trait of gene g, K-cis is the set of markers removing the *cis*-SNPs (±1 Mb) around the gene, w_k-cis,g_ is the effect size of the marker, and X_k-cis_ is the number of reference alleles for marker K.

#### co-eQTL discovery

For the identification of markers associated with ME of the co-expression matrix, we first used MAGMA (v1.09b)^117^ to map European 1000G SNPs^117^ within ± 100 kbp of each gene in the respective co-expression module. We next conducted a comprehensive analysis of co-expression quantitative trait loci (co-eQTLs) in a cross-validation framework. To reduce false positive findings, we partitioned the dataset into 4-folds, stratified according to the diagnosis groups. In each iterative step, we subset genotype and gene-expression data and computed the RNA-seq PC1 for the specific training set. We fit a robust linear model for each SNP with genotype as the explanatory variable and PC1 as the dependent variable. Subsequently, we conducted an F-test to assess the significance of SNP associations, extracting p-values using the “*f.robftest*” function from the R “*sfsmisc*” package. We next ranked SNPs in ascending order based on their p-values and reassigned new ranks for common associations across folds by computing the rank product. We calculated pair-wise R^2^ values between markers on the same chromosome within flanking regions of ± 250 kbp from the highest-ranked SNP associated with the PC1. SNPs were considered independent when R^2^ < 0.1. We performed priority linkage disequilibrium (LD) pruning by iteratively removing SNPs with weaker associations (lower rank) with the ME^38^. The final selection comprised SNPs within the top 5% of the rank distribution, resulting in a specific number of cross-validated co-eQTLs for each module within networks.

#### MODULE training and application to test dataset

Following the selection of co-eQTLs for genes within modules, we subset LIBD genotypes to derive a set of genetic predictors. To rule out that predictions were driven by residual *cis*-effects, SNPs within ± 1Mbp of the gene start and end coordinates were removed. We fine-tuned the lambda parameter (see Supplementary Methods: Section 1.3), and we extracted non-zero SNP weights and summary statistics to build a prediction database using the “SQLite” R package. We retained in the predictive models only significant genes with adjusted R^2^ ≥ 0.01, p-value < 0.05. We applied the 48 prediction models obtained to the testing genotypes using the MetaXcan^113^ “*Predict.py*” function (https://github.com/hakyimlab/MetaXcan). As de-scribed in the INGENE section, we employed an averaging strategy to consolidate predictions, resulting in a single imputed expression value for each predicted gene.

#### MODULE and INGENE Cross-dataset Training

To increase robustness and minimize dataset-specific signals, *trans*-models were trained independently in each expression/genotype resource (LIBD, GTEx, and CMC) for every available brain region. After training, all three model sets were applied to the same genotype panel (LIBD) to place predictions on a common basis.

We fit elastic-net models using the standard pipeline for INGENE and MODULE, retaining gene models that passed training criteria. For each gene *g*, we computed three predicted-expression vectors over LIBD individuals:

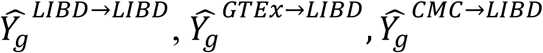

Replicability was assessed only between predicted vectors over the same LIBD individuals (i.e., model-to-model agreement rather than prediction-to-observed expression). For each gene *g*, we computed Pearson correlations:

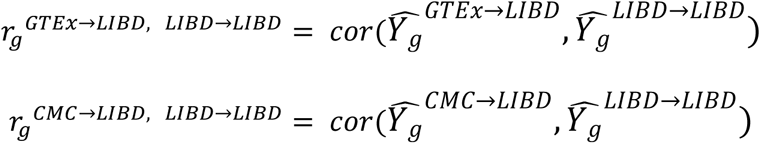

A gene was retained for a region if at least one available cross-dataset correlation was strictly positive (r>0). In practice, we used this filter to **subset the** LIBD-trained models, without using the testing datasets in predictive score generation. This way, only genes whose predictions were replicable in GTEx and/or CMC remained in the final database, while avoiding information leakage from the testing datasets.

### Functional and Regulomic Analysis of genes regulated by *trans*-SNPs identified as GTEx significant *cis*-eQTL

From each tissue-specific MODULE predictive model, we selected *trans*-SNPs predicting genes validated in the GTEx test dataset (refer to the “Evaluation of *cis-* and *trans*-models on a testing dataset” paragraph in the Result section). To enhance the precision of our analysis, we excluded *trans*-SNPs with absolute weight coefficients below 0.01, as these SNPs have minimal regulatory impact on *trans*-associated genes. We then focused on SNPs that were also significant *cis*-eQTLs across all 49 tissues cataloged by GTEx (data available at https://gtexportal.org/home/downloads/adult-gtex/qtl)^56^. The selected SNPs were categorized into quartiles based on the absolute value of their MODULE predictive coefficients and associated with their *cis*-genes from the GTEx catalog (Table S3). This stratification allowed us to investigate how the strength of *trans*-eQTL regulation influenced patterns of functional enrichment. Following categorization, we conducted GO enrichment analyses separately for each brain tissue and the fourth quartile using the R package “ClusterProfiler” and the “enrichGO” function (the universe was defined as all unique genes across GTEx tissues). We focused the GO analyses on molecular function categories (MF), aiming to investigate the specific biological activities enriched among the *cis*-regulated genes.

For the regulome analysis, we used the “gProfileR” package to identify TFs significantly enriched within the list of *cis*-eGenes from the GTEx catalog. We defined the universe of genes as brain-specific genes that are regulated in *trans* by GTEx eQTLs within the MODULE model. To account for multiple comparisons, we applied the FDR correction. Additionally, we conducted a hypergeometric test using the “*phyper*“ R function to evaluate the statistical significance of the observed enrichment between the query set and the defined universe. The p-values obtained from the hypergeometric test were further adjusted using the Bonferroni correction to control for Type I error.

### Correlation between EpiXcan and MODULE SNP weight and PGC3 OR

We downloaded summary statistics from the meta-analysis of European ancestry cohorts from the PGC website to obtain Odds Ratios (ORs) values for SCZ (https://figshare.com/articles/dataset/scz2022/19426775?file=34865091), bipolar disorder (https://figshare.com/articles/dataset/PGC3_bipolar_disorder_GWAS_summary_statistics/14102594) and major depressive disorder (https://figshare.com/articles/dataset/mdd2018/14672085). These ORs were used to study the association with the SNP weights assigned by the Elastic-Net algorithm in the predictive models. First, we computed the predictive mean absolute weights for each SNP predicting genes validated in the testing step in GTEx. Second, we subset the PGC summary statistics to only include SNPs with a significant association with the diagnosis of p < 0.05. We further selected SNPs present in the *cis-* or *trans-*model independently (Table S4). We evaluated the correlation of the weights with log(OR) values using Pearson correlation coefficients, with statistical significance determined by two-tailed tests. To stabilize the variance of correlation coefficients, we employed Fisher’s Z-transformation. First, we calculated the Fisher’s Z-value for CIS, EpiXcan and MODULE models. Next, to account for different numbers of SNPs in the models, we computed the standard error of each Fisher’s Z-value and quantified the difference computing the test statistic Z as:

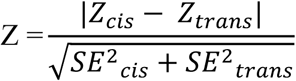

Finally, we derived the two-tailed p-value from the standard normal distribution to assess the statistical significance of the difference. This approach allowed us to robustly compare correlation coefficients, accounting for inherent variability in sample sizes and ensuring the reliability of our findings.

### Model Evaluation

#### ANOVA Analysis in GTEx

To assess the enhanced explanatory power of our combined *cis*- and *trans*- GTEx predictions on the observed gene expression, we performed a maximum likelihood estimation (MLE) analysis of the *full* vs. *null* model (“*anova*” R package) within individual brain regions. The *full* model incorporated observed GTEx expression in logTPM as the dependent variable, and included as covariates sex, mean age, RIN, RNA-rate, overall mapping rate, PMI, the first three PCs, and the first five GEs. We additionally incorporated *cis* and/or *trans* predictions as explanatory variables. The null model included only covariates.

For genes uniquely predicted by a specific model (CIS [C], EpiXcan [E], INGENE [I], or MODULE [M]):

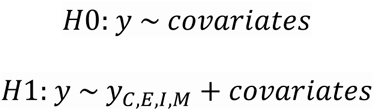

For commonly predicted genes, we applied MLE testing based on the presence of the *cis* term:

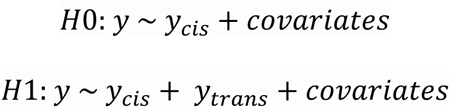

In the absence of the *cis* term (only-*trans* models), the null and alternative hypotheses were:

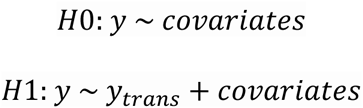

We selected the best model (*cis* only, *cis*-*trans*, or *trans*-only) for each gene based on the most significant increase in predictive power (MLE α = 0.05).

### coTWAS Analysis in PGC3 cohorts

#### Gene expression prediction and selection

For each PGC site, we imputed the expression of all available genes using EpiXcan (E), and the tissue-specific CIS (C), INGENE (I), and MODULE (M) models. As detailed in the “*MODULE/INGENE Training and Application to Test Dataset*” section, we averaged predictions for each gene that was predicted by more than one network within each imputed PGC3 site transcriptome. For further analysis, we retained genes in each tissue-specific model that were validated in the GTEx cohort, meeting the criteria of adjusted R² and Pearson’s r > 0 (refer to the “*Evaluation of cis and trans-models on a Testing Dataset*” section). Genes that met the threshold in only one model were classified as unimodal genes, while those included in at least two models were classified as multimodal genes.

#### Logistic regression analysis

For each PGC site, we performed a gene-wise logistic regression using diagnosis as a dependent outcome and tissue- and model-specific imputed gene expression as an independent predictor while covarying for sex and GEs previously associated with diagnosis by PGC^6^. For unimodal genes, we used the tissue- and model-specific predicted expression as the predictor (Eq. 1). For multimodal genes, we employed the combination of linear models that we trained in GTEx and validated in CMC as detailed in “Combination of *cis*- and *trans*-scores Enhances Gene Expression Prediction in GTEx” (Eq. 2).

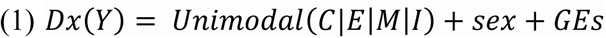

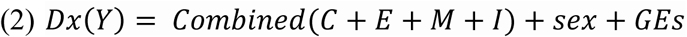

To identify *cis*- and *trans*-predicted genes potentially relevant to SCZ, we focused on gene predictors available in more than 32 PGC3 sites. This threshold balanced gene retention with cross-cohort consistency, avoiding biases introduced by missing SNPs coverage in a subset of cohorts. We then used the β of logistic regression to perform a meta-analysis of effect sizes across sites by using the Stouffer method, applying weights proportional to the square root of the sample size for each cohort, as previously described^118^ and implemented in the “*metap”* R package. To assess heterogeneity, we computed Cochran’s Q test and retained genes with a p-value threshold of p > 1 × 10^−3^ across PGC3 sites. We considered all tissue-gene pairs (96,535 tests).

#### Conditional analysis

To identify independent associations and control for correlation among predicted genes that might arise due to the LD between SNPs used in the different predictive models, we performed a conditional analysis as previously implemented by Huckins et al.^15^. We first identified 315 genomic regions, including the MHC, based on the distance between the 96,535 tissue-gene pairs previously meta-analyzed (1Mb window). As such, each region may contain multiple gene associations and/or genes associated across multiple tissues. To identify independent genic associations within these regions, we carried out a stepwise forward conditional analysis following ‘GCTA-COJO’ theory^119^ using ‘CoCo’, an R implementation of GCTA-COJO. CoCo allows the specification of custom correlation matrices by the user that will be used to filter out correlated genes (default colinear threshold = .9). We generated a predicted gene expression correlation matrix for all tissue-gene pairs within each region, as the root-effective sample size (*Neff*, Eq. 1) weighted average correlation across all cohorts for which the tissue-gene predicted expression was available.

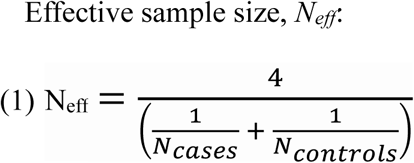

Within each region, forward stepwise conditional analysis of all uncorrelated tissue-gene pairs was carried out using joint linear regression modeling. First, based on the meta-analytic p-value, the top-ranked gene was added to the model, then the next most significant gene in a joint model is added if significant at a given p-value threshold, and so on until either all genes are added to the model or no joint statistic reaches the significance threshold. We then corrected the final output list for genome- wide multiple comparisons using the Benjamini-Hochberg false discovery rate (FDR) method at α = 0.01.

We calculated effect sizes and odds ratios for SCZ-associated genes by adjusting ‘CoCo’ betas to have unit variance (Eq. 2).

GREX beta adjustment:

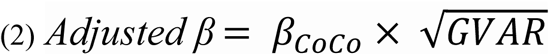

where GVAR is the pooled *Neff* weighted variance of the genetically predicted expression of each gene across all cohorts.

### coTWAS Replicability analysis

Since we did not have access to a PGC-independent dataset, we conducted a leave-site-out replicability analysis across the 62 PGC cohorts to ascertain the robustness of the results obtained from the conditional analysis. For each independent gene-tissue pair (conditional p[FDR] ≤ .01), we implemented two measures of replicability: a) the concordance rate, i.e., the percentage of sites where the β of the logistic regression fitted on all sites but one has the same direction (the same sign) as the β obtained from the model on the held-out site; b) the R^2^ of the correlation between the two type of β across sites.

To assess the statistical significance of the two measures obtained, we reshuffled the β of the held-out sites 10,000 times, and for each gene, we calculated the one-tailed critical value of the two replicability measures at α = .05 (the 95^th^ percentile of the null distribution obtained from the reshuffling). We then selected as replicable genes those with both the concordance rate and R^2^ higher than their respective critical values.

### Cell-type specificity analysis

We used specificity indices derived from single-nuclei RNA-seq of human brains^57^, which discriminated against ten different cell types (neuron and glia). We used the Mean-rank Gene Set Test in *limma* (v3.46) R package^120^ to evaluate the enrichment of our components for the specificity indices of each cell type. FDR correction served to control for multiple comparisons across genes and cell types tested (FDR α = 0.05).

## Data availability

All data supporting the conclusions of this study are provided within the article and its Supplementary Materials. The LIBD post-mortem raw RNA-Seq files from the BrainSeq Phase I, II and III studies for CN, DLPFC, and HP are available through the database of Genotypes and Phenotypes (dbGap) and Globus collections (CN: phs003495.v1.p1 [https://www.ncbi.nlm.nih.gov/projects/gap/cgi-bin/study.cgi?study_id=phs003495.v1.p1 DLPFC: jhpce#bsp2-dlpfc [http://research.libd.org/globus/jhpce_bsp2-dlpfc/index.html]; HP: jhpce#bsp2-hippo [http://research.libd.org/globus/jhpce_bsp2-hippo/index.html]). The LIBD post-mortem raw genotype data are available through dbGap under accession code phs000979.v3.p2 [https://www.ncbi.nlm.nih.gov/projects/gap/cgi-bin/study.cgi?study_id=phs000979.v3.p2]). The RNA-Seq data for amygdala and sACC from the BipSeq study is available through the PsychENCODE Consortium. Access to the data is managed by the NIMH Repository & Genomics Resource (NRGR) and the data are distributed via Synapse under the BipSeq study (syn5844980). All raw and processed data for DLPFC, amygdala and dACC samples from the VA PTSD study may be requested via the PTSD Brain Bank Resource Request process described on the “For Investigators” tab of the following page: https://www.research.va.gov/programs/tissue_banking/ptsd/default.cfm. The GTEx post-mortem processed RNA-seq data used in this study are publicly available at GTEx Portal (release V8). Individual genotype data can be accessed upon request through the database of Genotypes and Phenotypes (dbGaP) under study accession number phs000424.v8.p2. The CMC MSSM-PENN-Pitt RNA-seq data were obtained upon request from Synapse, with synapse numbers syn18097439 for DLPFC (release 3.0) and syn29442240 for ACC (version 6.0). Individual genotype data were accessed under synapse number syn18097441. The PGC3 wave individual-level phenotype and genotype data used in this study were obtained upon request and are subject to controlled access. Access details can be found at https://pgc.unc.edu/for-researchers/data-access-committee/data-access-information/. Researchers must apply for access and comply with the data use agreement. The GWAS summary statistics for SCZ, BP, and MDD used in this study are publicly available at: https://figshare.com/articles/dataset/scz2022/19426775, https://figshare.com/articles/dataset/PGC3_bipolar_disorder_GWAS_summary_statistics/14102594, and https://figshare.com/articles/dataset/mdd2018/14672085. GTEx *cis*-eQTLs data for 59 tissues is publicly available at https://gtexportal.org/home/downloads/adult-gtex/qtl. Additionally, this study employs WGCNA networks previously published. Details of these resources are provided as Supplementary Data 1 and in the Key Resource Table in the Supplementary Information to facilitate reproducibility.

## Code availability

The code used for this study, along with the co-expression prediction models, will be made publicly available. Additionally, this study employs MetaXcan software which is publicly available at https://github.com/hakyimlab/MetaXcan and no customization of the source code was applied. PrediXcan software can be found at https://github.com/hakyimlab/MetaXcan. EpiXcan software can be found at https://bitbucket.org/roussoslab/epixcan/src/master/.

## URLS

https://github.com/theboocock/coco/ “CoCo”, an R implementation of GCTA-COJO.

## Supporting information

Supplementary methods, Supplementary Results and Supplementary Figures

## Contributions

Conceptualization: G.P. and F.R. Data curation: D.R.W., T.M.H., J.E.K, and M.C.O. Formal analysis: F.R., L.S., G.C.K, G.G., F.D.C., D.F., C.B. and D.M. Funding acquisition: G.P., D.R.W., S.P., and B.V. Investigation: F.R. and G.P. Methodology: G.P., F.R., L.S., G.C.K, D.F., and D.M. Project administration: G.P. Resources: D.R.W., G.B., T.M.H., J.E.K., A.R., S.R., A.B., J.K., S.I.B., P.R.M., C.A., J.T.R.W., M.C.O., M.J.O., D.B., A.C., D.W.M., E.D., J.v.O., E.A., M.C.S., M.D.F., B.T.B., C.N.P., A.M., V.G., N.K., V.E.P., A.G., E.K., J.C., M.R., D.C., C.L.L., A.S., O.A.A., D.S.C., T.L., A.K.M., N.S.M., B.J.M., D.R., I.G., A.H., B.K., M.M.N., M.R., G.K., P.F.S., T.L.P., T.W., A.M., T.E., E.G.J., H.E., B.P.R., D.F.L., J.D.B., E.B., C.M.H., R.A.O., R.A., E.A.S., B.P.F.R., S.G., F.T., and C.D.

Software: F.R. Supervision: G.P. Visualization: F.R., L.S., G.C.K., and D.F. Writing (original draft): F.R., G.P., and D.R.W. Writing (reviewing and editing): all authors.

## Acknowledgments

Fabiana Rossi and Daniela Fusco are PhD students enrolled in the National PhD in Artificial Intelligence (resp. XXXVII and XXXVIII cycle) course on Health and Life Sciences, organized by Università Campus Bio-Medico di Roma. We thank Prof. Leonardo Fazio and Prof. Gianmarco Leggio for their scientific contribution to grants that funded this work. We thank Prof. Shizhong Han for his helpful suggestions and review of the manuscript. We thank Prof. Paolo Provero for his scientific insight and review of the manuscript. We are grateful for the contributions of the Office of the Chief Medical Examiner of the State of Maryland, Office of the Chief Medical Examiner of Kalamazoo County Michigan, Office of the Chief Medical Examiner University of North Dakota School of Medicine, Gift of Life of Michigan, Office of the Chief Medical Examiner of Santa Clara County California, and Medical University of Sofia, Bulgaria in assisting the Lieber Institute for Brain Development in the acquisition and curation of brain tissue donations for this study. All research at the Lieber Institute for Brain Development is made possible by generous gifts from the families of Steve and Connie Lieber and Milton and Tamar Maltz. We would like to thank all the family members of the donors for their exceptional contribution. Special recognition goes to Madhur Parihar and Dr. Nora Penzel for providing scientific and technical support.

## Competing interests

A. Bertolino received consulting fees from Biogen and lecture fees from Otsuka, Janssen, and Lundbeck. D. Weinberger serves on the scientific advisory boards of Sage Therapeutics and Pasithea Therapeutics. G. Pergola and G. C. Kikidis received lecture fees from Lundbeck. A. K. Malhotra is a consultant to Genomind, InformedDNA and Concert Pharmaceuticals. M. C. O’Donovan, M. J. Owen, and J. T. R. Walters are supported by collaborative research grants from Takeda Pharmaceuticals. O. A. Andreassen is a consultant for HealthLytix and received speaker’s honoraria from Lundbeck. C. Arango has been a consultant to or has received honoraria or grants from Acadia, Angelini, Gedeon Richter, Janssen Cilag, Lundbeck, Minerva, Otsuka, Roche, Sage, Servier, Shire, Schering Plough, Sumitomo Dainippon Pharma, Sunovion and Takeda.

## Funding

This project was co-funded by the University of Pisa on Fondo di Finanziamento Ordinario (FFO) sponsoring the PhD scholarship: “Identifying biological pathways of psychiatric risk from genes to cognition via AI”. Further support derived from the project “Hot for genes – the role of brain gene expression in identifying developmental trajectories and malleable risk factors for preventive interventions” (PRIN: Research Projects of National Relevance 2020 Prot. 2020WSCSLZ) awarded to B.V., S.P., and G.P. Additional support comes from the project “A Prosocial Training Program to Harness Genes and Environment Influencing Social Behavior” (PRIN 2022, Prot. 2022KXJYJA), awarded to G.P. and S.P., and the project “Unraveling new neural network activities for the treatment of negative symptoms and socio-cognitive abilities in schizophrenia” (PRIN PNRR 2022, Prot. P2022HNBJX), awarded to G.P. The LIBD funded the collection and analysis of postmortem brain tissue.

