## Supplementary methods, Supplementary Results and Supplementary Figures for "Co-expression-based models improve eQTL predictions and highlight novel transcriptome-wide genes associated with schizophrenia"

#### Section 1.1. Ethics statement

The research described herein complies with all relevant ethical regulations. Postmortem human brain tissues from the LIBD collection were primarily obtained by autopsy from the Offices of the Chief Medical Examiner of the District of Columbia and of the Commonwealth of Virginia, Northern District, all with informed consent from the legal next of kin (protocol 90-M-0142 approved by the National Institute of Mental Health (NIMH)/National Institutes of Health (NIH) Institutional Review Board). The National Institute of Child Health and Human Development Brain and Tissue Bank for Developmental Disorders (https://medschool.umaryland.edu/BTBank) provided infant, child, and adolescent brain tissue samples under the NO1-HD-43368 and NO1-HD-4-3383 contracts. Additionally, donations of postmortem human brain tissue from patients with SCZ were provided with informed consent by next of kin from the Office of the Chief Medical Examiner for the State of Maryland under protocol number 12–24 from the State of Maryland Department of Health and Mental Hygiene and from the Office of the Medical Examiner, Department of Pathology, Homer Stryker, Maryland School of Medicine under protocol number 20111080 from the Western Institute Review Board. The Institutional Review Board of the University of Maryland at Baltimore and the State of Maryland approved the study protocol. The Lieber Institute for Brain Development (LIBD) received the tissues by donation under the terms of a material transfer agreement.

The replication cohort of GTEx data adhered to the requirements established in the GTEx pilot study requirments[1]. Explicit authorization for tissue donation was obtained from the next-of-kin or legally authorized representatives, even though research on deceased individuals is not legally classified as human subjects research. Biospecimen Source Sites (BSS) either submitted a research protocol for IRB review or determined that further review was unnecessary. Consent was obtained in person and over the phone, and a sub-study evaluated donor families' concerns regarding the process. Training materials were developed to support ethical interactions with donor families. To safeguard privacy and confidentiality, only de-identified data were shared, with strict access controls based on user roles. The project implemented a Material Transfer and Data Use Agreement to define responsibilities for privacy protection across all parties involved. The template agreement is posted publicly at http://biospecimens.cancer.gov/global/pdfs/caHUB_Material_Transfer_and_Data_Use_Agreement_072512-508.pdf.

For the CMC replication cohort, ethical approvals are detailed in the original paper[2]. Briefly, the “MSSM” brain specimens were obtained through the Mount Sinai NIH Brain Bank and Tissue Repository (NBTR) (http://icahn.mssm.edu/research/labs/neuropathology-and-brain-banking) , which obtains brain specimens from the Pilgrim Psychiatric Center, collaborating nursing homes, Veteran Affairs Medical Centers and the Suffolk County Medical Examiner’s Office. Informed consent is obtained from the next of kin. The brain bank procedures are approved by the ISMMS IRB and exempted from further IRB review due to the collection and distribution of postmortem specimens. The “Pitt” sample were obtained through the University of Pittsburgh Brain Tissue Donation Program during routine autopsies conducted at the Allegheny County Office of the Medical Examiner (Pittsburgh) following the consent of the next of kin. All procedures for Pitt samples have been approved by the University of Pittsburgh’s Committee for the Oversight of Research involving the Dead and Institutional Review Board for Biomedical Research. The “Penn” brain specimens were collected through the University of Pennsylvania Brain Bank of Psychiatric illnesses and Alzheimer’s Disease Core Center (http://www.med.upenn.edu/cndr/biosamplesbrainbank.shtml). All procedures for Penn are approved by the Committee on Studies Involving Human Beings of the University of Pennsylvania, and the use of control postmortem tissues was considered exempted research in accordance with CFR 46.101 (b), item 65 of Federal regulations and University policy.

Ethical approval protocols for each of the 62 PGC wave 3 study sites used in this research are described in the Supplementary Cohort Descriptions from Trubetskoy et al[3].

**Section 1.2. LIBD post-mortem sample description**

Dorsolateral prefrontal cortex (DLPFC) samples were obtained from Brodmann Area (BA) 9/46 at the level of the rostrum of the corpus callosum[4-6]. Hippocampus (HP) samples included the mid-hippocampus proper, with dissections covering the dentate gyrus, CA3, CA2, CA1, and the subicular complex[5]. The caudate nucleus (CN) was dissected from its anterior "head" portion, representing the part most tightly connected to the prefrontal cortex[7]. Amygdala samples[6,8] were dissected from the medial temporal lobe, covering all subnuclei at the level of the largest circumference. Subgenual anterior cingulate cortex (sACC) samples were dissected from the ventral part of the corpus callosum to the dorsal part of the orbital frontal cortex (BA11)[8].

All amygdala, CN, dACC and sACC samples underwent sequencing via the Illumina Ribo-Zero Kit. For DLPFC, RNA-seq was performed using the Illumina Ribo-Zero kit, and the RNeasy kit (Qiagen). RNA-seq for HP samples was performed via the Illumina Ribo-Zero kit and the Illumina TruSeq Stranded Total RNA Library Prep Human/Mouse/Rat kit. When DLPFC or HP data were available with both techniques, we retained those with maximum RNA Integrity Number (RIN) and Ribo-Zero kit protocol.

#### Section 1.3. Lambda tuning (MODULE Training)

We used the same fold indices that we used in the "co-eQTL discovery" **s**tep to fine-tune the Enet lambda parameter. We used the "*lambdaseq*" function from the "method" package[9] to generate a sequence of lambdas for CV testing (alpha = 0.5, lambdaRatio = 1e-02, nLambda = 100). In each training set, we computed the PC1 and projected its loadings onto the testing set to obtain a testing PC1. For reproducibility, we set a seed and generate random 4-fold indices to perform an inner loop CV and tuned the optimal lambda for each outer loop. We fit the model on the projected testing-fold PC1 with the best lambda and computed various performance statics, including mean square error (MSE), $R^2$, adjusted $R^2$ and the Pearson's correlation between the observed target gene-expression level and the predicted fold PC1. After CV, we selected the optimal nested-CV lambda, corresponding to the fold with the minimum average MSE. Before training the final model on the genotype and PC1 computed on all data, we first controlled for the correlation sign between the gene and the PC1 of the co-expression matrix, as the sign of the PC1 can be arbitrary. If Pearson's correlation was negative, we inverted the sign of the PC1. We computed Pearson's correlation between the gene-level expression and the cv model when fitted on the training genotype.

#### Section 1.4. SNP-proxy LD

In this analysis, the objective was to maximize the predictive accuracy of all predictive models by compensating for the absence of predictive SNPs in the testing datasets. To achieve this, we identified SNPs that were in high LD with the missing SNPs from our predictive models. Using the 1000 Genomes Project[10] data as a reference, and the PLINK v2.00[11] we computed the LD values for all potential proxy SNPs within a 500 kbp around each missing SNP with the following command:

--ld-snp-list SNPs_mismatch.txt --ld-window 1000 --ld-window-kb 500 --ld-window-r2 0.8 --out

--pfile g1000/g1000_eur --r-unphased

These proxy SNPs were then integrated into our models as substitutes for the missing SNPs. By incorporating these correlated proxies, we aimed to preserve the integrity and efficiency of our SNP-based predictions, ensuring that our models remain robust and accurate despite variations in SNP availability across different datasets.

**Section 1.5. Sample-size sensitivity analysis for CIS and EpiXcan**

We evaluated the effect of training sample size on *cis*-model performance using LIBD, GTEx, and CMC cohorts. The goal was to estimate how the number of predictable genes and cross-validated accuracy scale with *n*, and whether enforcing cross-dataset replication would disproportionately penalize *cis* models due to reduced effective training size. For every pair of datasets in a region, we identified the intersection of predictable genes and calculated replicability rate as the proportion of shared genes relative to each dataset's total. For each between-dataset comparison we computed the sample-size ratio (max/min) and related it to (i) the absolute difference in mean adjusted $R^2$ between datasets and (ii) the replicability rate. Simple linear models (*lm* function in R) were fit with sample-size ratio as the predictor; 95% confidence intervals, significance and $R^2$ were obtained from the model fits.

#### Section 1.6. PGC weight Ratio and Connectivity Enrichment

To evaluate whether genes more heavily influenced by SCZ-associated variants exhibit increased connectivity to PGC3-prioritized genes[3], we performed a permutation-based test of monotonic enrichment across quintiles of a PGC-weight metric.

**PGC-weight Metric Calculation.** For each gene predicted by CIS, EpiXcan, and MODULE models, we computed a PGC-weight metric defined as:

$$\text{PGC-weight ratio} = \left(\frac{\sum_{\text{PGC3 SNPs}} |\text{ model weight }|}{\sum_{\text{all SNPs}} |\text{ model weight }|}\right) \times \sqrt{\frac{\text{number of PGC3 SNPs}}{\textit{total number of SNPs}}}$$

This metric captures both the relative contribution and density of SCZ-associated SNPs ($p < 0.05$ in PGC3 summary statistics[3]) to the gene's expression prediction.

**Connectivity Quantification.** Gene-wise connectivity scores were obtained from Borcuk et al.[12] who constructed transcriptomic co-expression networks across five brain regions (amygdala, DLPFC, CN, HP, sACC). Each score reflects the strength of connectivity between a given gene and the set of 120 PGC3-prioritized genes[3].

**Quintile Binning and Trend Assessment.** Genes were binned into five equal-sized quintiles based on their PGC-weight metric. For each model-region pair, we computed the mean connectivity within each quintile, resulting in a 5-point trajectory.

To assess monotonic trends, we calculated:

- Linear regression slope across quintile indices (1 to 5)
- Spearman's rank correlation coefficient (ρ) between quintile index and mean connectivity

**Permutation Testing.** To generate a null distribution, we performed 1,000 gene-level permutations per model-region pair. In each permutation:

1. Genes were randomly reassigned to quintiles (preserving bin sizes).
2. Mean connectivity was recalculated per permuted quintile.
3. Both the linear slope and Spearman's ρ were recomputed.

Empirical two-sided *p*-values were defined as the proportion of permuted statistics whose absolute value equaled or exceeded the observed statistic. This approach does not assume linearity and directly tests whether the observed ordering of quintile means departs from chance.

**Section 1.7. MAGMA Z-score derivation**

We performed gene-based association analyses using MAGMA v1.09b[13] to derive gene-level Z-scores. SNP-to-gene mapping was conducted with the default MAGMA annotation files, using a window of 35 kb upstream and 10 kb downstream of each gene. As input, we used SNP *p*-values from the PGC3 SCZ GWAS summary statistics[3], together with the 1000 Genomes European reference panel[10] to account for linkage disequilibrium. MAGMA aggregates SNP-level association statistics within the defined cis-window, correcting for LD, to produce a gene-level *p*-value, which is then transformed into a standardized Z-score. In our study, these Z-scores were considered the MAGMA importance score, representing the strength of cis-genetic association between each gene and SCZ risk.

### Supplementary Notes

#### Section 2.1. Cross-dataset replication of predictive models

To assess the reproducibility of predictive weights across independent brain collections, we evaluated training-level performance and cross-dataset replication for CIS, EpiXcan, INGENE, and MODULE using LIBD, GTEx, and CMC data.

**Training-level overview.** Sample sizes varied substantially, from 116 donors in CMC amygdala to nearly 600 in LIBD DLPFC (Fig. S3A), affecting eQTL discovery power and cross-validation metrics[14,15]. We compared the number of predictable genes (Fig. S3B) and cross-validated performance across datasets (Fig. S3C). Across regions, both *cis*- and *trans*-models exhibited heterogeneous CV adjusted $R^2$ values that reflected sample-size differences rather than intrinsic model behaviour. Mean CV adjusted $R^2$ ranged from 0.11–0.78 for CIS, 0.10–0.32 for EpiXcan, 0.10–0.34 for INGENE, and 0.15–0.20 for MODULE (Fig. S3C). Because such variability may inflate apparent model fit, we next quantified independent cross-dataset replication for INGENE and MODULE models as a more reliable measure of generalizability.

**Cross-dataset replication.** We assessed reproducibility by comparing predictions across datasets using LIBD as the common reference (see Methods – *MODULE and INGENE Cross-dataset Training* for detailed procedures). Briefly, each gene predicted expression values from GTEx, or CMC were correlated with the corresponding predictions from LIBD. Two directional comparisons were performed: (i) GTEx → LIBD versus LIBD → LIBD, and (ii) CMC → LIBD versus LIBD → LIBD. Genes were retained as replicable if the Pearson correlation between external and

LIBD predictions exceeded 0. This design isolates cross-cohort consistency while holding sample size and model complexity constant.

Across brain regions, both INGENE and MODULE displayed broad cross-dataset overlaps (Fig. S2A-B). For INGENE, reproducible gene counts ranged from 9,590 in amygdala to 16,560 in DLPFC. For MODULE, reproducibility ranged from 5,807 in HP to 12,599 in sACC. In regions represented in both external datasets (dACC, DLPFC, sACC), GTEx and CMC each contributed comparably to the reproducible gene pool: for INGENE, GTEx-validated genes accounted for 57–78 % and CMC-validated for 87–90 % of the total reproducible set (shared overlap ≈ 50–65 %); for MODULE, the respective contributions were 64–70 % for each dataset (shared overlap ≈ 30–40 %) (Fig. S2A–B).

Anchoring replication to LIBD as a shared reference and applying this correlation-based filtering ($r > 0$) delineated a subset of gene-level predictions consistently reproduced across independent post-mortem brain cohorts. This subset was carried forward for downstream postmortem validation and coTWAS analyses.

**Section 2.2. Sample-size sensitivity of cis-based predictors**

To evaluate the influence of training-sample size on the reproducibility of *cis*-based models, we examined cross-cohort consistency for CIS and EpiXcan predictors across LIBD, GTEx, and CMC datasets (see Methods: *Sample-size sensitivity analysis for CIS and EpiXcan*). Analyses quantified the relationship between training-sample size ratios and cross-dataset differences in prediction accuracy (Fig. S4).

**1. CIS shows strong sample-size dependence.** Across brain regions, CIS performance differences scaled positively with the ratio of training-sample sizes ($R^2 = 0.476$, $p = 0.019$; Fig. S4A). At the gene level, retention of overlapping predictions declined proportionally to sample-size imbalance (Fig. S4B). In the DLPFC, where LIBD and CMC sample sizes were comparable (see Fig. S3A), CIS retained ~53% of overlapping genes. In more unbalanced comparisons, retention dropped to ~25%. These results indicate that CIS predictors are particularly sensitive to differences in discovery-cohort size and may overfit small training datasets.

**2. EpiXcan exhibits reduced sample-size sensitivity.** EpiXcan showed no significant association between sample-size ratio and cross-cohort performance differences ($R^2 = 0.033$, $p = 0.62$; Fig. S4C). In the balanced DLPFC comparison (CMC–LIBD), EpiXcan retained > 55 % of overlapping genes, and in unbalanced regions the decline was modest (Fig. S4D). Across all regions, retention ranged from ~35 % to 60 %, consistently exceeding CIS.

**3. Limited overlap across cohorts.** Despite improved stability, overall cross-cohort overlap remained modest. Even under balanced sample sizes (e.g., CMC–LIBD DLPFC), only about half of overlapping genes were retained, and in most other pairs both CIS and EpiXcan lost 70–85 % of predictions. This limited reproducibility underscores the impact of dataset-specific variance on *cis*-based models.

#### Section 2.3. Benchmarking with BGW-TWAS and MOSTWAS

To benchmark our *cis*- and *trans*-aware predictive frameworks against previously established models, we compared them with BGW-TWAS[16] (models downloaded from https://www.synapse.org/Synapse:syn22316791/wiki/605024) and MOSTWAS[17] (weights downloaded from https://zenodo.org/records/4314067), two reference methods that incorporate distal regulatory information. Analyses were performed in the DLPFC, the only brain region where all reference models were available.

**BGW-TWAS model integration and application in testing datasets.** The BGW models, trained in the ROS/MAP cohort[16], comprised approximately 28 million SNPs (6 million unique). SNP genomic coordinates were lifted from GRCh37 to GRCh38, and multi-allelic variants and indels were removed (842,781 unique and ~3.5 million non-unique values excluded). To optimize prediction in the GTEx and CMC testing datasets, we incorporated proxy variants in linkage disequilibrium (LD $r^2 \geq 0.8$) for missing SNPs. Final SNP weights were computed as the product of the estimated effect size (β) and its posterior probability (pp), i.e., $\beta/SE \times pp$[16], and stored in an SQLite database compatible with the MetaXcan[18] *Predict.py* implementation. Only SNPs present in each testing dataset were retained, and genes were classified based on the regulatory origin of their predictors as *cis*, *trans*, or *cis* + *trans*.

In GTEx, BGW predicted 34 *cis*, 12,540 *cis* + *trans*, and 1,523 *trans* genes (total = 14,097). In CMC, the respective counts were 59 *cis*, 12,513 *cis* + *trans*, and 1,518 *trans* genes (total = 14,090). Predicted gene expression values were matched to observed DLPFC expression by intersecting common genes (GTEx: 14,062; CMC: 13,501) and scaled prior to analysis.

Performance was quantified by Pearson's r and adjusted $R^2$ using linear regression, computed separately for *cis*, *cis* + *trans*, and *trans* genes. In GTEx, 6,098 had Pearson's $r > 0$ and adjusted $R^2 > 0$ (N *(cis)* = 21; N(*cis* + *trans)* = 5.581; N(*trans)* = 496), with a mean adjusted $R^2$ of 0.05. In CMC, 5,849 met the same thresholds (N(*cis)* = 32; N (*cis* + *trans)* = 5.366; N(*trans)* = 451), yielding a mean adjusted $R^2$ of 0.04 (Fig. S7C-D).

**MOST-TWAS models integration and application in testing datasets.** Across all genes, MeTWAS incorporated 673,247 SNPs (445,816 unique) and DePMA 700,843 SNPs (536,183 unique). Because allele and coordinate information were not provided, these were retrieved from BGW annotations and the GRCh37 reference genome, designating the effective allele as the alternative. SNP genomic coordinates were lifted from GRCh37 to GRCh38, and multi-allelic SNPs and indels were removed (N = 75,569 unique for both models). *MetaXcan* was applied directly to the weights, and all remaining preprocessing steps were identical to those used for BGW-TWAS. For MeTWAS, *cis*/*trans* annotations were corrected based on genomic distance, with variants located ±1 Mb from the gene start/end (or on another chromosome) reclassified as *trans*. Of 910 genes provided, 755 passed quality control and were retained for analysis.

For MeTWAS, in GTEx predictions were obtained for 278 *cis*, 460 *cis* + *trans*, and 9 *trans* genes (total = 747).; in CMC, 282 *cis*, 455 *cis* + *trans*, and 8 *trans* genes were predicted (total = 745). Predicted expression values were matched to observed DLPFC expression (GTEx: 697 genes; CMC: 624) and scaled prior to evaluation. In GTEx, 268 genes (N(*cis)* = 111; N (*cis* + *trans)* = 154; N(*trans)* =3), had adjusted $R^2 > 0$ (mean $R^2 = 0.02$), while in CMC, 275 (N (*cis)* =128; N (*cis* + *trans)* = 144; N (*trans)* =3), satisfied thresholds (mean adjusted $R^2 = 0.012$).

For DePMA, 2,941 genes were provided without *cis*/*trans* annotation and were classified using the same genomic-distance rule (± 1 Mb form gene start/end as *trans*). In GTEx, predictions were obtained for 136 *cis*, 2,223 *cis* + *trans*, and 567 *trans* genes (total = 2,926). In CMC, 214 *cis*, 2,081 *cis* + *trans*, and 615 *trans* genes were predicted (total = 2,910).

Predicted values were matched to observed DLPFC expression (GTEx: 2,710 genes; CMC: 2,412) and scaled. In GTEx, 1,122 (N(*cis)* = 67; N(*cis* + *trans)* =885; N(*trans)* = 170), had Pearson's $r$ >0 and adjusted $R^2 > 0$ (mean adjusted $R^2 = 0.032$), while in CMC, 1,094 (N(*cis)* =106; N (*cis* + *trans)* = 828; N(*trans)* = 160), survived both thresholds (mean adjusted $R^2 = 0.023$) (Fig. S7C-D).

### Supplementary Figures

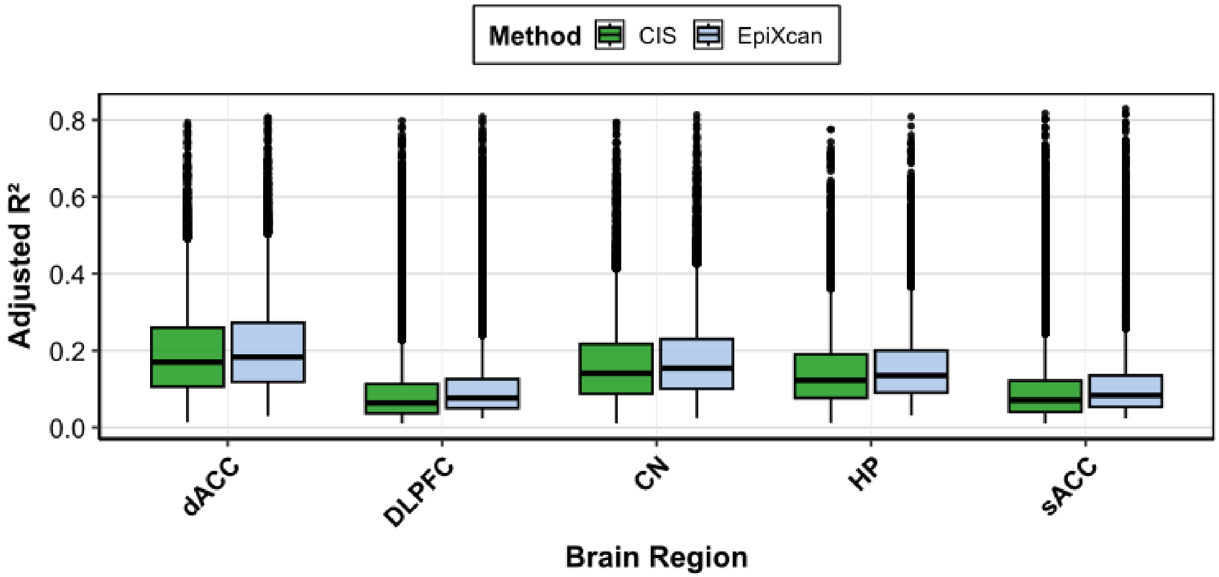


***S1. EpiXcan improves cis-based prediction accuracy across brain regions.*** *Boxplots show the distribution of cv adjusted $R^2$ values for CIS (green) and EpiXcan (light blue) models trained in five LIBD brain regions (caudate nucleus [CN], dorsal anterior cingulate cortex [dACC], dorsolateral prefrontal cortex [DLPFC], hippocampus [HP], and subgenual anterior cingulate cortex [sACC]).*

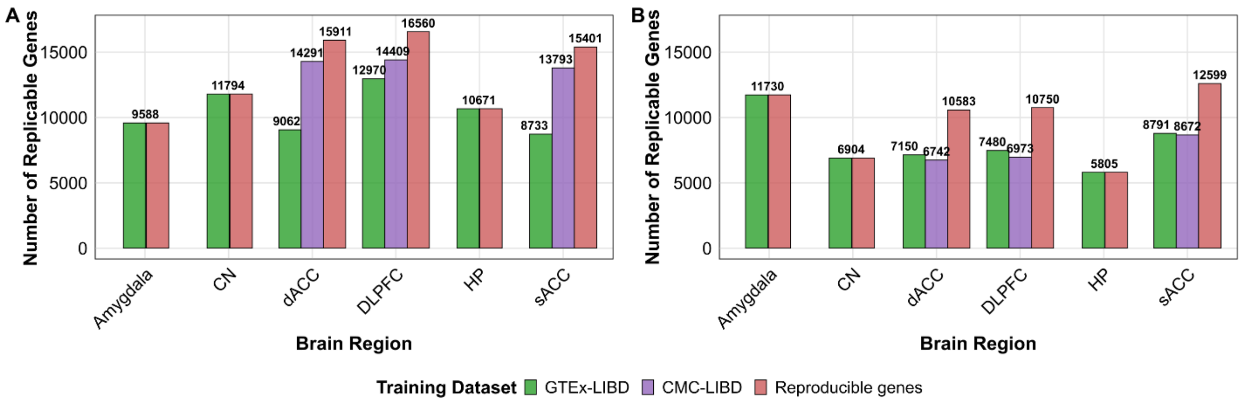


***S2. Cross-dataset replication of trans-aware predictive models across brain regions. A)*** *INGENE and* ***B)*** *MODULE models were trained in LIBD and independently tested in GTEx and CMC to evaluate reproducibility of gene-level predictions across cohorts. Bars show the number of genes passing the replication criterion ($r > 0$ between external and LIBD-predicted expression) in GTEx → LIBD (green), CMC → LIBD (violet), and the total number of reproducible genes (red). CN =*

caudate nucleus; dACC = dorsal anterior cingulate cortex; DLPFC = dorsolateral prefrontal cortex; HP = hippocampus; sACC = subgenual anterior cingulate cortex.*

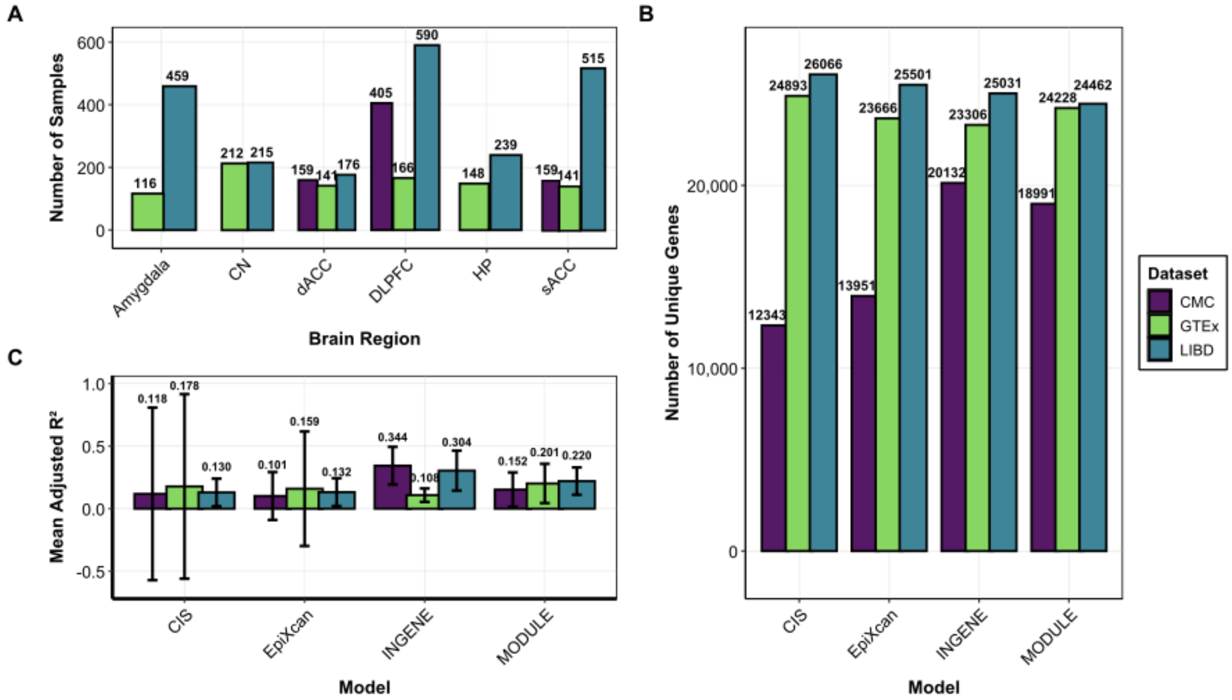


***S3. Training performance of predictive models across cohorts and brain regions**. A) Sample sizes available for training across brain regions in CMC, GTEx, and LIBD, highlighting substantial variability in statistical power. B) Number of unique genes within predictive models across cohorts for each framework (CIS, EpiXcan, INGENE, MODULE). C) Mean cross-validated adjusted $R^2$ values across datasets, showing high variability for CIS and EpiXcan, and greater stability for INGENE and MODULE.*

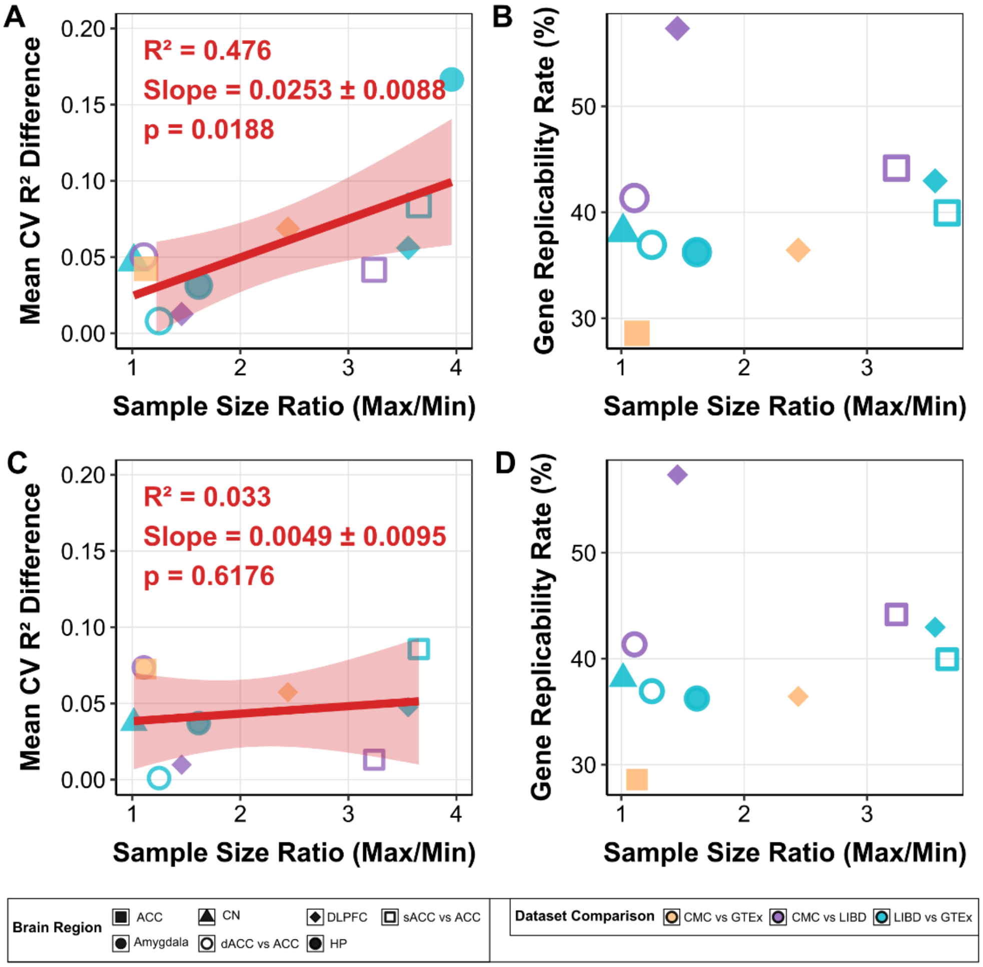


***S4. Sample-size sensitivity of CIS and EpiXcan predictors***. *A) Relationship between sample-size ratio (maximum/minimum) and cross-cohort performance differences for CIS models, showing strong sensitivity to heterogeneity ($R^2 = 0.476$, $p = 0.019$).* ***B)*** *Gene-level replicability rates for CIS across cohort comparisons, with retention declining as sample-size imbalance increases. N represents the number of genes that pass the cross-dataset filtering threshold.* ***C)*** *Relationship between sample-size ratio and performance differences for EpiXcan models, showing no significant association ($R^2 = 0.033$, $p = 0.62$).* ***D)*** *Gene-level replicability rates for EpiXcan across cohorts consistently outperforming CIS but still show limited overlap between datasets. N represents the number of genes that pass the cross-dataset filtering threshold.*

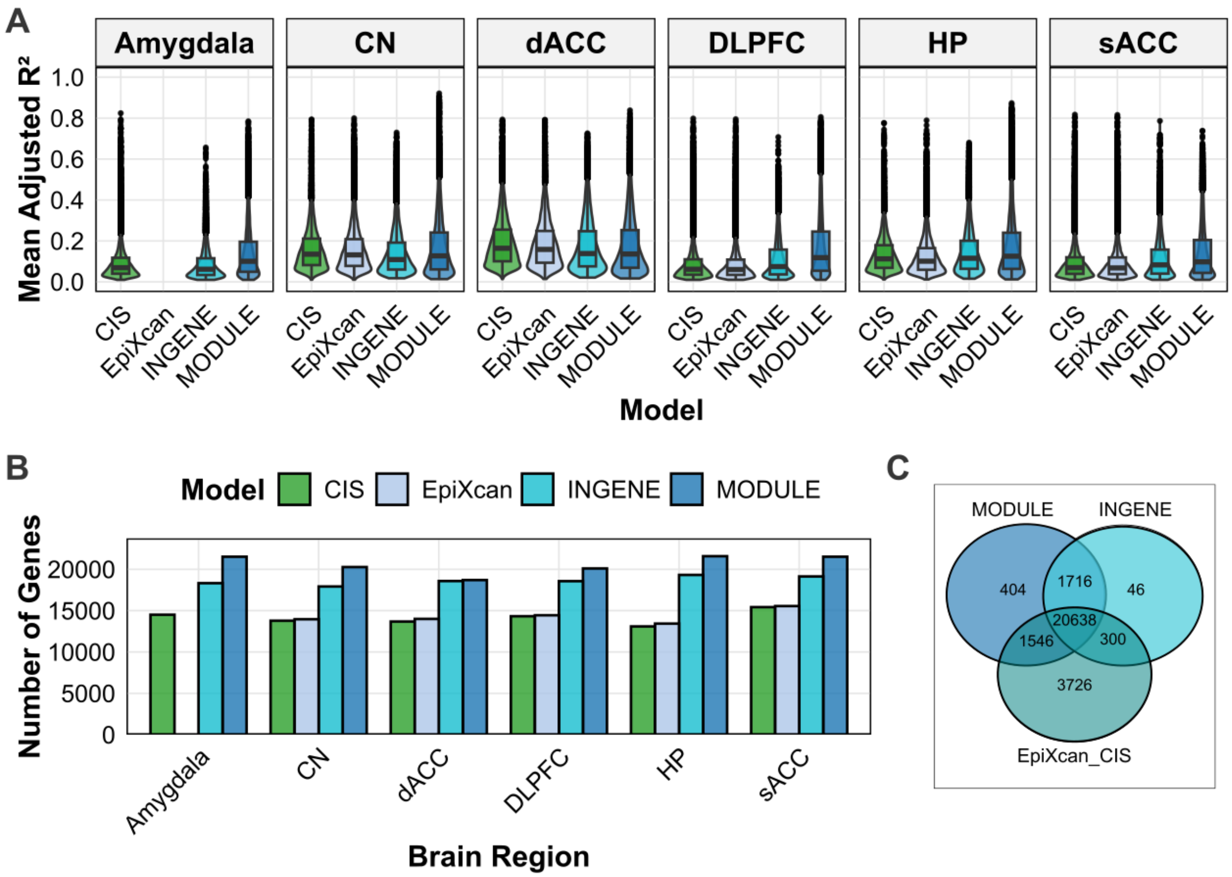


***S5. Comparison of CIS, EpiXcan, INGENE and MODULE model training performances. A)*** *Violin/Boxplot plots show the distribution of cv adjusted $R^2$ values for CIS (green), EpiXcan (light blue), INGENE (cyan), and MODULE (dark blue) models across brain regions. The median is represented by the central line, with the interquartile range (IQR) as the box. Whiskers extend to 1.5×IQR, and outliers are plotted as individual points.* ***B)*** *Barplots report the number of genes with significant prediction models* ***C)*** *Venn diagram showing overlap among genes predicted by the combined cis models (EpiXcan + CIS) and trans models (INGENE, MODULE).*

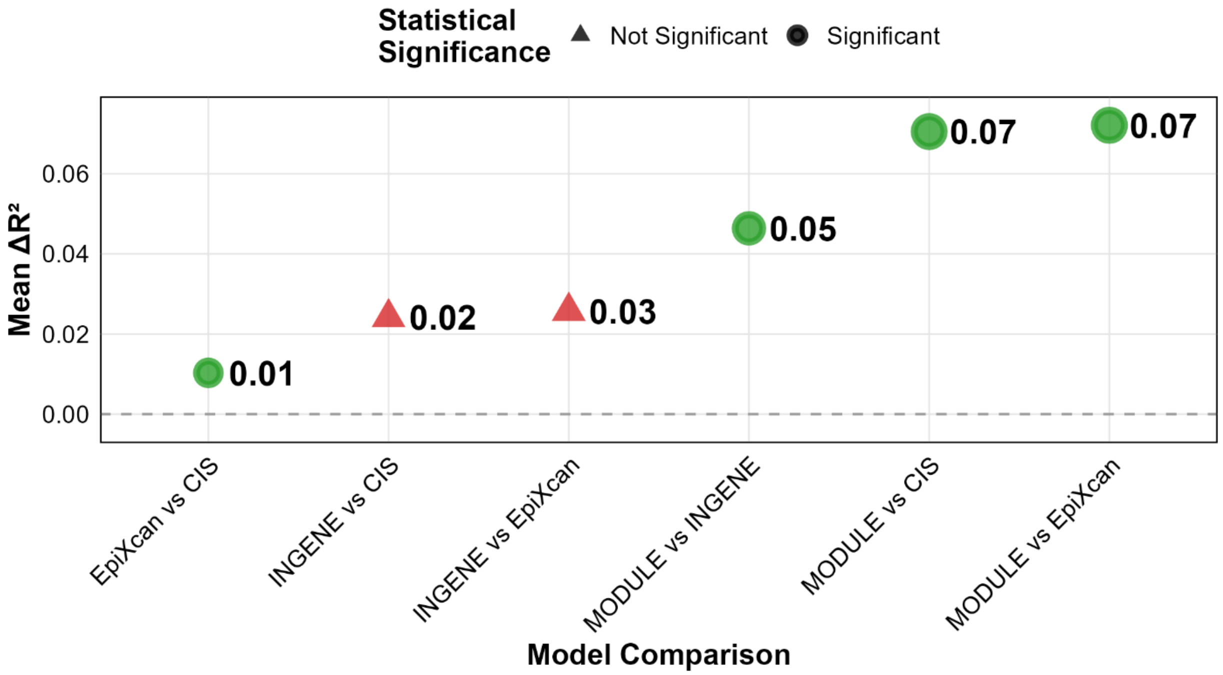


***S6. Pairwise comparison of predictive performance across models.*** *Mean differences in adjusted $R^2$ ($\Delta R^2$) between models are shown for genes commonly predicted across brain regions. Green circles denote statistically significant differences (Wilcoxon one-tailed test, $p < 0.05$), and red triangles indicate non-significant comparisons.*

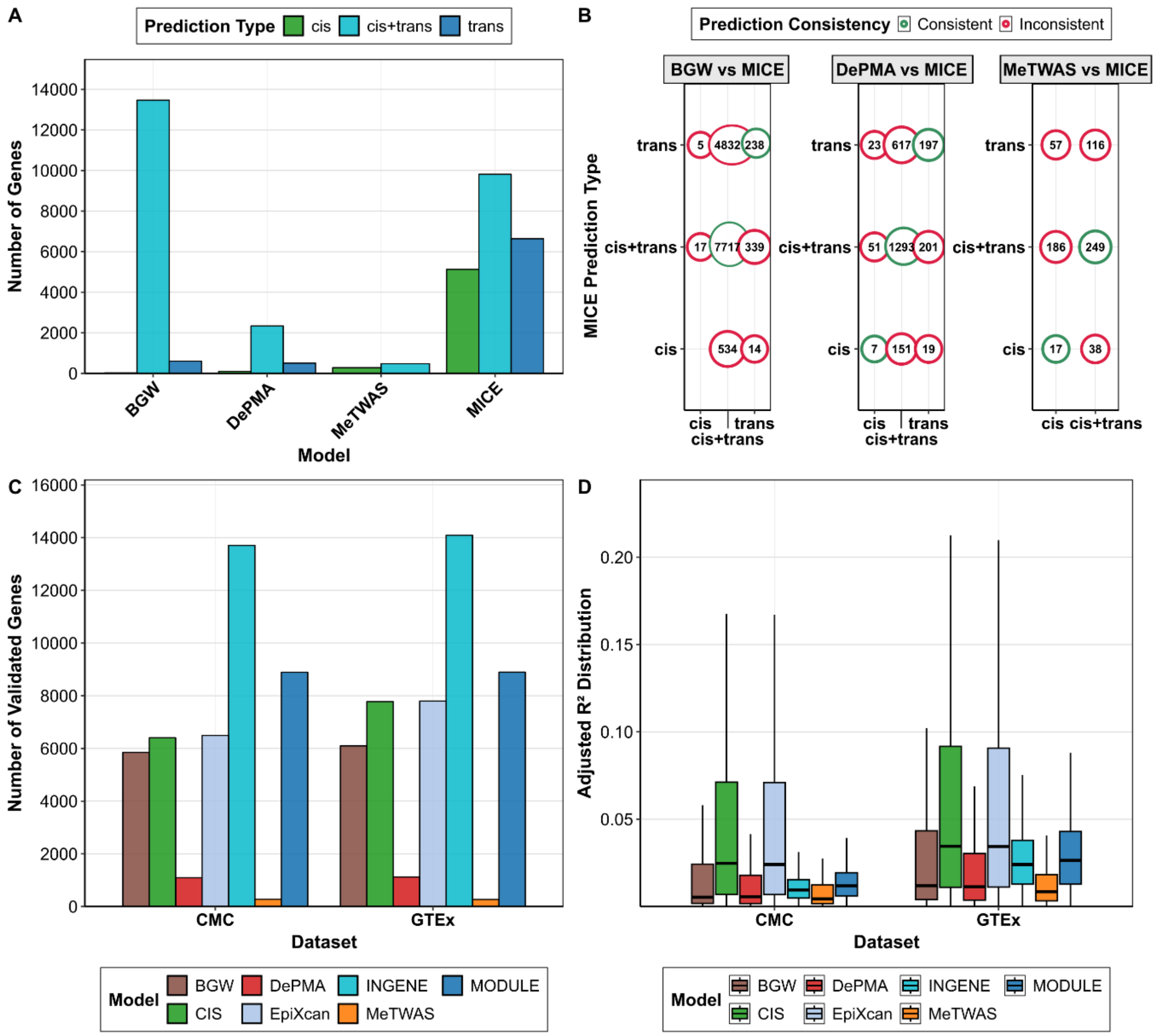


***S7. Comparison with MOSTWAS and BGW-TWAS models.*** *A) Number of genes with trained prediction models across frameworks. B) Consistency of gene-level predictions between MICE and MOSTWAS/BGW-TWAS, measured in terms of predicted genes common across techniques. C) Number of validated genes in independent cohorts (CMC and GTEx) across models. D) Distribution of adjusted $R^2$ values for validated genes in CMC and GTEx, illustrating predictive performance across frameworks.*

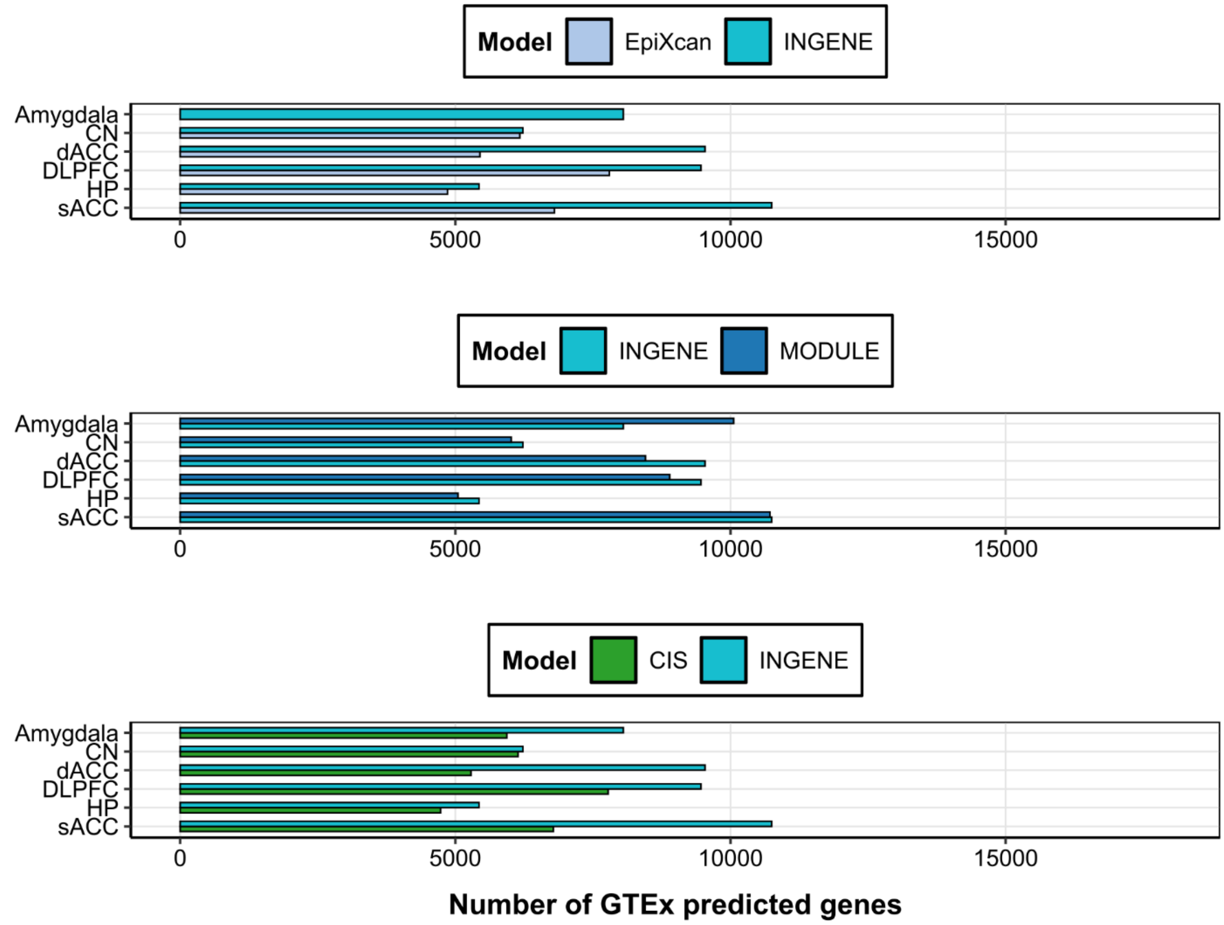


***S8. Predictive models replicate across brain regions in GTEx external dataset and predict different genes at different performance.*** *Barplots show the number of predicted genes (x axis) in the GTEx dataset by CIS (green), EpiXcan (light blue), INGENE (cyan) and MODULE (dark blue) models.*

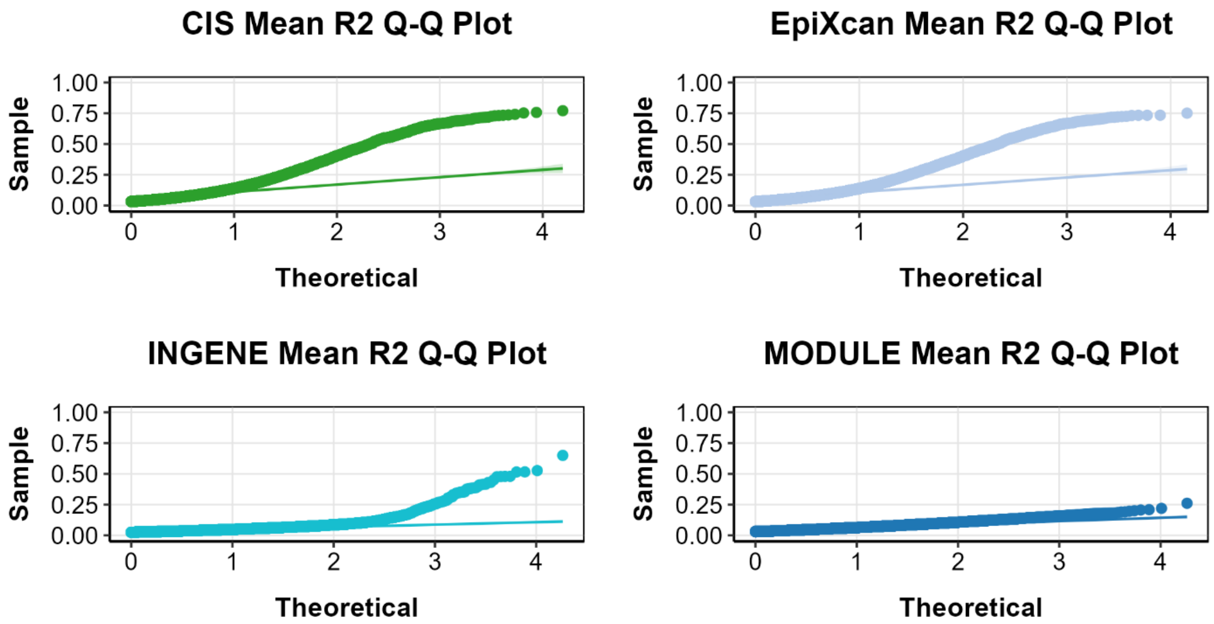


***S9. Comparison of model performances in GTEx testing dataset.*** *Quantile-Quantile (Q-Q) Plot comparing the distribution of adjusted R2 values (y-axis) for CIS (green), EpiXcan (light blue), INGENE (cyan) and MODULE (dark blue) when applied in GTEx. Theoretical quantiles correspond to the null distribution.*

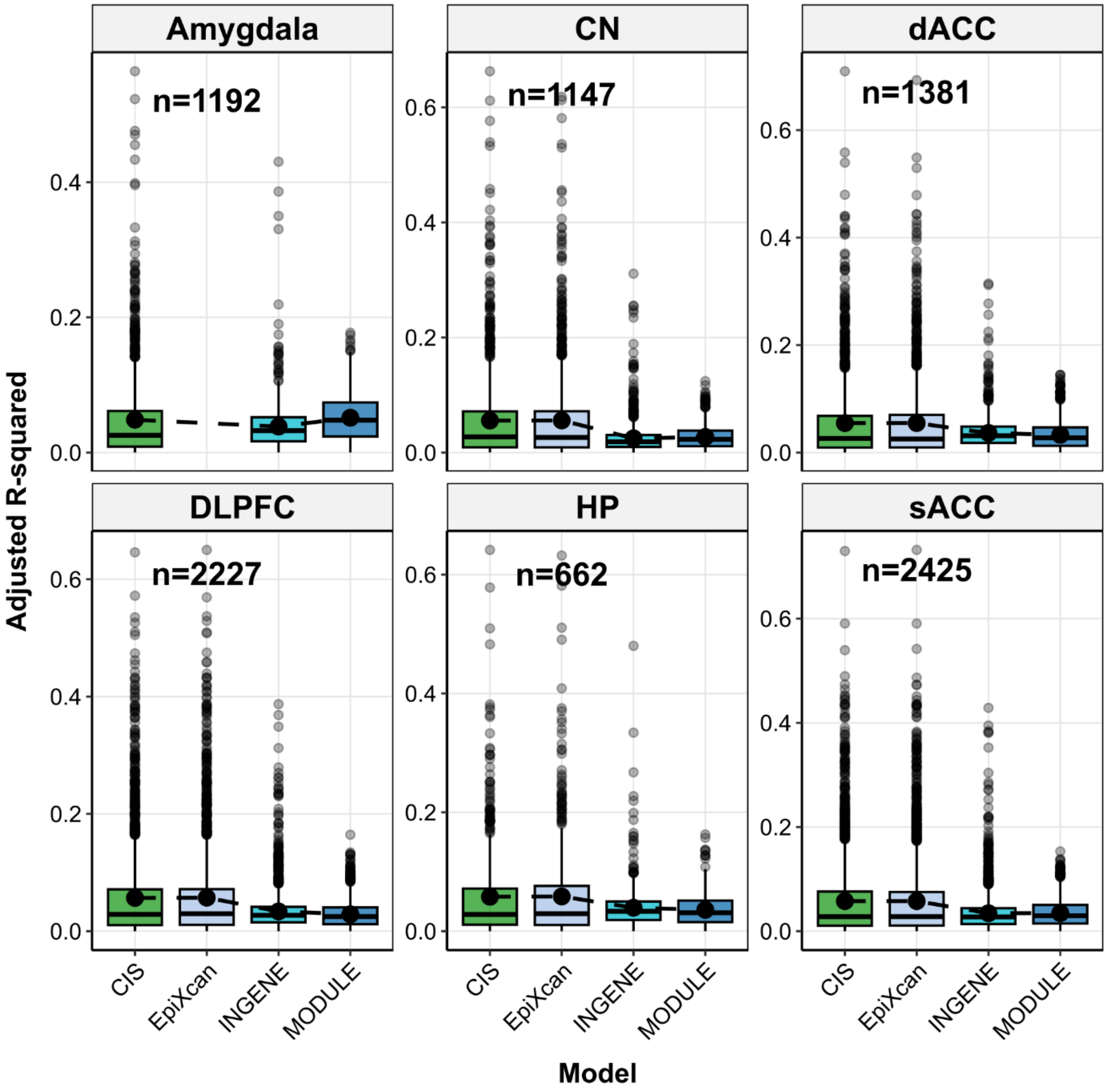


***S10. Models predict common genes at different performance across brain regions in GTEx external dataset***. *Box plots of adjusted $R^2$ values (y-axis) in predicting gene-level expression in GTEx using CIS (green), EpiXcan (light blue), INGENE (cyan) and MODULE (dark blue) for commonly" n" predicted genes within brain regions. The median is represented by the central line, with the interquartile range (IQR) as the box. Whiskers extend to 1.5×IQR, and outliers are plotted as individual points.*

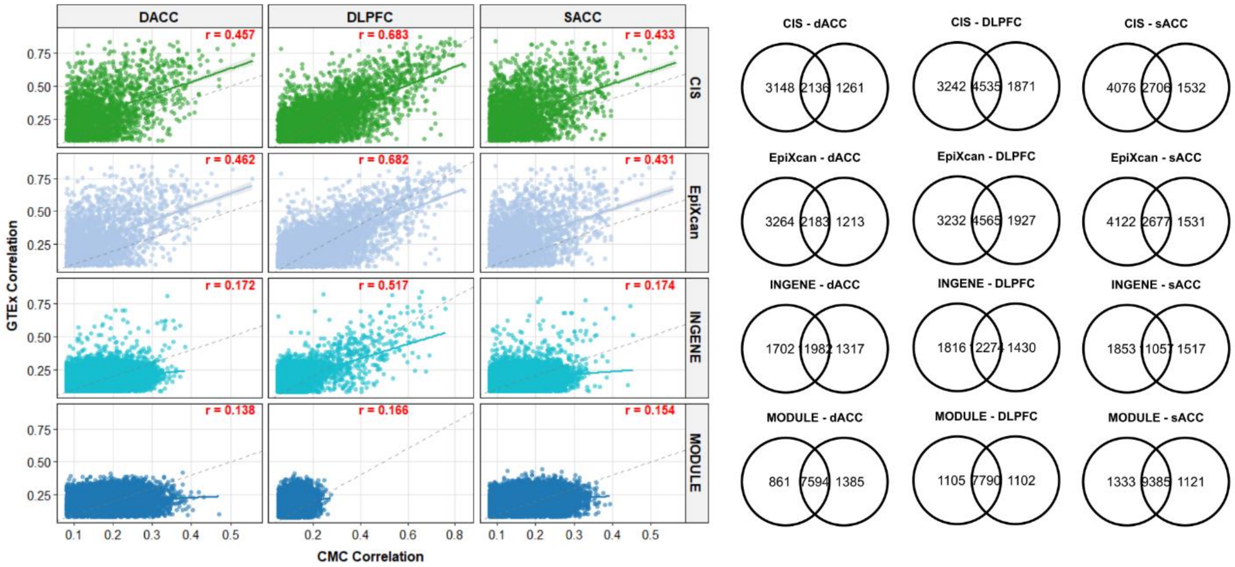


***S11. Cross-cohort replicability of cis- and trans-prediction models.*** *Prediction performance was compared between LIBD-trained models evaluated in CMC and GTEx. Scatterplots (left) show the correlation of predictive accuracy (Pearson's r) across cohorts for CIS (green), EpiXcan (blue), INGENE (cyan), and MODULE (dark blue). Venn diagrams (right) display the number and percentage of genes predicted in both cohorts relative to the total predicted in at least one.*

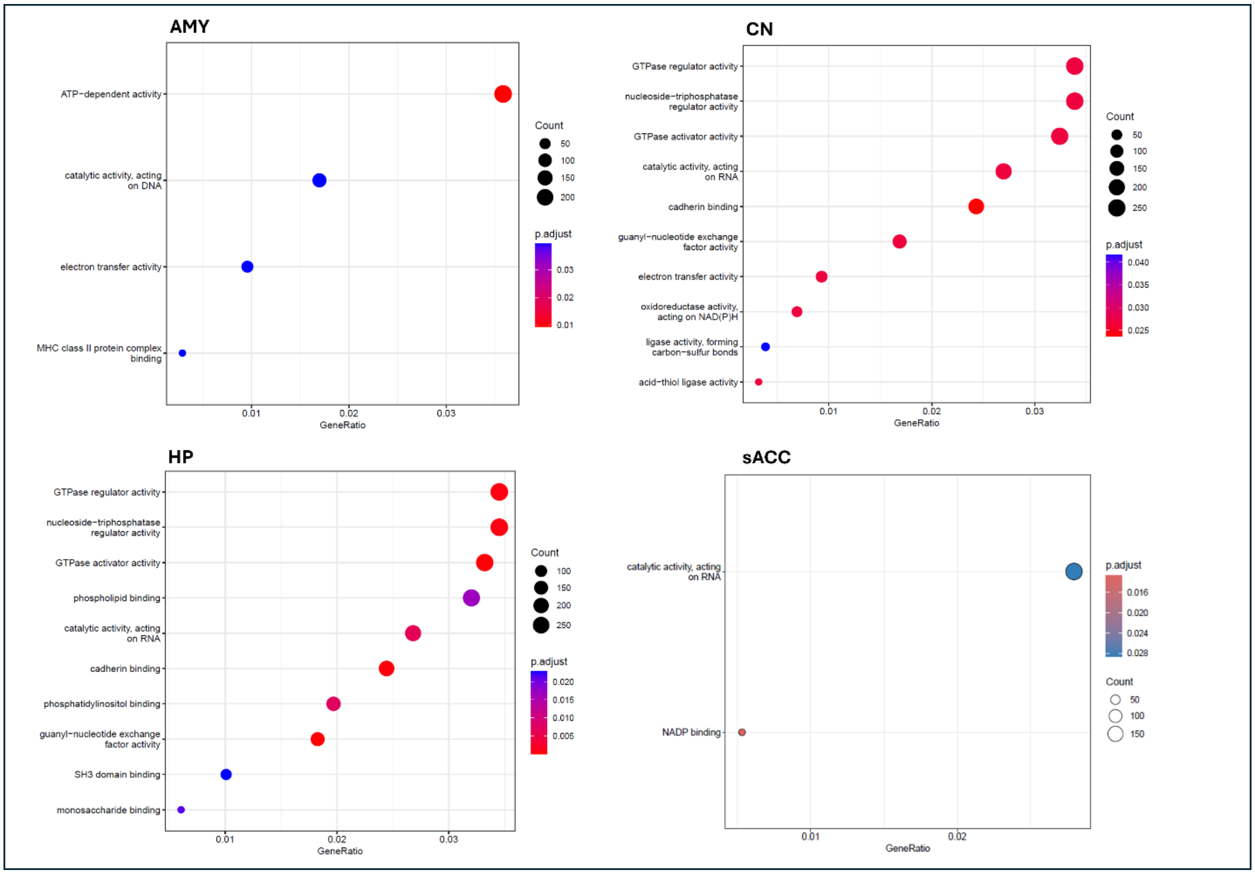


***S12. GO Enrichment Analysis on GTEx eGenes.*** *The x-axis shows the gene ratio for each molecular function category (y-axis). P-adjusted values refer to BH correction. Abbreviations: AMY: amygdala; CN: caudate nucleus bulk tissue data; HP: hippocampus bulk tissue data; sACC: subgenual anterior cingulate cortex bulk tissue data.*

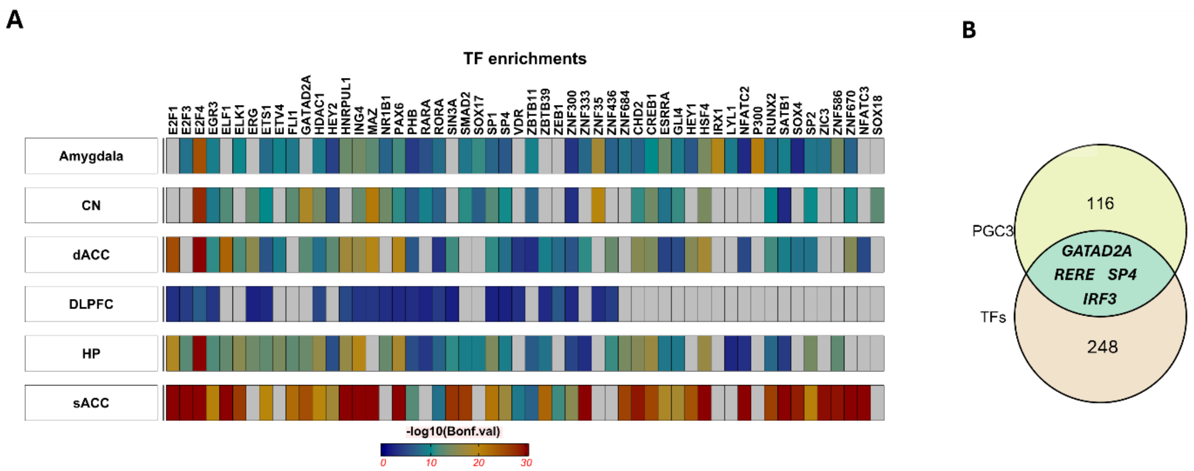


***S13. Regulome Enrichment Analysis of GTEx cis-eGenes for MODULE Trans-eQTLs****. **A)** Enrichment for TFs across brain regions. To generate this visualization, we identified the top 20 most significant TFs for each brain region and assessed their overrepresentation. A grey block in the figure denotes that TF is not significantly overrepresented in that region. **B)** Venn diagram of the overlap between the overrepresented TFs and the 120 prioritized genes from the PGC3 across regions.*

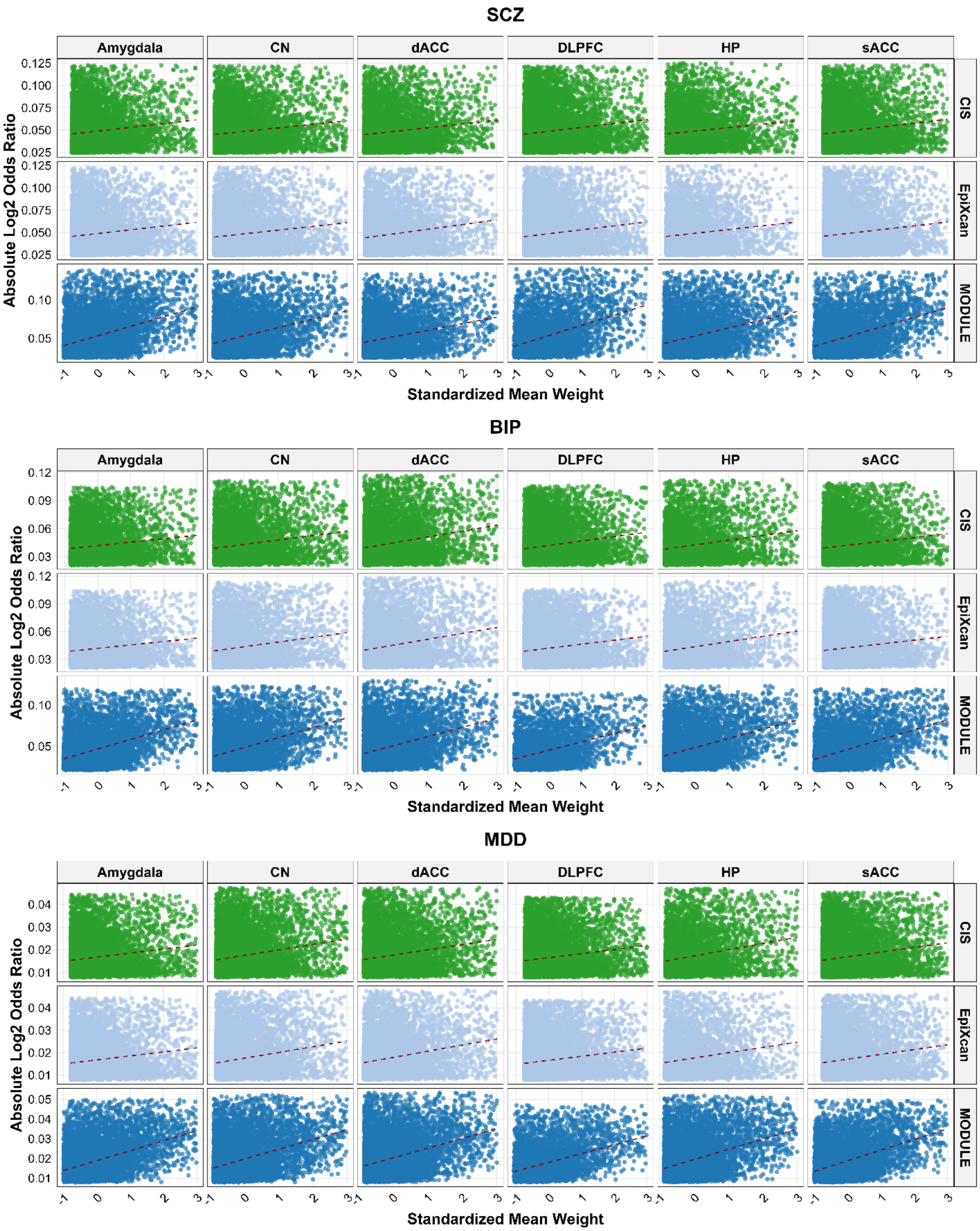


***S14. Scatterplot of PGC3 (p-value < 0.05) SNP log(OR) and mean weight absolute value values in CIS, EpiXcan and MODULE .*** *The x-axis shows the absolute Z-score mean weight of each SNP in CIS (green), EpiXcan (light blue) and MODULE (dark blue), while the y-axis represents the*

*absolute value of the log2 of PGC3 Odd Ratios. Abbreviations: SCZ: schizophrenia; BP: bipolar disorder; MDD: major depressive disorder; CN: caudate nucleus bulk tissue data; dACC: dorsal anterior cingulate cortex bulk tissue data; DLPFC: dorsolateral prefrontal cortex bulk tissue data; HP: hippocampus bulk tissue data; sACC: subgenual anterior cingulate cortex bulk tissue data.*

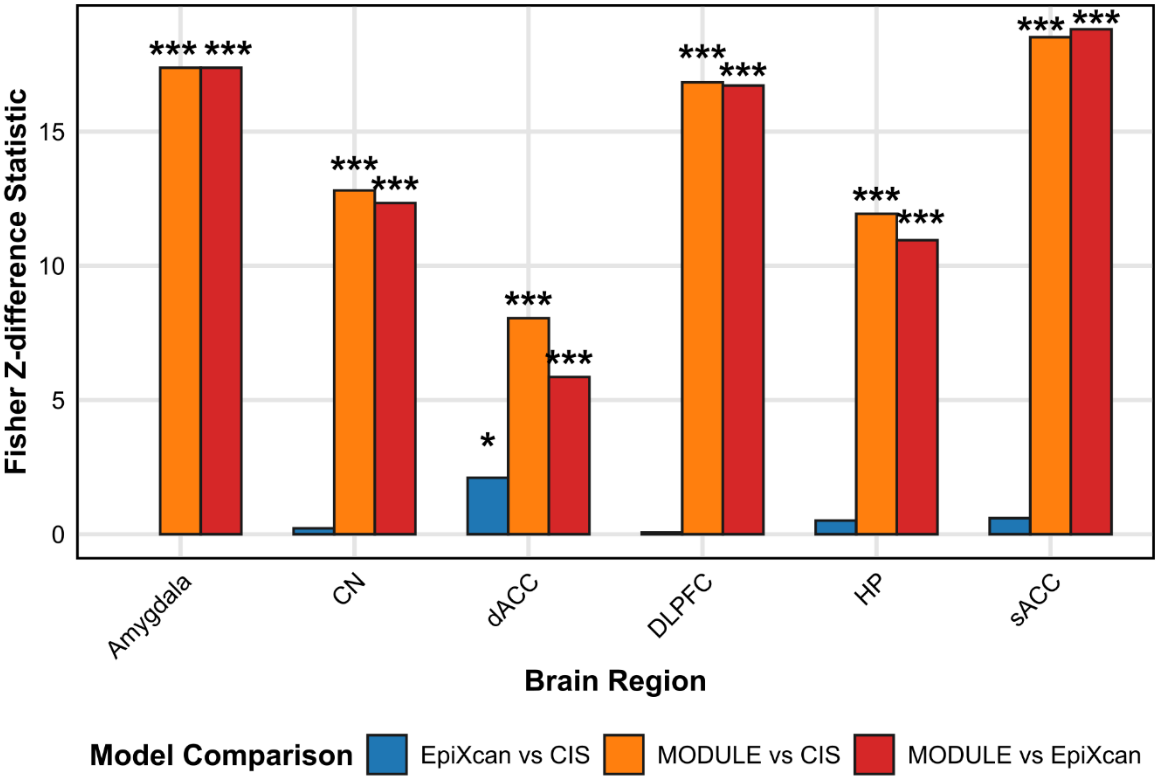


***S15. Statistical Comparison using Fisher's Z-transformation of model correlations between SCZ PGC3 log odds ratios (logOR) and model's mean absolute weight values.*** *Bar plot showing the Z-difference statistics from Fisher's Z-transformation test comparing correlation coefficients between pairs of prediction models (CIS, EpiXcan, and MODULE) across brain regions. Significance levels are indicated by asterisks: * p < 0.05, ** p < 0.01, *** p < 0.001.*

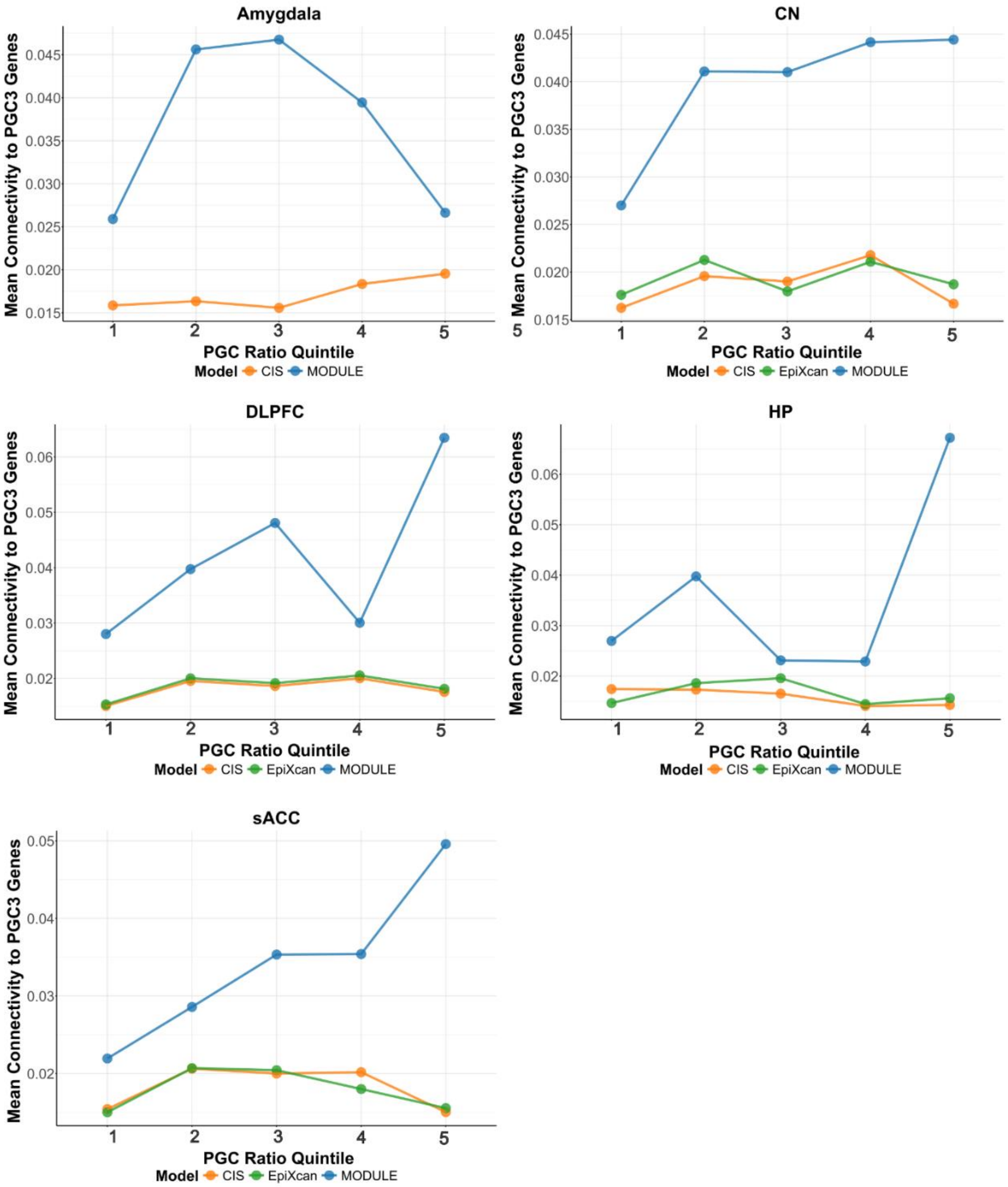


***S16. Mean Connectivity measure of MODULE PGC SNPs ($p < 0.05$) predicted genes with the 120 PGC3 prioritized genes.*** *Connectivity measure published by Borcuk et al.[12]. Abbreviations: CN: caudate nucleus bulk tissue data; DLPFC: dorsolateral prefrontal cortex bulk tissue data; HP: hippocampus bulk tissue data; sACC: subgenual anterior cingulate cortex bulk tissue data.*

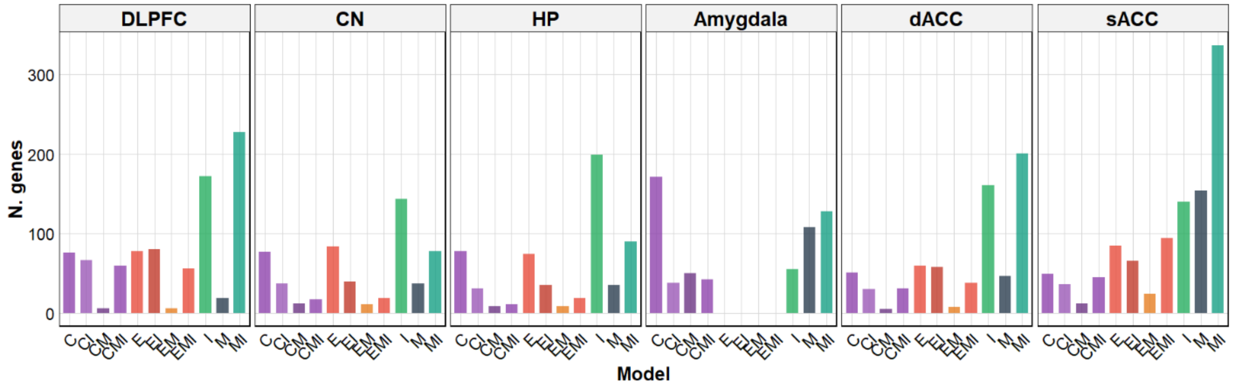


***S17. Distribution of predicted genes by brain region and model across PGC3 cohorts***. *Barplot shows the number of predicted genes pooling predictions from all predictive models and PGC3 cohorts. Abbreviations: C: CIS, E: EpiXcan; M: MODULE; I: INGENE; CN: caudate nucleus data; dACC: dorsal anterior cingulate cortex; DLPFC: dorsolateral prefrontal cortex; HP: hippocampus; sACC: subgenual anterior cingulate cortex.*

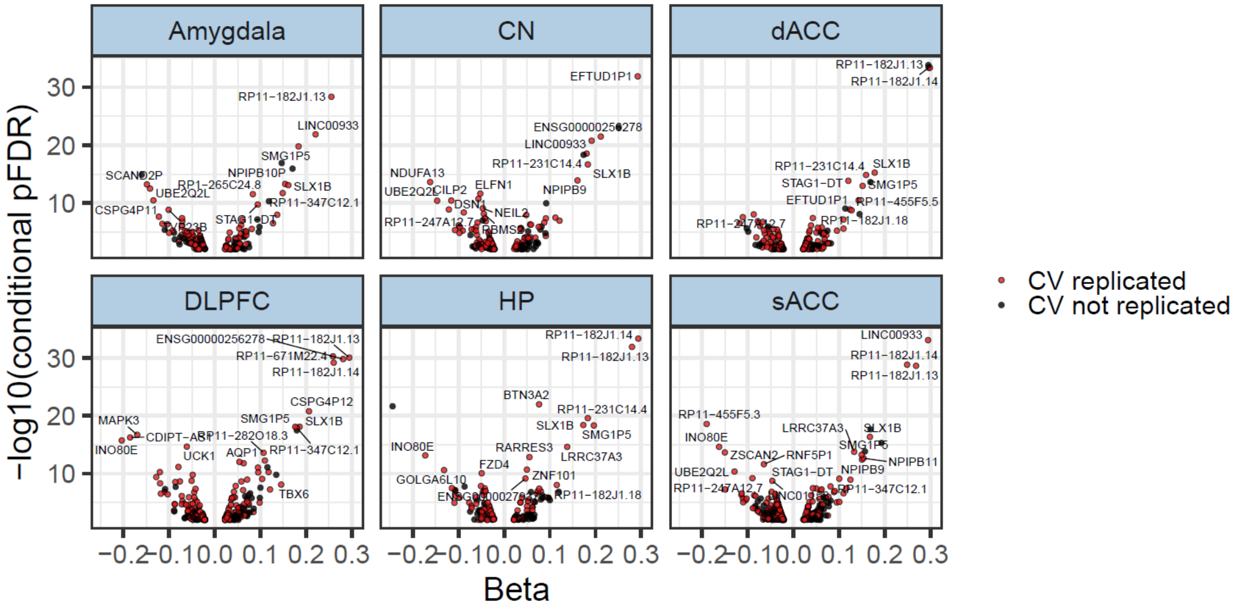


***S18. Cross-validation replicability of coTWAS associations across brain regions.*** *Scatterplots show the relationship between effect size (BETA) and statistical significance (−log$_{10}$ conditional p$_{(FDR)}$) for significant gene–tissue pairs across brain regions. Each point represents a gene identified by coTWAS. Red points indicate genes with associations replicated in the leave-site-out CV analysis, while black points denote non-replicated genes.*

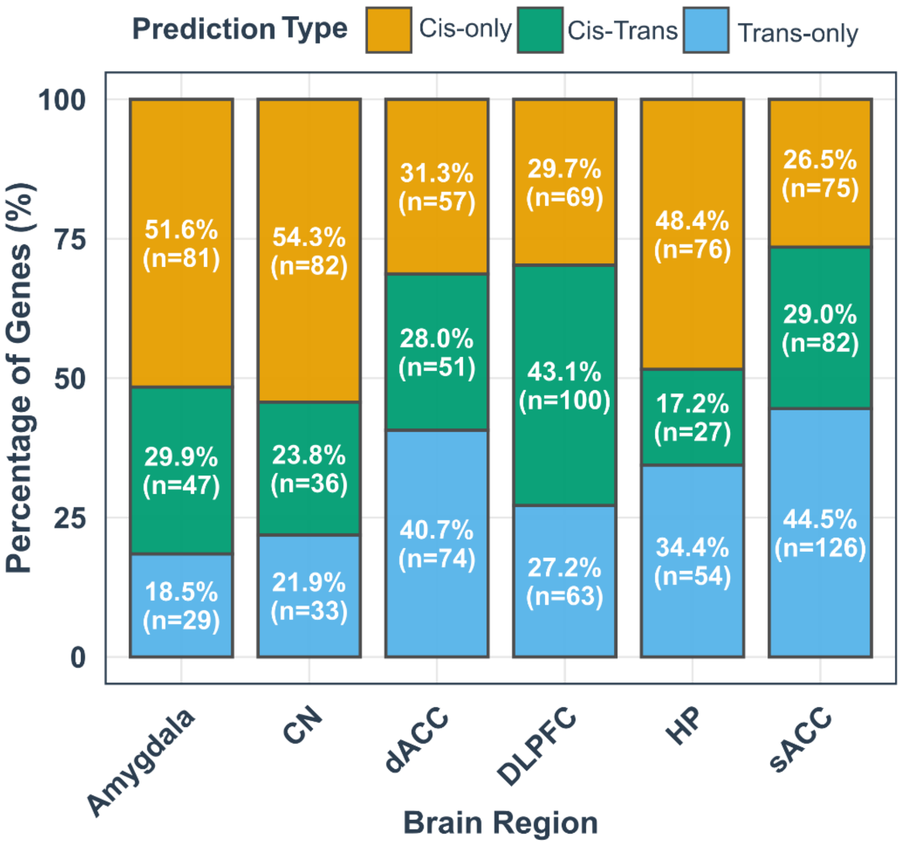


***S19. Distribution of coTWAS FDR 0.01 significant genes with cis-only, trans-only, and cis-trans components.*** *Distribution of prediction types as percentage of genes within regions (n represents the absolute gene count).*

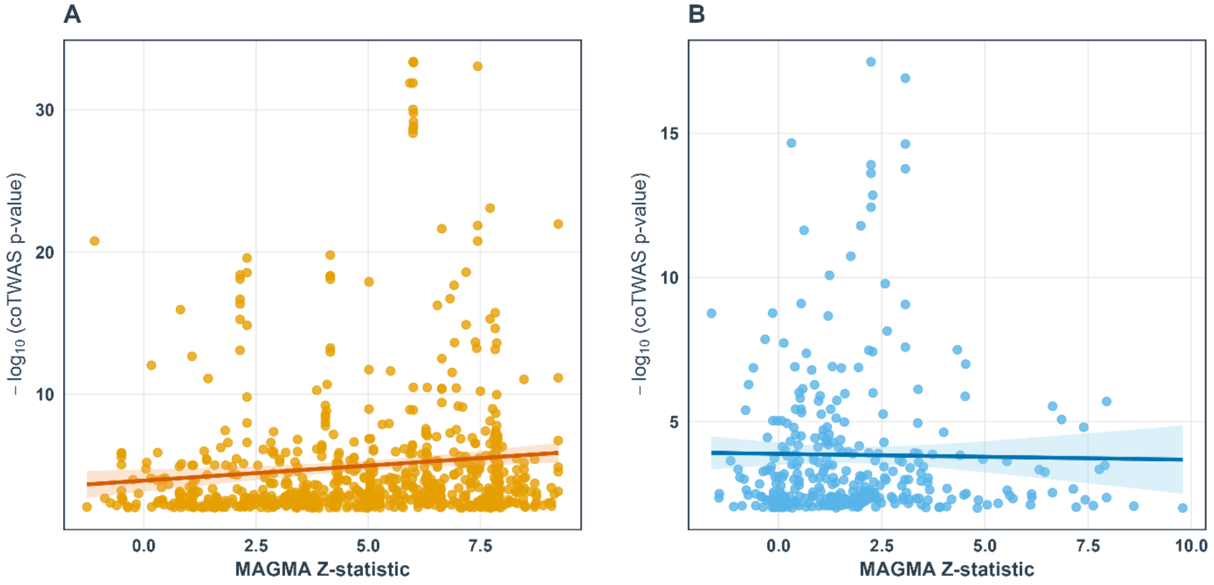

***S20. Correlation between MAGMA Z-score and coTWAS results**. Scatter plots showing the relationship between MAGMA Z-statistics and coTWAS -log10 p, separately for cis-derived (**A**) and trans-derived (**B**) predictions.*

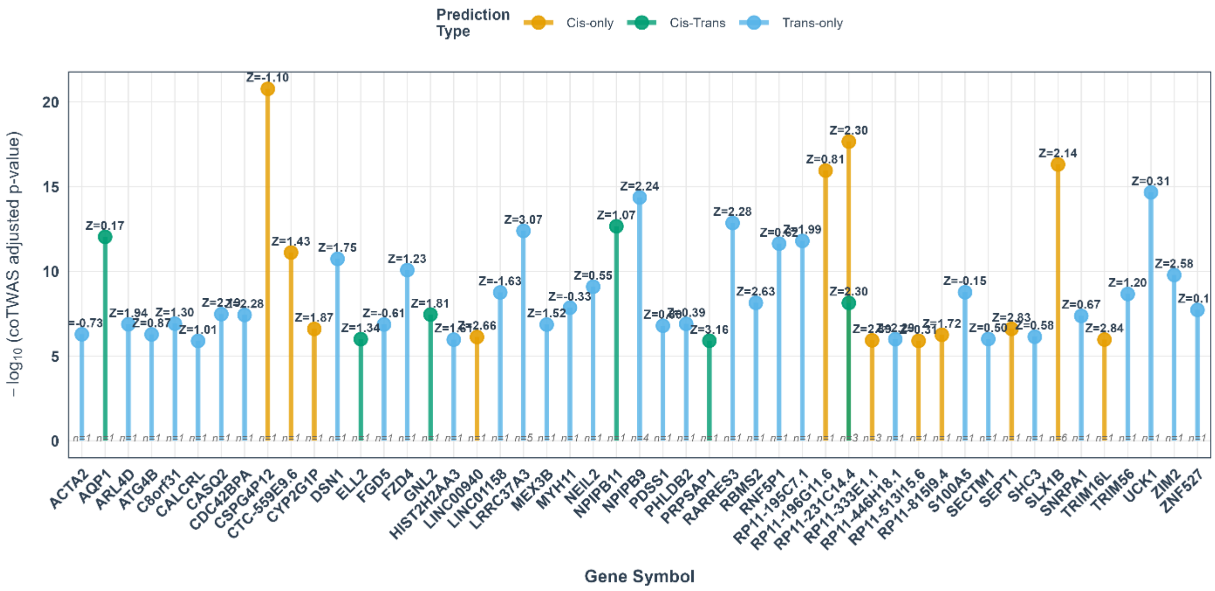


***S21**. Top 50 genes with strong coTWAS evidence (-log10 adjusted p > 4) but weak MAGMA association (|Z| < 4). Z-scores shown above each point indicate MAGMA statistics.*

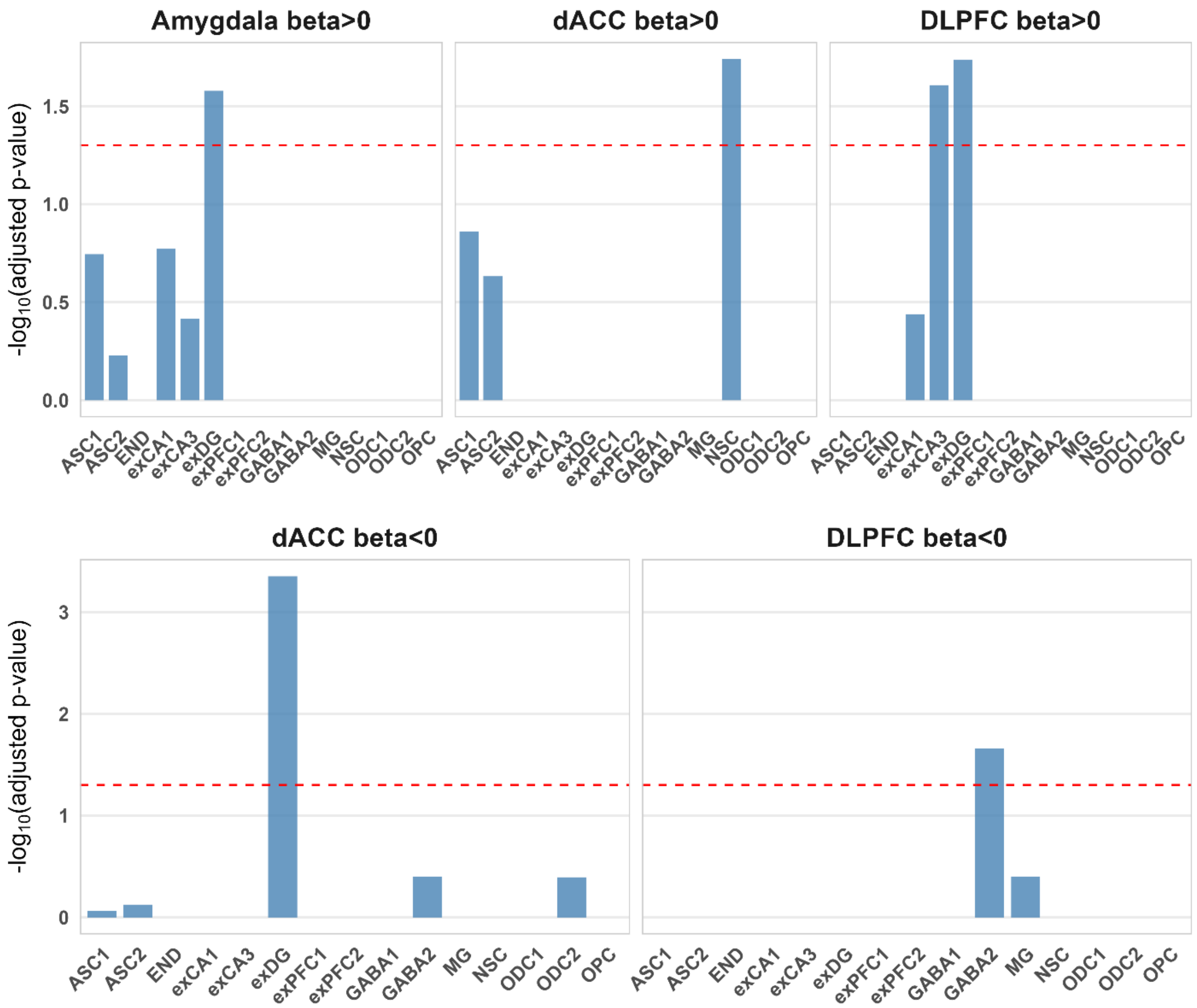


***S22. Cell-type specificity of coTWAS significant genes using human single-cell atlas***. *Mean-rank Gene Set Test in the limma R package was used to obtain the enrichment p-values shown. y-axes show FDR-adjusted p-values after correcting multiple comparisons across genes and cell types. Red dashed lines represent FDR α = 0.05. Top panel shows upregulated (Adjusted Beta > 0); bottom panel shows downregulated (Beta <0) genes in individuals with SCZ based on the logistic analysis.*

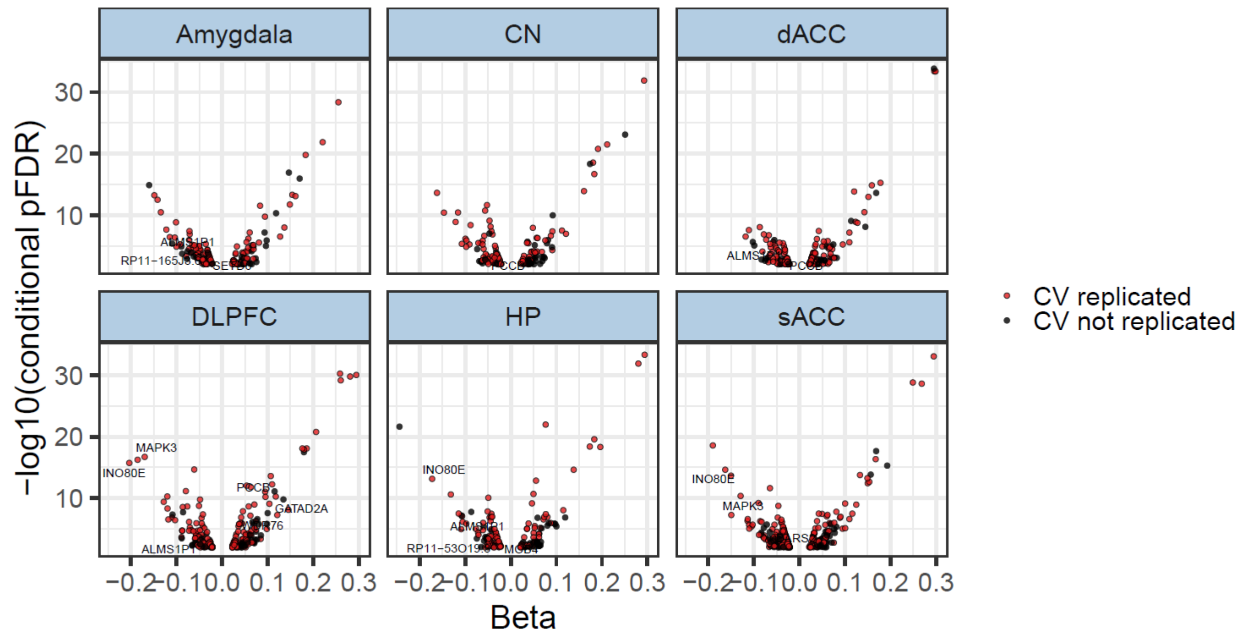


***S23. Intersection between PGC3 120 prioritized genes and coTWAS significant genes across tissues****. Scatterplots show the relationship between effect size (BETA) and statistical significance ($-\log_{10}$ conditional $p_{(FDR)}$) for significant gene–tissue pairs across brain regions. Each point represents a gene identified by coTWAS. Red points indicate genes with associations replicated in the leave-site-out cross-validation (CV) analysis, while black points denote non-replicated genes. Gene Symbols represent the intersection of genes between the sets.*

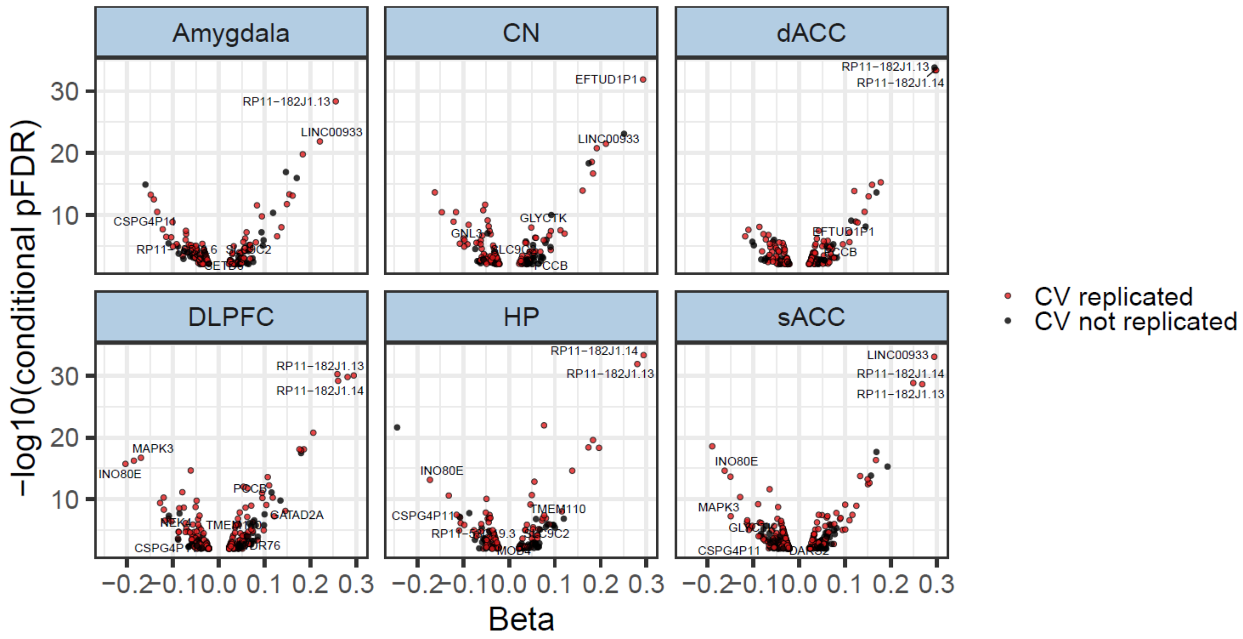


***S24. Intersection between PGC3 Mendelian Randomization genes and coTWAS significant genes across tissues***. *Scatterplots show the relationship between effect size (BETA) and statistical significance ($-log_{10}$ conditional $p_{(FDR)}$) for significant gene–tissue pairs across brain regions. Each point represents a gene identified by coTWAS. Red points indicate genes with associations replicated in the leave-site-out cross-validation (CV) analysis, while black points denote non-replicated genes. Gene Symbols represent the intersection of genes between the sets.*

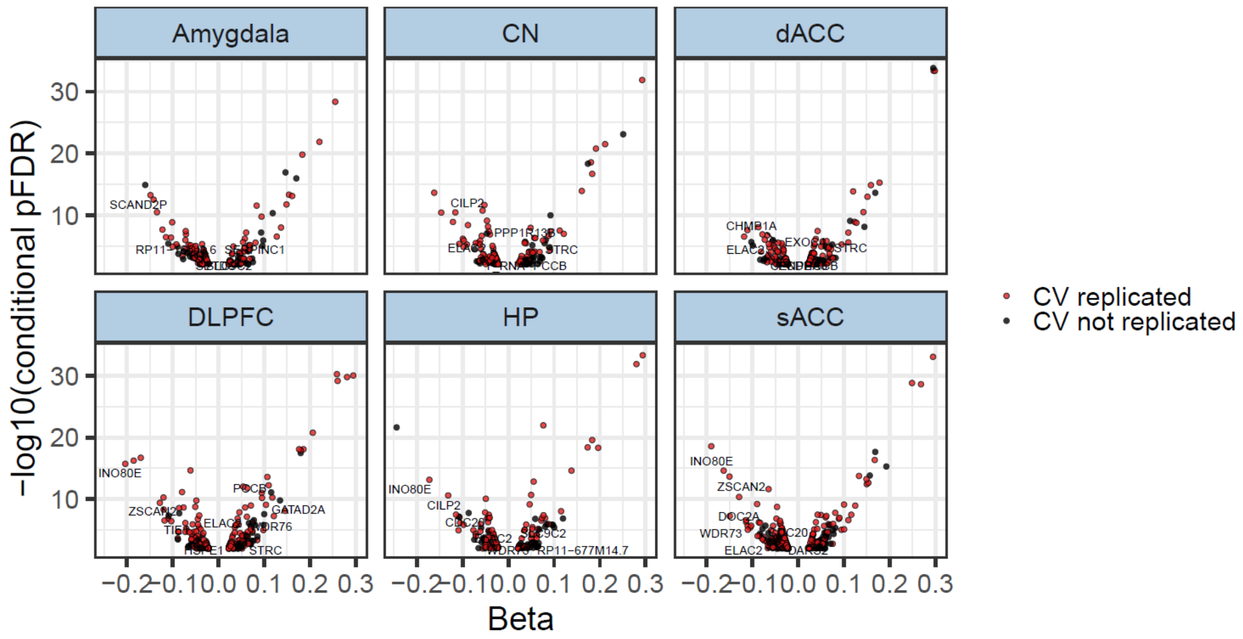


***S25. Intersection between PGC3 Fine Mapping genes and coTWAS significant genes across tissues****. Scatterplots show the relationship between effect size (BETA) and statistical significance ($-\log_{10}$ conditional $p_{(FDR)}$) for significant gene–tissue pairs across brain regions. Each point represents a gene identified by coTWAS. Red points indicate genes with associations replicated in the leave-site-out cross-validation (CV) analysis, while black points denote non-replicated genes. Gene Symbols represent the intersection of genes between the sets.*

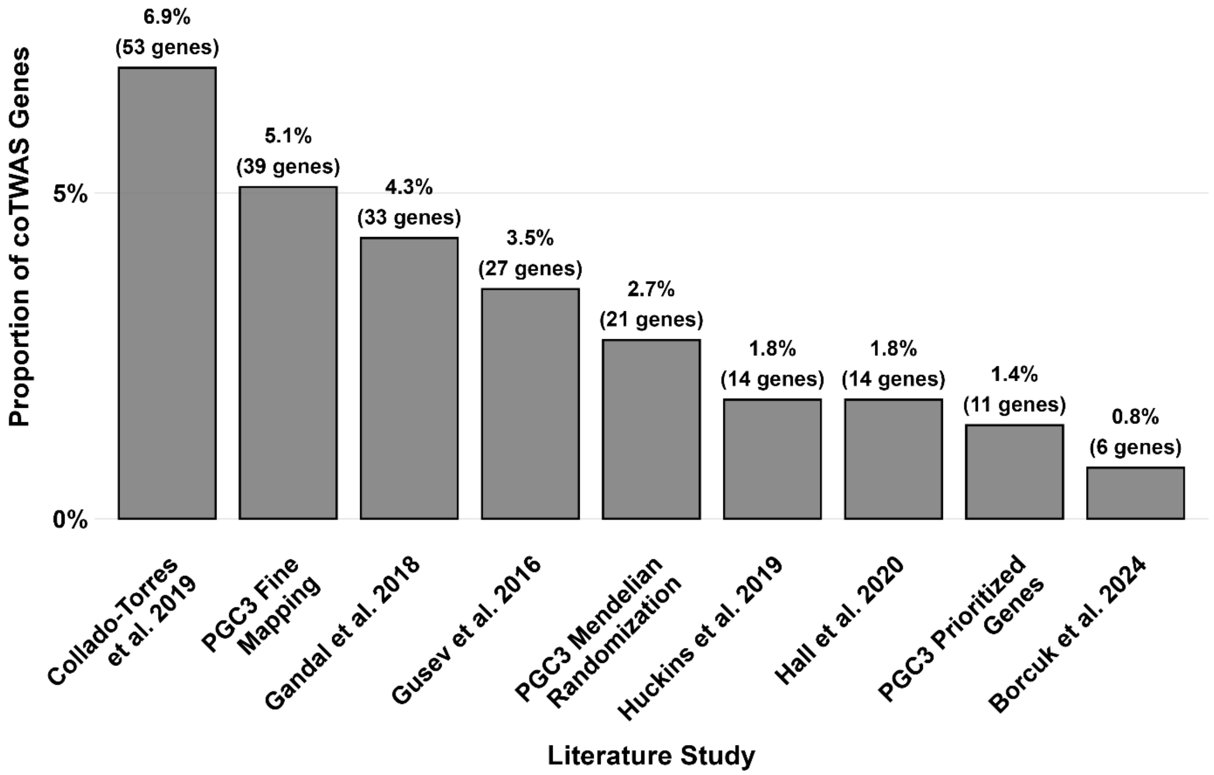


***S26. Overlap of coTWAS-identified genes with previously reported SCZ-associated genes.*** *Bar plot showing the proportion (y-axis) of coTWAS-significant genes overlapping with genes identified in published SCZ studies (x-axis). Percentages and corresponding gene counts are indicated above each bar.*

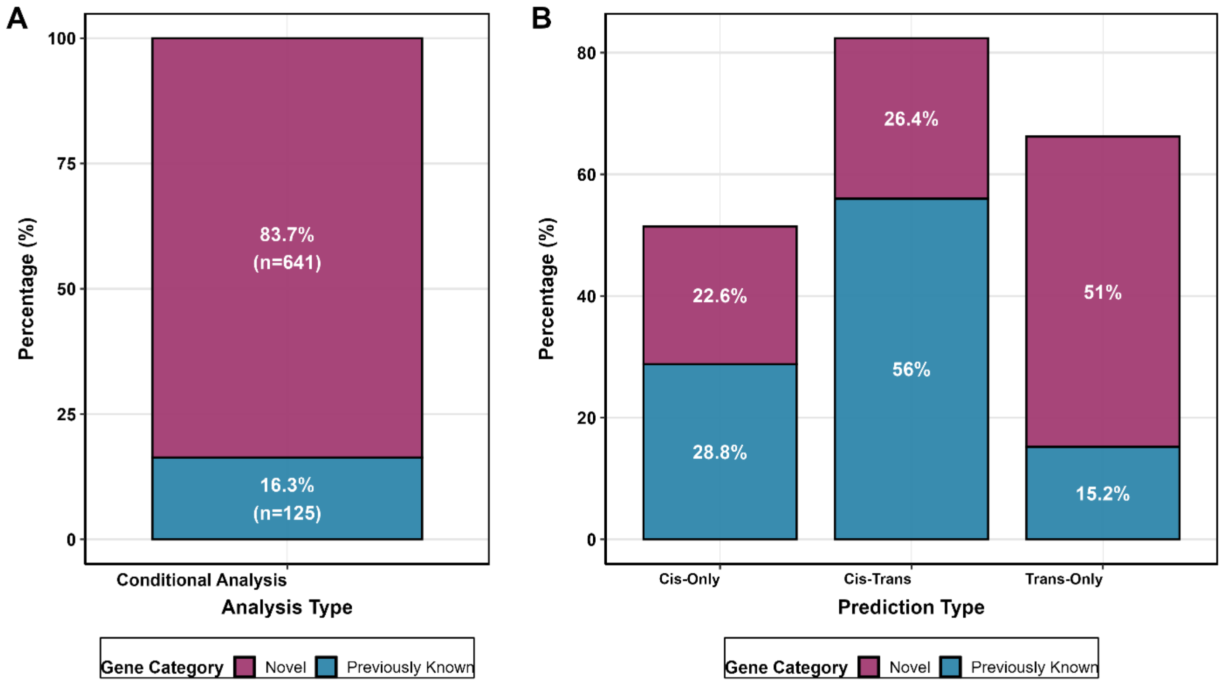


***S27. Contribution of prediction type to novel vs previously reported SCZ associations. A)*** *Proportion of novel and previously known genes identified by coTWAS.* ***B)*** *Stratification of discoveries by prediction type (cis-only, cis+trans, trans-only). Previously known genes were largely recovered through cis-only and cis+trans models, whereas novel associations were strongly enriched in the trans-only category.*

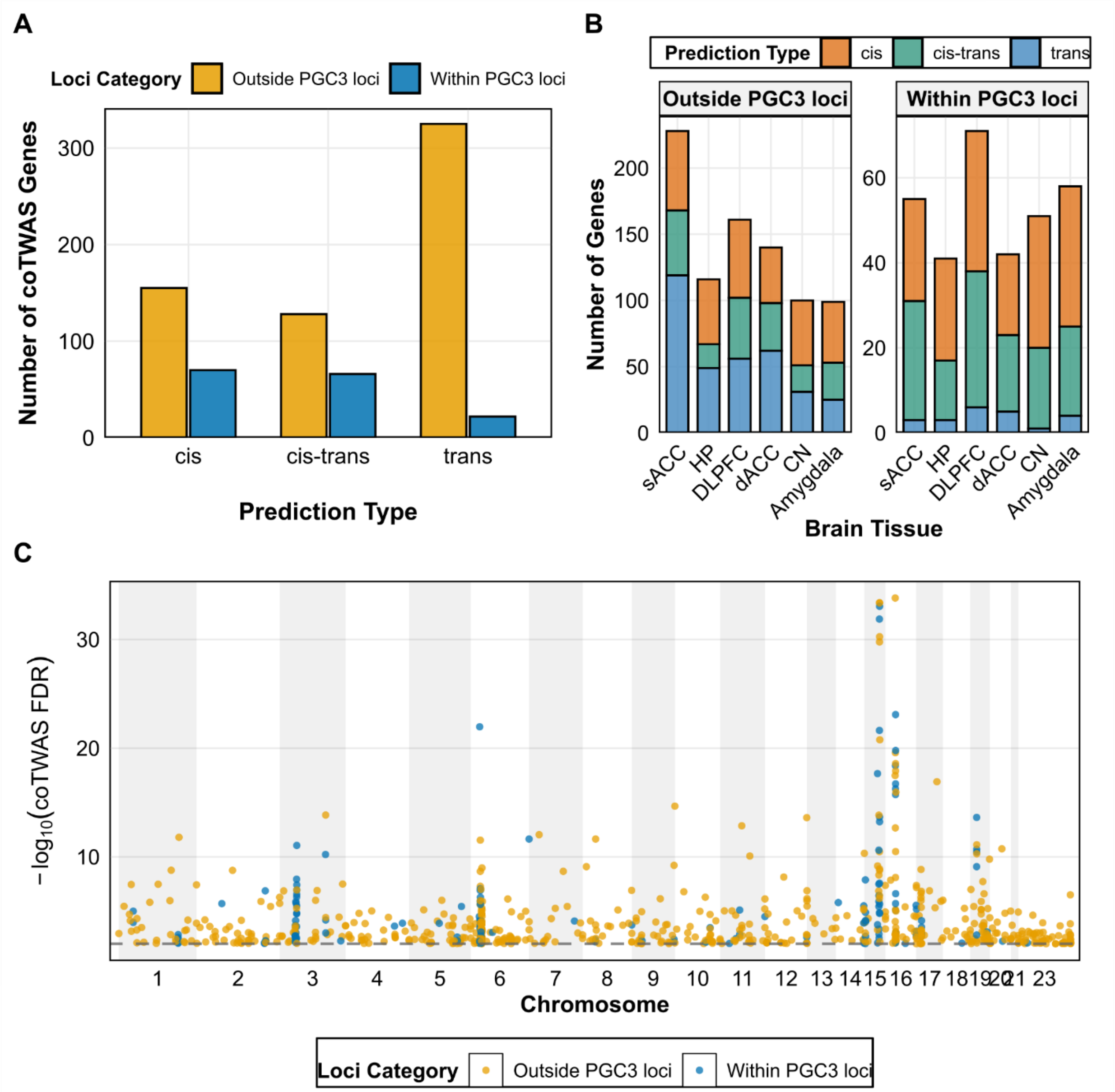


***S28. Distribution of coTWAS associations within and outside PGC3 schizophrenia loci.*** *A) Number of significant coTWAS genes stratified by prediction type (cis, cis–trans, trans) and their genomic position relative to genome-wide significant PGC3 loci[3]. B) Regional distribution of coTWAS genes across six brain tissues, shown separately for genes outside (left) and within (right) PGC3 loci. C) Manhattan plot showing the chromosomal distribution of coTWAS-significant genes ($-\log_{10}$ FDR) color-coded by locus category. Blue points denote genes within PGC3 loci and yellow points those located outside.*

### Key Resource Table

| DATA | SOURCE | IDENTIFIER |
|---|---|---|
| **WGCNA Networks** | | |
| Hartl2021<br>Pergola2019<br>Pergola2023<br>Radulescu2020<br>Gandal2018<br>Fromer2016_control<br>Werling2020<br>Li2018<br>Walker2019 | Hartl et al.[19]<br>Pergola et al.[20]<br>Pergola et al.[21]<br>Radulescu et al. [22]<br>Gandal et al. [23,24]<br>Fromer et al.[2]<br>Werling et al. [25]<br>Li et al. [26]<br>Walker et al. [27] | |
| *cis*-eQTLs across 49 tissues cataloged by GTEx | | https://gtexportal.org/home/downloads/adult-gtex/qtl |
| **Post-Mortem Data** | | |
| Lieber Institute for brain development (LIBD) | CN[7], DLPFC[4,2,6], HP[5,29], amygdala[6,8], dACC[6] and sACC[8] | CN: https://erwinpaquola-lab.libd.org/caudate_eqtl/<br>DLPFC (BrainSeq Phase I): http://eqtl.brainseq.org/phase1;<br>DLPFC, HP (BrainSeq Phase II): https://eqtl.brainseq.org/phase2/ ;<br>Amygdala, sACC: https://psychencode.synapse.org/ (syn5844980) |

| | | |
|---|---|---|
| CommonMind Consortium (CMC) Knowledge Portal | https://doi.org/10.7303/syn2759792 | DLPFC: syn18097439, release 3.0.<br>ACC: syn29442240, version 6.0. |
| GTEx consortium v8 | http://www.gtexportal.org | dbGaP: Accession phs000424.v8.p2 |
| **Association Data** | | |
| PGC3 | Trubetskoy et al.[3] | https://pgc.unc.edu/for-researchers/data-access-committee/data-access-information/ |
| PGC Summary Statistics | Schizophrenia<br><br><br><br>Bipolar Disorder<br><br><br>Major Depressive Disorder | SCZ: https://figshare.com/articles/dataset/scz2022/19426775?file=34865091;<br>BP: https://figshare.com/articles/dataset/PGC3_bipolar_disorder_GWAS_summary_statistics/14102594.<br>MDD: https://figshare.com/articles/dataset/mdd2018/14672085 |
| **Software** | | |
| PLINK v1.9, v2.0 | Purcell et al.[30]<br>Chang et al. [31] | http://pngu.mgh.harvard.edu/purcell/plink/ |
| MAGMA v1.09b | De Leeuw et al.[13] | |
| MetaXcan | Barbeira et al. [32] | https://github.com/hakyim-lab/MetaXcan |
| R (v4.3.3) | R Core Team | https://www.r-project.org/ |
| Bioconductor packages | Bioconductor | https://www.bioconductor.org/ |

| | | |
|---|---|---|
| EpiXcan | Zhang et al.[28] | https://bitbucket.org/rous-soslab/epixcan/src/master/ |
