## Supplementary Tables for "Co-expression-based models improve eQTL predictions and highlight novel transcriptome-wide genes associated with schizophrenia"

| Region | N° of genes regulated by co-expression partner *cis*-SNPs |
|---|---|
| Amygdala | 15.1% |
| CN | 10.3% |
| dACC | 24.3% |
| DLPFC | 25.4% |
| HP | 6.3% |
| sACC | 29.6% |

***Table S1. Percentage of MODULE genes predicted by cis-SNPs of co-expression partners across brain regions.*** *Abbreviations: CN: caudate nucleus bulk tissue data; dACC: dorsal anterior cingulate cortex; DLPFC: dorsolateral prefrontal cortex bulk tissue data; HP: hippocampus bulk tissue data; sACC: subgenual anterior cingulate cortex.*

| Model | Mean Adj.$R^2$ | SD Adj.$R^2$ | $N_{genes}$ | Download Link |
|---|---|---|---|---|
| EpiXcan | 0.043 | 0.081 | 8.137 | https://predictdb.org/categories/downloads/ |
| CMC | 0.051 | 0.09 | 5.832 | |
| MASHR | 0.034 | 0.08 | 4.158 | |
| PrediXcan | 0.072 | 0.10 | 2.828 | |

***Table S2.*** *Performance of PrediXcan family predictive models on the LIBD DLPFC dataset. Abbreviations: $N_{genes}$: Number of predicted gene.*

| Region | N° *trans*-SNPs as GTEx *cis*-eQTLs | % of *trans*-SNPs as GTEx *cis*-eQTLs | N° *cis*-eGenes (GTEx) |
|---|---|---|---|
| Amygdala | 21.549 | 0.29 | 10.314 |
| CN | 22.791 | 0.30 | 11.347 |
| dACC | 32.027 | 0.42 | 19.276 |
| DLPFC | 11.595 | 0.25 | 5.821 |
| HP | 26.306 | 0.36 | 12.834 |
| sACC | 25.778 | 0.44 | 17.896 |

***Table S3. Number and proportion of MODULE-derived trans-SNPs that are also significant cis-eQTLs in the GTEx QTL resource, and the corresponding number of cis-regulated eGenes for each brain region.*** *This analysis shows that many variants exert local cis effects while also contributing to* *trans regulation*. *Abbreviations: CN: caudate nucleus; dACC: dorsal anterior cingulate cortex; DLPFC: dorsolateral prefrontal cortex; HP: hippocampus; sACC: subgenual anterior cingulate cortex.*

| Region | Model | N° SNPs In Model | Overlapping SNPs | Pearson's Coeff. (log(OR), abs(weights)) |
|---|---|---|---|---|
| Amygdala | CIS | 68,021 | 8,481 | 0.14 |
| Amygdala | MODULE | 74,666 | 7,979 | 0.40 |
| CN | CIS | 73,151 | 8,959 | 0.15 |
| CN | EpiXcan | 67,292 | 8,245 | 0.15 |
| CN | MODULE | 72,090 | 7,698 | 0.34 |
| dACC | CIS | 67,945 | 7,856 | 0.15 |
| dACC | EpiXcan | 63,482 | 7,128 | 0.18 |
| dACC | MODULE | 69,966 | 6,545 | 0.28 |
| DLPFC | CIS | 89,748 | 12,252 | 0.15 |
| DLPFC | EpiXcan | 84,741 | 11,465 | 0.15 |
| DLPFC | MODULE | 46,889 | 5,248 | 0.41 |
| HP | CIS | 51,815 | 6,391 | 0.14 |
| HP | EpiXcan | 45,437 | 5,568 | 0.15 |
| HP | MODULE | 69,221 | 7,174 | 0.33 |
| sACC | CIS | 84,074 | 11,057 | 0.15 |
| sACC | EpiXcan | 80,271 | 10,377 | 0.14 |
| sACC | MODULE | 56,957 | 6,505 | 0.42 |

***Table S4. Number of CIS, EpiXcan and MODULE SNPs overlapping 900,090 SNPs from the PGC3 SCZ summary statistics (p-value < 0.05) across regions for GTEx validated genes.*** *Abbreviations: CN: caudate nucleus bulk tissue data; dACC: dorsal anterior cingulate cortex bulk tissue data; DLPFC: dorsolateral prefrontal cortex bulk tissue data; HP: hippocampus bulk tissue data; sACC: subgenual anterior cingulate cortex bulk tissue data; OR: Odd Ratios; abs: absolute value.*

| Model | Region | N genes | Mean pgc-ratio | Min pgc-ratio | Max pgc-ratio | Linear *p* | Spearman *p* | Perm Slope *p* | Perm Spearman *p* |
|---|---|---|---|---|---|---|---|---|---|
| CIS | AMY | 2129 | 0.57 | 0.09 | 2.0 | 0.06 | 0.23 | 0.04 | 0.21 |
| CIS | CN | 2202 | 0.56 | 0.09 | 1.98 | 0.73 | 0.68 | 0.6 | 0.65 |
| CIS | DLPFC | 3219 | 0.58 | 0.08 | 2.12 | 0.46 | 0.68 | 0.23 | 0.68 |
| CIS | HP | 1617 | 0.57 | 0.09 | 2.27 | 0.03 | 0.08 | 0.05 | 0.08 |
| CIS | sACC | 2836 | 0.57 | 0.08 | 2.13 | 0.91 | 0.68 | 0.78 | 0.7 |
| EpiXcan | CN | 2255 | 0.57 | 0.09 | 2.0 | 0.77 | 0.68 | 0.74 | 0.71 |
| EpiXcan | DLPFC | 3214 | 0.59 | 0.07 | 1.89 | 0.42 | 0.68 | 0.17 | 0.68 |
| EpiXcan | HP | 1681 | 0.59 | 0.1 | 1.98 | 0.81 | 0.95 | 0.67 | 0.94 |
| EpiXcan | sACC | 2814 | 0.58 | 0.09 | 2.23 | 0.88 | 1.0 | 0.68 | 1.0 |
| MODULE | AMY | 9767 | 0.2 | 0.13 | 0.31 | 0.91 | 0.95 | 0.18 | 0.95 |
| MODULE | CN | 6002 | 0.21 | 0.12 | 0.32 | 0.08 | 0.08 | 0.0 | 0.08 |
| MODULE | DLPFC | 5410 | 0.19 | 0.15 | 0.28 | 0.22 | 0.23 | 0.0 | 0.23 |
| MODULE | HP | 4982 | 0.21 | 0.11 | 0.32 | 0.35 | 0.95 | 0.0 | 0.94 |
| MODULE | sACC | 10718 | 0.26 | 0.12 | 0.43 | 0.01 | 0.02 | 0.0 | 0.02 |

***Supplementary Table 5. Summary of connectivity enrichment analysis across models and brain regions****. For each model–region pair, genes were binned into quintiles based on their PGC-weight metric, and connectivity to PGC3-prioritized genes was assessed. The table reports the number of genes analyzed, the range of PGC-weight ratios, and statistical results from linear trend, Spearman correlation, and permutation tests. Permutation p-values were derived from 1,000 gene-level permutations (see Supplementary Methods: Permutation Testing for Connectivity Enrichment).*

| PGC Site | N | Sex at birth: Males/Females | Cases/Controls | Ancestry |
|---|---|---|---|---|
| bep1b | 849 | 352/497 | 293/556 | EUR |
| braz2 | 433 | 238/195 | 102/331 | EUR |
| butr | 1,298 | 656/642 | 649/649 | EUR |
| celso | 3,499 | 2139/1360 | 1993/1506 | EUR |
| clz2a | 11,970 | 6532/5434 | 5063/6907 | EUR |

| | | | | |
|---|---|---|---|---|
| cogs1 | 870 | 517/353 | 401/469 | EUR |
| du2aa | 555 | 333/222 | 321/234 | EUR |
| enric | 1,274 | 735/539 | 700/574 | EUR |
| eu5me | 790 | 402/387 | 613/177 | EUR |
| eusp2 | 792 | 512/280 | 308/484 | EUR |
| eutu2 | 1,083 | 570/513 | 393/690 | EUR |
| gap1a | 281 | 166/115 | 120/161 | EUR |
| geba1 | 1,037 | 491/546 | 358/679 | EUR |
| gpc2a | 3,979 | 2264/1715 | 1923/2056 | EUR |
| gro2a | 376 | 238/138 | 210/166 | EUR |
| grtr | 290 | 238/52 | 145/145 | EUR |
| lemu | 1,032 | 690/342 | 516/516 | EUR |
| mcqul | 2,506 | 1302/1204 | 1222/1284 | EUR |
| mosc2 | 841 | 442/399 | 408/433 | EUR |
| price | 1,568 | 887/681 | 841/727 | EUR |
| rive1 | 1,521 | 836/685 | 319/1202 | EUR |
| rouen | 389 | 220/169 | 204/185 | EUR |
| sb2aa | 469 | 296/173 | 236/233 | EUR |
| serri | 454 | 241/213 | 216/238 | EUR |
| to10c | 5,961 | 3030/2931 | 961/5000 | EUR |
| uktr | 140 | 110/30 | 70/70 | EUR |
| xaber | 1,325 | 916/409 | 642/683 | EUR |
| xajsz | 2,487 | 1744/743 | 893/1594 | EUR |
| xasrb | 755 | 445/310 | 463/292 | EUR |
| xboco | 3,936 | 1923/2013 | 1778/2158 | EUR |
| xbuls | 801 | 381/420 | 193/608 | EUR |
| xcati | 614 | 466/148 | 397/217 | EUR |
| xcaws | 668 | 391/277 | 387/281 | EUR |
| xcgs3 | 3,580 | 1838/1742 | 474/3106 | EUR |
| xcims | 126 | 95/31 | 65/61 | EUR |

| | | | | |
|---|---|---|---|---|
| xclm2 | 7,425 | 4476/2949 | 3345/4080 | EUR |
| xclo3 | 4,023 | 2476/1547 | 2053/1970 | EUR |
| xcou3 | 1,200 | 654/546 | 521/679 | EUR |
| xdenm | 930 | 542/388 | 474/456 | EUR |
| xdubl | 1,100 | 423/677 | 260/840 | EUR |
| xedin | 598 | 370/228 | 319/279 | EUR |
| xegcu | 1,387 | 372/1015 | 236/1151 | EUR |
| xersw | 591 | 369/222 | 271/320 | EUR |
| xgras | 2,234 | 1437/797 | 1065/1169 | EUR |
| xirwt | 2,231 | 1374/857 | 1235/996 | EUR |
| xlacw | 394 | 344/50 | 153/241 | EUR |
| xlie2 | 401 | 219/182 | 132/269 | EUR |
| xlie5 | 878 | 550/328 | 489/389 | EUR |
| xmsaf | 458 | 280/178 | 321/137 | EUR |
| xmunc | 731 | 411/320 | 421/310 | EUR |
| xpewb | 2,158 | 1167/991 | 456/1702 | EUR |
| xpews | 381 | 222/159 | 148/233 | EUR |
| xport | 560 | 293/267 | 345/215 | EUR |
| xs234 | 4,232 | 2325/1907 | 1967/2265 | EUR |
| xswe1 | 421 | 219/202 | 213/208 | EUR |
| xswe5 | 4,348 | 2410/1938 | 1770/2578 | EUR |
| xswe6 | 2,132 | 1148/984 | 978/1154 | EUR |
| xtop8 | 746 | 401/345 | 369/377 | EUR |
| xucla | 1,117 | 693/424 | 580/537 | EUR |
| xuclo | 977 | 549/428 | 492/485 | EUR |
| xume2 | 2,032 | 974/1056 | 544/1488 | EUR |
| xzhh1 | 379 | 219/160 | 190/189 | EUR |

***Table S6. Distribution of PGC3 cohorts by site, sex at birth, diagnosis and ancestry.*** *PGC Site: Unique site identifier; N: Total number of participants at each site; Sex at birth Males/Females: Number of* *males and females individuals at birth. Cases/Controls: Number of healthy individuals and individuals with diagnosis.*

| Gene Name | Tissue | GVAR | Adjusted Beta | Adjusted OR | -log p[FDR] | CV Replicated |
|---|---|---|---|---|---|---|
| NPIPB10P | dACC | 0 | 0.3 | 1.34 | 33.81 | FALSE |
| RP11-182J1.14 | dACC | 0.36 | 0.3 | 1.34 | 33.37 | TRUE |
| RP11-182J1.13 | dACC | 0.12 | 0.3 | 1.35 | 33.37 | TRUE |
| RP11-182J1.14 | HP | 0.38 | 0.29 | 1.34 | 33.31 | TRUE |
| LINC00933 | sACC | 0.05 | 0.29 | 1.34 | 33.05 | TRUE |
| EFTUD1P1 | CN | 0 | 0.29 | 1.34 | 31.88 | TRUE |
| RP11-182J1.13 | HP | 0.14 | 0.28 | 1.32 | 31.88 | TRUE |
| ENSG00000256278 | DLPFC | 0.02 | 0.26 | 1.3 | 30.25 | TRUE |
| RP11-182J1.13 | DLPFC | 0.16 | 0.29 | 1.34 | 30.03 | TRUE |
| RP11-671M22.4 | DLPFC | 0.12 | 0.28 | 1.32 | 29.78 | TRUE |
| RP11-182J1.14 | DLPFC | 0.81 | 0.26 | 1.3 | 29.15 | TRUE |
| RP11-182J1.14 | sACC | 0.36 | 0.25 | 1.28 | 28.81 | TRUE |
| RP11-182J1.13 | sACC | 0.17 | 0.27 | 1.31 | 28.62 | TRUE |
| RP11-182J1.13 | Amygdala | 0.09 | 0.26 | 1.29 | 28.35 | TRUE |
| TBX6 | CN | 0.01 | 0.25 | 1.29 | 23.09 | FALSE |
| BTN3A2 | HP | 0.91 | 0.08 | 1.08 | 21.96 | TRUE |
| LINC00933 | Amygdala | 0.02 | 0.22 | 1.25 | 21.86 | TRUE |
| UBE2Q2L | HP | 0.18 | -0.24 | 0.78 | 21.63 | FALSE |
| ENSG00000256278 | CN | 0 | 0.21 | 1.24 | 21.48 | TRUE |
| CSPG4P12 | DLPFC | 0.2 | 0.21 | 1.23 | 20.77 | TRUE |
| LINC00933 | CN | 0 | 0.19 | 1.21 | 20.77 | TRUE |
| SMG1P5 | Amygdala | 0.21 | 0.18 | 1.2 | 19.78 | TRUE |
| RP11-231C14.4 | HP | 0.07 | 0.18 | 1.2 | 19.58 | TRUE |
| RP11-455F5.3 | sACC | 0.01 | -0.19 | 0.83 | 18.58 | TRUE |
| RP11-231C14.4 | CN | 0.04 | 0.18 | 1.2 | 18.54 | TRUE |
| SLX1B | HP | 0.03 | 0.17 | 1.19 | 18.39 | TRUE |
| SMG1P5 | CN | 0.2 | 0.17 | 1.19 | 18.32 | FALSE |
| SMG1P5 | HP | 0.13 | 0.2 | 1.22 | 18.32 | TRUE |
| SLX1B | DLPFC | 0.09 | 0.19 | 1.2 | 18.09 | TRUE |
| SMG1P5 | DLPFC | 0.16 | 0.18 | 1.19 | 18.09 | TRUE |
| RP11-347C12.1 | DLPFC | 0.31 | 0.18 | 1.2 | 17.9 | TRUE |
| HYKK | sACC | 0.06 | 0.17 | 1.18 | 17.66 | FALSE |
| NPIPB9 | DLPFC | 1 | 0.18 | 1.2 | 17.48 | FALSE |
| LRRC37A3 | Amygdala | 0.75 | 0.15 | 1.16 | 16.91 | FALSE |
| MAPK3 | DLPFC | 0.07 | -0.17 | 0.84 | 16.71 | TRUE |
| SLX1B | CN | 0.09 | 0.18 | 1.2 | 16.67 | TRUE |
| SLX1B | sACC | 0.12 | 0.17 | 1.18 | 16.35 | TRUE |
| CDIPT-AS1 | DLPFC | 0.13 | -0.19 | 0.83 | 16.25 | TRUE |
| RP11-196G11.6 | Amygdala | 0 | 0.17 | 1.19 | 15.95 | FALSE |
| INO80E | DLPFC | 0.1 | -0.2 | 0.82 | 15.73 | TRUE |
| TBX6 | sACC | 0.02 | 0.19 | 1.21 | 15.29 | FALSE |
| SLX1B | dACC | 0.03 | 0.18 | 1.19 | 15.25 | TRUE |
| RP11-455F5.3 | Amygdala | 0.02 | -0.16 | 0.85 | 14.88 | FALSE |
| RP11-231C14.4 | dACC | 0.04 | 0.16 | 1.17 | 14.85 | TRUE |
| UCK1 | DLPFC | 3.58 | -0.06 | 0.94 | 14.66 | TRUE |
| INO80E | sACC | 0.56 | -0.16 | 0.85 | 14.63 | TRUE |
| LRRC37A3 | HP | 0.66 | 0.14 | 1.15 | 14.63 | TRUE |

| | | | | | | |
|---|---|---|---|---|---|---|
| NPIPB9 | CN | 1 | 0.16 | 1.17 | 13.91 | TRUE |
| STAG1-DT | dACC | 0 | 0.12 | 1.13 | 13.84 | TRUE |
| UBE2Q2P6 | sACC | 1 | 0.16 | 1.17 | 13.83 | FALSE |
| LRRC37A3 | sACC | 0.5 | 0.13 | 1.14 | 13.77 | TRUE |
| ZSCAN2 | sACC | 0.01 | -0.15 | 0.86 | 13.66 | TRUE |
| NDUFA13 | CN | 0 | -0.16 | 0.85 | 13.63 | TRUE |
| NPIPB9 | dACC | 1 | 0.17 | 1.18 | 13.61 | FALSE |
| RP11-282O18.3 | DLPFC | 0.02 | 0.11 | 1.11 | 13.6 | TRUE |
| NPIPB10P | Amygdala | 0.01 | 0.15 | 1.17 | 13.31 | TRUE |
| SMG1P5 | sACC | 0.26 | 0.15 | 1.16 | 13.25 | FALSE |
| SCAND2P | Amygdala | 6.8 | -0.15 | 0.86 | 13.23 | TRUE |
| INO80E | HP | 0.86 | -0.17 | 0.84 | 13.15 | TRUE |
| SLX1B | Amygdala | 0.11 | 0.16 | 1.17 | 13.09 | TRUE |
| SMG1P5 | dACC | 0.16 | 0.15 | 1.16 | 12.97 | TRUE |
| RARRES3 | HP | 0.6 | 0.06 | 1.06 | 12.85 | TRUE |
| NPIPB11 | sACC | 17.97 | 0.15 | 1.17 | 12.66 | TRUE |
| UBE2Q2L | Amygdala | 0.09 | -0.14 | 0.87 | 12.5 | TRUE |
| NPIPB9 | sACC | 0.23 | 0.15 | 1.16 | 12.44 | TRUE |
| STAG1-DT | DLPFC | 0.05 | 0.11 | 1.12 | 12.26 | TRUE |
| AQP1 | DLPFC | 2 | 0.05 | 1.06 | 12.04 | TRUE |
| RP11-195C7.1 | DLPFC | 0.85 | 0.06 | 1.07 | 11.79 | TRUE |
| RP11-347C12.1 | Amygdala | 0.01 | 0.15 | 1.16 | 11.73 | TRUE |
| ELFN1 | CN | 0.08 | -0.05 | 0.95 | 11.64 | TRUE |
| RNF5P1 | sACC | 0.71 | -0.06 | 0.94 | 11.64 | TRUE |
| RP1-265C24.8 | Amygdala | 0.01 | 0.08 | 1.09 | 11.53 | TRUE |
| BTN3A2 | DLPFC | 0.31 | -0.08 | 0.92 | 11.16 | TRUE |
| CTC-559E9.6 | DLPFC | 0 | 0.12 | 1.12 | 11.11 | FALSE |
| ITIH4 | DLPFC | 0.18 | 0.09 | 1.1 | 11.05 | TRUE |
| DSN1 | CN | 0.85 | -0.06 | 0.95 | 10.74 | TRUE |
| ZNF101 | HP | 0 | 0.05 | 1.05 | 10.69 | TRUE |
| GOLGA6L10 | HP | 0.28 | -0.13 | 0.88 | 10.61 | TRUE |
| RP11-455F5.5 | dACC | 0.04 | 0.14 | 1.15 | 10.48 | TRUE |
| CSPG4P11 | Amygdala | 0.17 | -0.13 | 0.87 | 10.47 | TRUE |
| CILP2 | CN | 0.03 | -0.12 | 0.89 | 10.44 | TRUE |
| UBE2Q2L | CN | 0.02 | -0.15 | 0.86 | 10.41 | TRUE |
| UBE2Q2L | sACC | 0.12 | -0.13 | 0.88 | 10.37 | TRUE |
| RP11-133K1.12 | Amygdala | 0.01 | 0.12 | 1.13 | 10.32 | FALSE |
| CTC-412M14.6 | DLPFC | 0 | -0.12 | 0.89 | 10.28 | FALSE |
| RP11-133K1.12 | DLPFC | 0.01 | 0.12 | 1.12 | 10.28 | TRUE |
| PCCB | DLPFC | 0.05 | 0.1 | 1.1 | 10.21 | TRUE |
| FZD4 | HP | 0.39 | -0.05 | 0.95 | 10.07 | TRUE |
| RP11-282O18.3 | CN | 0.01 | 0.09 | 1.1 | 9.97 | FALSE |
| RP11-231C14.4 | DLPFC | 0.35 | 0.13 | 1.14 | 9.8 | FALSE |
| ZIM2 | DLPFC | 1 | -0.05 | 0.95 | 9.78 | TRUE |
| STAG1-DT | Amygdala | 0.02 | 0.09 | 1.1 | 9.75 | TRUE |
| UBE2Q2L | DLPFC | 0.56 | -0.13 | 0.88 | 9.41 | TRUE |
| RP11-247A12.7 | sACC | 0.17 | -0.09 | 0.91 | 9.21 | TRUE |
| NA | HP | 0 | 0.05 | 1.05 | 9.16 | TRUE |
| STAG1-DT | sACC | 0.01 | 0.1 | 1.11 | 9.13 | TRUE |
| NEIL2 | CN | 1 | -0.05 | 0.95 | 9.09 | TRUE |
| GATAD2A | DLPFC | 0.6 | 0.1 | 1.11 | 9.09 | TRUE |
| LRRC37A3 | dACC | 0.26 | 0.11 | 1.12 | 9.06 | FALSE |
| HLA-DMB | DLPFC | 0.07 | 0.07 | 1.07 | 8.98 | TRUE |

| | | | | | | |
|---|---|---|---|---|---|---|
| RP11-347C12.1 | sACC | 0.01 | 0.12 | 1.13 | 8.96 | FALSE |
| EFTUD1P1 | dACC | 0 | 0.12 | 1.13 | 8.91 | TRUE |
| DNM1P51 | CN | 0.09 | -0.12 | 0.89 | 8.89 | TRUE |
| TVP23B | Amygdala | 0.03 | -0.1 | 0.9 | 8.83 | TRUE |
| S100A5 | DLPFC | 1 | -0.05 | 0.95 | 8.77 | TRUE |
| RP11-182J1.18 | dACC | 0.06 | 0.13 | 1.14 | 8.76 | TRUE |
| LINC01158 | sACC | 1 | -0.05 | 0.95 | 8.75 | TRUE |
| TRIM56 | DLPFC | 0.32 | 0.06 | 1.06 | 8.66 | TRUE |
| ZNF602P | DLPFC | 0.21 | -0.08 | 0.93 | 8.66 | TRUE |
| RP11-247A12.7 | DLPFC | 0.18 | -0.09 | 0.92 | 8.56 | TRUE |
| RP11-247A12.7 | CN | 0.05 | -0.09 | 0.92 | 8.38 | TRUE |
| CSPG4P10 | DLPFC | 0.57 | -0.12 | 0.89 | 8.33 | TRUE |
| RBMS2 | CN | 0.65 | -0.04 | 0.96 | 8.14 | TRUE |
| TBX6 | DLPFC | 0.05 | 0.15 | 1.16 | 8.12 | TRUE |
| ENSG00000278713 | dACC | 0 | 0.14 | 1.16 | 8.1 | FALSE |
| RP11-182J1.18 | HP | 0.04 | 0.12 | 1.12 | 8.05 | TRUE |
| RP11-247A12.7 | dACC | 0.08 | -0.09 | 0.92 | 8.05 | TRUE |
| RP11-231C14.4 | Amygdala | 0.08 | 0.14 | 1.15 | 7.99 | TRUE |
| GLYCTK | CN | 0.01 | 0.05 | 1.05 | 7.94 | TRUE |
| ADAL | DLPFC | 4.24 | 0.05 | 1.05 | 7.88 | TRUE |
| MYH11 | HP | 1 | -0.05 | 0.95 | 7.85 | TRUE |
| RP11-282O18.3 | sACC | 0.07 | 0.08 | 1.08 | 7.8 | TRUE |
| RP11-247A12.7 | HP | 0.07 | -0.09 | 0.92 | 7.77 | FALSE |
| ZNF527 | DLPFC | 0.5 | -0.09 | 0.92 | 7.72 | FALSE |
| CSPG4P10 | Amygdala | 0.21 | -0.12 | 0.88 | 7.65 | FALSE |
| LRRC37A3 | DLPFC | 0.57 | 0.1 | 1.11 | 7.58 | FALSE |
| GOLGA6L5P | dACC | 0.01 | -0.11 | 0.9 | 7.57 | FALSE |
| CILP2 | HP | 0 | -0.11 | 0.89 | 7.5 | TRUE |
| ETV5 | sACC | 0.67 | 0.04 | 1.04 | 7.49 | TRUE |
| ADAMTS7P1 | CN | 0 | 0.11 | 1.12 | 7.49 | TRUE |
| RP11-182J1.18 | sACC | 0.05 | 0.12 | 1.12 | 7.49 | TRUE |
| CASQ2 | HP | 1 | -0.04 | 0.96 | 7.47 | TRUE |
| GNL2 | sACC | 1.12 | 0.05 | 1.05 | 7.45 | TRUE |
| CDC42BPA | dACC | 0.4 | 0.04 | 1.04 | 7.42 | TRUE |
| SFMBT1 | HP | 0.1 | 0.08 | 1.08 | 7.41 | TRUE |
| ZNF602P | Amygdala | 0.08 | -0.07 | 0.93 | 7.41 | TRUE |
| STAG1-DT | CN | 0.03 | -0.04 | 0.96 | 7.39 | TRUE |
| SNRPA1 | DLPFC | 0.54 | -0.04 | 0.96 | 7.37 | TRUE |
| RP11-333E1.1 | CN | 0.07 | 0.09 | 1.1 | 7.37 | TRUE |
| GOLGA6L5P | DLPFC | 0.08 | -0.11 | 0.9 | 7.34 | FALSE |
| ZNF603P | sACC | 0.72 | 0.06 | 1.06 | 7.29 | TRUE |
| MAPK3 | sACC | 0.85 | -0.15 | 0.86 | 7.27 | TRUE |
| UBE2Q2P6 | DLPFC | 0.52 | 0.12 | 1.13 | 7.27 | TRUE |
| GOLGA6L5P | HP | 0.08 | -0.11 | 0.9 | 7.19 | FALSE |
| ZNF603P | Amygdala | 0.25 | 0.06 | 1.06 | 7.17 | TRUE |
| CTC-524C5.2 | Amygdala | 0.18 | 0.09 | 1.1 | 7.17 | FALSE |
| ZNF603P | DLPFC | 0.65 | 0.06 | 1.06 | 7.16 | TRUE |
| UBE2Q2P6 | dACC | 1 | 0.11 | 1.12 | 7.16 | TRUE |
| CSPG4P11 | HP | 0.15 | -0.11 | 0.9 | 7.08 | FALSE |
| ZNF192P1 | sACC | 0.33 | 0.06 | 1.06 | 7.06 | TRUE |
| HCG4P5 | CN | 0.13 | -0.05 | 0.95 | 7.01 | FALSE |
| SGTA | sACC | 0.31 | 0.09 | 1.09 | 6.99 | TRUE |
| RP11-282O18.3 | HP | 0 | -0.04 | 0.96 | 6.99 | TRUE |

| | | | | | | |
|---|---|---|---|---|---|---|
| RP11-182J1.18 | CN | 0.04 | 0.12 | 1.13 | 6.97 | TRUE |
| RP11-894J14.2 | HP | 0 | 0.07 | 1.08 | 6.95 | TRUE |
| TMEM110 | HP | 0 | 0.08 | 1.08 | 6.94 | TRUE |
| C8orf31 | HP | 1 | -0.04 | 0.96 | 6.91 | TRUE |
| PHLDB2 | HP | 1 | -0.04 | 0.96 | 6.9 | TRUE |
| ZSCAN12P1 | Amygdala | 0.01 | -0.07 | 0.93 | 6.9 | TRUE |
| DNAH10OS | dACC | 0.42 | -0.08 | 0.92 | 6.88 | TRUE |
| ARL4D | DLPFC | 0.71 | 0.04 | 1.04 | 6.87 | TRUE |
| RP11-347C12.1 | HP | 0 | 0.12 | 1.13 | 6.87 | FALSE |
| FTCDNL1 | CN | 0 | -0.04 | 0.96 | 6.87 | TRUE |
| FGD5 | HP | 0.39 | -0.05 | 0.95 | 6.86 | TRUE |
| MEX3B | DLPFC | 0.59 | -0.11 | 0.9 | 6.86 | TRUE |
| NPIPB10P | sACC | 0 | -0.05 | 0.95 | 6.86 | TRUE |
| RP11-78O7.2 | sACC | 0.03 | 0.09 | 1.09 | 6.82 | TRUE |
| ZNF192P1 | HP | 0.2 | 0.06 | 1.06 | 6.82 | FALSE |
| PDSS1 | DLPFC | 0.5 | -0.05 | 0.96 | 6.79 | TRUE |
| BTN3A2 | dACC | 0.19 | -0.07 | 0.94 | 6.74 | TRUE |
| PRPSAP2 | CN | 0.19 | 0.09 | 1.09 | 6.65 | TRUE |
| ZNF602P | CN | 0.25 | -0.06 | 0.94 | 6.65 | TRUE |
| RP11-231C14.4 | sACC | 0.06 | 0.11 | 1.12 | 6.61 | TRUE |
| SEPT1 | sACC | 0 | -0.11 | 0.89 | 6.61 | TRUE |
| CYP2G1P | DLPFC | 0.02 | 0.08 | 1.08 | 6.6 | FALSE |
| ZSCAN2 | DLPFC | 0.04 | -0.12 | 0.89 | 6.55 | TRUE |
| SNORD19 | HP | 0 | 0.07 | 1.07 | 6.54 | TRUE |
| CSPG4P10 | dACC | 2.51 | -0.12 | 0.89 | 6.52 | TRUE |
| TBX6 | Amygdala | 0.02 | 0.13 | 1.14 | 6.52 | FALSE |
| IRAK1 | sACC | 0.28 | -0.04 | 0.96 | 6.5 | TRUE |
| UBE2Q2P11 | DLPFC | 0.13 | -0.1 | 0.9 | 6.44 | TRUE |
| GOLGA2P10 | Amygdala | 0.17 | -0.11 | 0.89 | 6.43 | TRUE |
| UBE2Q2P11 | Amygdala | 0.66 | -0.1 | 0.9 | 6.35 | FALSE |
| PBRM1 | DLPFC | 0.07 | 0.07 | 1.08 | 6.31 | TRUE |
| GOLGA2P10 | sACC | 0.2 | -0.11 | 0.89 | 6.31 | TRUE |
| ZNF603P | CN | 0.19 | 0.06 | 1.06 | 6.31 | TRUE |
| ACTA2 | HP | 0.08 | -0.04 | 0.96 | 6.28 | TRUE |
| ATG4B | sACC | 0.9 | 0.04 | 1.04 | 6.28 | TRUE |
| ZSCAN23 | CN | 0.25 | 0.06 | 1.06 | 6.28 | TRUE |
| RP11-815I9.4 | sACC | 0 | -0.09 | 0.92 | 6.26 | TRUE |
| ZNF192P1 | Amygdala | 0.32 | 0.06 | 1.06 | 6.18 | TRUE |
| GOLGA2P10 | CN | 0.2 | -0.1 | 0.91 | 6.16 | TRUE |
| RP11-342A23.1 | HP | 0.05 | -0.11 | 0.9 | 6.16 | TRUE |
| CTC-412M14.6 | sACC | 0.02 | -0.08 | 0.92 | 6.15 | TRUE |
| RP11-380L11.4 | DLPFC | 0.6 | -0.07 | 0.93 | 6.15 | TRUE |
| SHC3 | dACC | 0.36 | 0.04 | 1.04 | 6.14 | TRUE |
| LINC00940 | DLPFC | 0.01 | 0.07 | 1.07 | 6.12 | FALSE |
| SEC63 | sACC | 0.27 | -0.04 | 0.96 | 6.12 | TRUE |
| RP11-380L11.4 | dACC | 0.96 | -0.07 | 0.93 | 6.01 | TRUE |
| SECTM1 | sACC | 0.42 | 0.08 | 1.08 | 6.01 | TRUE |
| ZNF602P | HP | 0.03 | -0.06 | 0.94 | 6 | TRUE |
| ELL2 | sACC | 0.42 | -0.04 | 0.96 | 6 | TRUE |
| RP11-446H18.1 | DLPFC | 1 | 0.04 | 1.04 | 6 | TRUE |
| TRIM16L | CN | 0.09 | 0.08 | 1.09 | 5.97 | FALSE |
| HIST2H2AA3 | sACC | 0.53 | 0.07 | 1.07 | 5.97 | FALSE |
| RP11-333E1.1 | HP | 0.01 | 0.09 | 1.09 | 5.97 | TRUE |

| | | | | | | |
|---|---|---|---|---|---|---|
| RP11-78O7.2 | HP | 0 | 0.08 | 1.09 | 5.97 | FALSE |
| RP11-214K3.21 | Amygdala | 0 | -0.07 | 0.93 | 5.96 | TRUE |
| AL022393.9 | dACC | 0.09 | -0.06 | 0.95 | 5.96 | FALSE |
| ZSCAN26 | DLPFC | 0.74 | -0.06 | 0.95 | 5.93 | TRUE |
| PRPSAP1 | DLPFC | 1.78 | -0.04 | 0.96 | 5.9 | TRUE |
| RP11-513I15.6 | HP | 0.01 | 0.04 | 1.04 | 5.9 | TRUE |
| CALCRL | DLPFC | 0.75 | 0.04 | 1.04 | 5.89 | TRUE |
| RGS9BP | CN | 0.01 | 0.08 | 1.08 | 5.89 | TRUE |
| GOLGA6L5P | CN | 0.03 | -0.1 | 0.91 | 5.89 | TRUE |
| GOLGA6L9 | Amygdala | 0.31 | 0.1 | 1.1 | 5.89 | FALSE |
| UBE2Q2P6 | HP | 0.06 | 0.1 | 1.1 | 5.88 | FALSE |
| SEMA3F | sACC | 0.31 | 0.07 | 1.07 | 5.88 | TRUE |
| UBE2Q2P11 | HP | 0.2 | -0.1 | 0.91 | 5.86 | FALSE |
| ZKSCAN3 | CN | 0.05 | -0.06 | 0.94 | 5.84 | TRUE |
| GOLGA6L9 | HP | 0.27 | 0.1 | 1.1 | 5.82 | FALSE |
| EFTUD1P1 | DLPFC | 0.09 | 0.1 | 1.1 | 5.82 | FALSE |
| CDC7 | DLPFC | 0.5 | 0.04 | 1.04 | 5.81 | TRUE |
| ZSCAN23 | dACC | 0.49 | 0.06 | 1.06 | 5.8 | TRUE |
| RP11-333E1.1 | DLPFC | 0.23 | 0.08 | 1.08 | 5.8 | FALSE |
| PRKD1 | DLPFC | 0.47 | 0.04 | 1.04 | 5.79 | TRUE |
| CISH | sACC | 0.42 | 0.07 | 1.07 | 5.73 | FALSE |
| HLA-DMB | dACC | 0.02 | 0.05 | 1.05 | 5.73 | TRUE |
| ING5 | sACC | 0.35 | -0.07 | 0.93 | 5.71 | FALSE |
| DOC2A | sACC | 0.3 | -0.1 | 0.9 | 5.7 | TRUE |
| ALMS1P1 | CN | 0.14 | 0.04 | 1.04 | 5.7 | FALSE |
| ZNF602P | sACC | 0.34 | -0.05 | 0.95 | 5.69 | TRUE |
| ZSCAN23 | sACC | 0.28 | 0.06 | 1.06 | 5.67 | TRUE |
| SFMBT1 | DLPFC | 0.02 | 0.07 | 1.07 | 5.66 | FALSE |
| WDR73 | sACC | 0.78 | -0.11 | 0.9 | 5.65 | TRUE |
| GOLGA2P10 | dACC | 0.37 | -0.1 | 0.9 | 5.63 | FALSE |
| RP1-97D16.1 | dACC | 0.03 | -0.06 | 0.94 | 5.63 | TRUE |
| ZSCAN31 | Amygdala | 0.09 | 0.05 | 1.06 | 5.62 | TRUE |
| GOLGA6L9 | dACC | 1.08 | 0.11 | 1.12 | 5.59 | TRUE |
| PRPSAP2 | HP | 0.65 | 0.08 | 1.08 | 5.55 | FALSE |
| PRPSAP2 | Amygdala | 0.14 | 0.08 | 1.08 | 5.54 | FALSE |
| CHMP1A | dACC | 0.33 | -0.04 | 0.96 | 5.53 | TRUE |
| PPP1R13B | CN | 0.09 | 0.04 | 1.04 | 5.49 | TRUE |
| GNL3 | CN | 0.06 | -0.07 | 0.94 | 5.49 | TRUE |
| ZNF603P | HP | 0.14 | 0.05 | 1.05 | 5.48 | TRUE |
| RP11-522N14.2 | CN | 0 | -0.07 | 0.93 | 5.47 | TRUE |
| NBPF1 | DLPFC | 0.63 | 0.04 | 1.04 | 5.45 | TRUE |
| TMEM168 | dACC | 0.62 | -0.04 | 0.96 | 5.43 | TRUE |
| LINC01470 | CN | 0.06 | 0.04 | 1.04 | 5.43 | TRUE |
| SPC24 | CN | 1 | 0.08 | 1.08 | 5.43 | FALSE |
| ATIC | DLPFC | 1 | -0.04 | 0.96 | 5.43 | TRUE |
| UBE2Q2P2 | HP | 0.07 | 0.1 | 1.11 | 5.43 | FALSE |
| CSPG4P10 | sACC | 1.78 | -0.1 | 0.9 | 5.42 | TRUE |
| KIFC3 | sACC | 0.87 | -0.04 | 0.96 | 5.39 | TRUE |
| INO80E | Amygdala | 0.11 | -0.11 | 0.9 | 5.39 | FALSE |
| NBPF1 | HP | 0.01 | -0.06 | 0.94 | 5.37 | FALSE |
| AC004448.2 | sACC | 0.02 | 0.08 | 1.08 | 5.35 | FALSE |
| CDIPT-AS1 | CN | 0 | -0.11 | 0.9 | 5.34 | TRUE |
| CYP21A1P | dACC | 0.06 | -0.06 | 0.94 | 5.32 | TRUE |

| | | | | | | |
|---|---|---|---|---|---|---|
| PTRH2 | Amygdala | 0.45 | -0.04 | 0.96 | 5.31 | TRUE |
| BCL2 | dACC | 0.4 | 0.04 | 1.04 | 5.26 | TRUE |
| GOLGA6L5P | Amygdala | 0.03 | -0.09 | 0.91 | 5.26 | TRUE |
| RP5-1157M23.2 | DLPFC | 0 | -0.07 | 0.93 | 5.26 | TRUE |
| PHTF2 | sACC | 0.26 | -0.04 | 0.97 | 5.26 | TRUE |
| ZNF192P1 | dACC | 0.4 | 0.06 | 1.06 | 5.25 | TRUE |
| RP11-73M18.7 | CN | 0.03 | -0.06 | 0.94 | 5.24 | TRUE |
| GOLGA6L10 | CN | 0.21 | -0.09 | 0.91 | 5.24 | FALSE |
| UBE2Q2P2 | dACC | 0.12 | 0.1 | 1.1 | 5.24 | TRUE |
| ZNF577 | dACC | 0.07 | 0.07 | 1.08 | 5.23 | FALSE |
| RP11-121M22.1 | sACC | 0 | 0.04 | 1.04 | 5.2 | TRUE |
| DNAH10OS | Amygdala | 0.03 | -0.07 | 0.94 | 5.19 | TRUE |
| ITIH4 | HP | 0.88 | 0.06 | 1.07 | 5.18 | TRUE |
| NUP88 | sACC | 0.21 | -0.08 | 0.92 | 5.18 | FALSE |
| ZSCAN31 | HP | 1.28 | 0.05 | 1.05 | 5.17 | TRUE |
| PDCD5 | Amygdala | 0.12 | 0.07 | 1.07 | 5.15 | FALSE |
| SNORA68 | DLPFC | 0 | 0.07 | 1.07 | 5.14 | FALSE |
| QPRT | sACC | 0.17 | 0.1 | 1.11 | 5.13 | TRUE |
| BRD2 | CN | 0.04 | 0.05 | 1.05 | 5.12 | FALSE |
| PCCB | HP | 0.14 | 0.07 | 1.07 | 5.12 | FALSE |
| CCL28 | DLPFC | 1 | -0.04 | 0.96 | 5.12 | TRUE |
| RP11-73M18.8 | Amygdala | 0.03 | -0.06 | 0.94 | 5.11 | TRUE |
| FZD6 | sACC | 1.8 | 0.04 | 1.04 | 5.11 | TRUE |
| SLC43A1 | Amygdala | 0.01 | 0.07 | 1.07 | 5.1 | FALSE |
| GDPD3 | dACC | 0.21 | -0.1 | 0.91 | 5.07 | FALSE |
| CDIPT-AS1 | sACC | 0.03 | -0.11 | 0.9 | 5.06 | TRUE |
| ALMS1P1 | dACC | 0.07 | -0.05 | 0.95 | 5.06 | TRUE |
| GOLGA6L9 | sACC | 0.57 | 0.09 | 1.1 | 5.05 | FALSE |
| METAP2 | CN | 0.57 | -0.03 | 0.97 | 5.03 | TRUE |
| CD44 | HP | 0.54 | 0.04 | 1.04 | 5.03 | TRUE |
| RASSF10 | HP | 0.44 | 0.04 | 1.04 | 5.03 | TRUE |
| SCARB2 | sACC | 0.48 | -0.03 | 0.97 | 5.02 | TRUE |
| CABLES2 | HP | 0.28 | -0.04 | 0.96 | 5.01 | TRUE |
| PRPSAP2 | sACC | 0.24 | 0.08 | 1.08 | 5.01 | FALSE |
| PSORS1C1 | Amygdala | 0.17 | -0.05 | 0.95 | 5.01 | TRUE |
| KAT8 | Amygdala | 0.13 | 0.1 | 1.1 | 5.01 | FALSE |
| NPIPB11 | DLPFC | 0 | 0.1 | 1.1 | 5 | TRUE |
| CDC20 | HP | 0 | -0.08 | 0.93 | 5 | TRUE |
| PNLDC1 | dACC | 1.2 | -0.03 | 0.97 | 4.99 | TRUE |
| CFAP70 | sACC | 0.91 | -0.03 | 0.97 | 4.98 | TRUE |
| CDIPT-AS1 | HP | 0.02 | -0.11 | 0.9 | 4.97 | TRUE |
| CYP21A1P | sACC | 0.18 | -0.06 | 0.94 | 4.97 | TRUE |
| FAM214B | dACC | 0.36 | 0.03 | 1.03 | 4.95 | TRUE |
| PLN | HP | 1 | -0.04 | 0.96 | 4.94 | TRUE |
| MRPL28 | dACC | 0.56 | 0.07 | 1.07 | 4.92 | FALSE |
| ZNF603P | dACC | 0.4 | 0.05 | 1.05 | 4.92 | TRUE |
| AF064858.7 | Amygdala | 0.01 | 0.03 | 1.03 | 4.91 | TRUE |
| STRC | Amygdala | 0.06 | 0.07 | 1.07 | 4.89 | FALSE |
| BTN3A2 | Amygdala | 0.19 | -0.05 | 0.95 | 4.88 | TRUE |
| CDIPT-AS1 | Amygdala | 0.02 | -0.1 | 0.9 | 4.88 | TRUE |
| DNM1P51 | Amygdala | 0.04 | -0.09 | 0.91 | 4.88 | FALSE |
| RP11-78O7.2 | DLPFC | 0.1 | 0.07 | 1.08 | 4.87 | TRUE |
| GOLGA2P7 | CN | 0.02 | -0.1 | 0.91 | 4.86 | TRUE |

| | | | | | | |
|---|---|---|---|---|---|---|
| RP11-282O18.3 | dACC | 0 | 0.06 | 1.06 | 4.85 | FALSE |
| DNTTIP1 | sACC | 0.24 | -0.03 | 0.97 | 4.81 | TRUE |
| CYP21A1P | HP | 0.11 | -0.06 | 0.94 | 4.81 | TRUE |
| NEK4 | DLPFC | 0.4 | -0.07 | 0.94 | 4.8 | TRUE |
| RFC1 | sACC | 1.21 | -0.03 | 0.97 | 4.8 | TRUE |
| TNXA | DLPFC | 0.04 | -0.05 | 0.95 | 4.8 | TRUE |
| RP11-380L11.4 | Amygdala | 0.45 | -0.06 | 0.94 | 4.79 | TRUE |
| UBE2Q2P1 | CN | 0.04 | 0.09 | 1.09 | 4.79 | FALSE |
| NT5DC2 | dACC | 5.19 | -0.06 | 0.94 | 4.77 | TRUE |
| GOLGA6L10 | DLPFC | 0.25 | -0.09 | 0.92 | 4.75 | TRUE |
| RP11-144O23.8 | HP | 25.73 | -0.04 | 0.96 | 4.75 | FALSE |
| ANKRD27 | DLPFC | 0.21 | 0.06 | 1.07 | 4.74 | TRUE |
| ANKRD27 | dACC | 0.25 | 0.07 | 1.07 | 4.74 | TRUE |
| SLCO4C1 | sACC | 0.02 | -0.04 | 0.96 | 4.74 | TRUE |
| C4A | DLPFC | 0.03 | -0.07 | 0.93 | 4.72 | TRUE |
| CYP21A1P | DLPFC | 0.14 | -0.05 | 0.95 | 4.72 | TRUE |
| FAM167B | sACC | 0.52 | 0.07 | 1.07 | 4.71 | TRUE |
| UBE2Q2P1 | DLPFC | 0.01 | -0.09 | 0.92 | 4.71 | TRUE |
| ITIH4 | sACC | 0.13 | 0.06 | 1.06 | 4.7 | FALSE |
| CYP17A1-AS1 | Amygdala | 0.06 | 0.06 | 1.06 | 4.69 | TRUE |
| PDCD5 | sACC | 0.04 | 0.06 | 1.07 | 4.65 | TRUE |
| ZKSCAN3 | dACC | 2.88 | -0.05 | 0.95 | 4.65 | TRUE |
| DNAH10OS | DLPFC | 0.48 | -0.06 | 0.94 | 4.64 | TRUE |
| MAPK8IP1 | CN | 0.53 | -0.03 | 0.97 | 4.63 | TRUE |
| KDM6A | sACC | 0.69 | -0.04 | 0.96 | 4.63 | TRUE |
| ELAC2 | DLPFC | 0.85 | 0.04 | 1.04 | 4.6 | TRUE |
| LGALS3 | sACC | 0.13 | -0.03 | 0.97 | 4.58 | TRUE |
| PRPSAP2 | DLPFC | 0.13 | 0.07 | 1.08 | 4.58 | FALSE |
| OSGIN1 | sACC | 0.41 | 0.06 | 1.06 | 4.56 | FALSE |
| RP11-73M18.7 | Amygdala | 0.08 | -0.06 | 0.94 | 4.55 | TRUE |
| BTN3A2 | sACC | 0.37 | -0.05 | 0.95 | 4.55 | TRUE |
| SELPLG | dACC | 0.65 | 0.06 | 1.07 | 4.54 | TRUE |
| HLA-DMA | DLPFC | 0.13 | 0.05 | 1.05 | 4.52 | TRUE |
| RP11-380L11.4 | CN | 0.03 | -0.06 | 0.94 | 4.51 | TRUE |
| SHPRH | dACC | 0.5 | -0.03 | 0.97 | 4.5 | TRUE |
| CMAHP | DLPFC | 1.45 | -0.03 | 0.97 | 4.5 | TRUE |
| CCDC78 | sACC | 0.48 | -0.07 | 0.94 | 4.5 | FALSE |
| SNX19 | Amygdala | 1.13 | -0.07 | 0.93 | 4.5 | TRUE |
| ZKSCAN3 | Amygdala | 0.04 | -0.05 | 0.95 | 4.48 | TRUE |
| SPNS3 | dACC | 0.01 | 0.07 | 1.07 | 4.48 | TRUE |
| NUP88 | CN | 0.2 | -0.07 | 0.93 | 4.47 | FALSE |
| UBE2V1P2 | HP | 0.81 | -0.03 | 0.97 | 4.45 | TRUE |
| RP11-333E1.1 | Amygdala | 0.04 | 0.07 | 1.07 | 4.45 | TRUE |
| ANKRD50 | HP | 0.52 | -0.03 | 0.97 | 4.43 | TRUE |
| RP11-731C17.2 | Amygdala | 0.02 | 0.06 | 1.07 | 4.41 | TRUE |
| SCAND2P | sACC | 0.03 | -0.08 | 0.92 | 4.4 | FALSE |
| PSMB5 | dACC | 0.19 | -0.03 | 0.97 | 4.4 | TRUE |
| RP5-858L17.1 | sACC | 0.52 | 0.03 | 1.03 | 4.4 | TRUE |
| ADAMTS19 | Amygdala | 1 | -0.03 | 0.97 | 4.39 | TRUE |
| SNX19 | HP | 0.88 | -0.07 | 0.93 | 4.38 | TRUE |
| PLIN2 | sACC | 0.07 | -0.03 | 0.97 | 4.37 | TRUE |
| ZSCAN26 | Amygdala | 0 | -0.05 | 0.95 | 4.37 | TRUE |
| RP11-133K1.12 | sACC | 0.01 | 0.08 | 1.08 | 4.37 | FALSE |

| | | | | | | |
|---|---|---|---|---|---|---|
| ZSCAN23 | HP | 0.88 | 0.05 | 1.05 | 4.36 | TRUE |
| DMXL2 | Amygdala | 0.35 | -0.03 | 0.97 | 4.32 | TRUE |
| MRPL37 | sACC | 0.37 | 0.03 | 1.03 | 4.32 | TRUE |
| ESD | sACC | 0.72 | -0.03 | 0.97 | 4.32 | TRUE |
| HYKK | CN | 0.35 | 0.09 | 1.1 | 4.31 | FALSE |
| TPST1 | sACC | 0.1 | -0.03 | 0.97 | 4.31 | TRUE |
| TMEM110 | DLPFC | 0 | 0.06 | 1.06 | 4.3 | TRUE |
| PSORS1C1 | dACC | 0.09 | -0.05 | 0.95 | 4.3 | TRUE |
| CDH7 | sACC | 1 | 0.03 | 1.03 | 4.29 | TRUE |
| FAM193B | sACC | 0.26 | 0.03 | 1.03 | 4.28 | TRUE |
| RHOT2 | sACC | 0.18 | 0.03 | 1.03 | 4.26 | TRUE |
| RP11-731C17.2 | HP | 0.03 | 0.06 | 1.06 | 4.24 | FALSE |
| ZSCAN23 | DLPFC | 0.08 | 0.06 | 1.06 | 4.24 | TRUE |
| NCOA1 | sACC | 1.13 | -0.04 | 0.96 | 4.2 | TRUE |
| ALMS1P1 | Amygdala | 0.28 | -0.05 | 0.95 | 4.18 | TRUE |
| NCK1-AS1 | HP | 0.02 | 0.06 | 1.06 | 4.18 | FALSE |
| RNPEP | DLPFC | 0.4 | -0.03 | 0.97 | 4.18 | FALSE |
| FAR2P2 | DLPFC | 1 | 0.03 | 1.03 | 4.15 | TRUE |
| BTN3A3 | DLPFC | 0.37 | 0.05 | 1.05 | 4.15 | TRUE |
| CATSPER2P1 | Amygdala | 0.02 | 0.06 | 1.07 | 4.14 | FALSE |
| BSN-AS2 | DLPFC | 0.04 | -0.06 | 0.94 | 4.14 | TRUE |
| HLA-F-AS1 | CN | 0.11 | 0.05 | 1.05 | 4.14 | TRUE |
| GJA4 | sACC | 0.33 | 0.06 | 1.06 | 4.13 | FALSE |
| USP32P3 | Amygdala | 0.02 | -0.08 | 0.93 | 4.13 | FALSE |
| GOLGA6L5P | sACC | 0.06 | -0.08 | 0.92 | 4.13 | TRUE |
| BAG3 | sACC | 0.05 | -0.03 | 0.97 | 4.11 | TRUE |
| ZSCAN23 | Amygdala | 0.06 | 0.05 | 1.05 | 4.11 | TRUE |
| RP11-513M16.7 | Amygdala | 0.01 | 0.06 | 1.06 | 4.11 | TRUE |
| NKX3-1 | dACC | 4.58 | -0.04 | 0.96 | 4.1 | TRUE |
| PSORS1C1 | DLPFC | 0.19 | -0.05 | 0.96 | 4.1 | TRUE |
| HCG23 | CN | 0.63 | 0.06 | 1.06 | 4.1 | TRUE |
| EXOC4 | dACC | 0.92 | -0.03 | 0.97 | 4.09 | TRUE |
| TMEM167A | sACC | 0.64 | -0.03 | 0.97 | 4.09 | TRUE |
| WDR76 | DLPFC | 0.03 | 0.06 | 1.06 | 4.09 | TRUE |
| PGBD1 | CN | 0.63 | 0.05 | 1.05 | 4.09 | FALSE |
| UBE2Q2P11 | dACC | 0.14 | -0.08 | 0.92 | 4.09 | TRUE |
| RGS9BP | DLPFC | 0.03 | 0.06 | 1.06 | 4.06 | FALSE |
| RRN3P2 | sACC | 0.04 | -0.09 | 0.91 | 4.04 | FALSE |
| CYP17A1-AS1 | DLPFC | 0.02 | 0.05 | 1.05 | 4.04 | TRUE |
| ELAC2 | HP | 0.16 | -0.07 | 0.93 | 4.04 | TRUE |
| AC110781.3 | sACC | 0.05 | -0.05 | 0.95 | 4.03 | TRUE |
| STRC | CN | 0.03 | 0.07 | 1.07 | 4.03 | TRUE |
| PCSK5 | sACC | 2.39 | 0.03 | 1.03 | 4 | TRUE |
| TBK1 | sACC | 0.36 | -0.03 | 0.97 | 4 | TRUE |
| ZSCAN29 | DLPFC | 0.35 | 0.06 | 1.07 | 4 | TRUE |
| SNX19 | dACC | 0.36 | -0.06 | 0.94 | 4 | TRUE |
| LINC01431 | CN | 0.6 | -0.03 | 0.97 | 3.99 | FALSE |
| ZSCAN31 | CN | 0.21 | 0.04 | 1.05 | 3.99 | TRUE |
| AL022393.9 | Amygdala | 0.06 | -0.05 | 0.95 | 3.98 | TRUE |
| TIE1 | DLPFC | 0.03 | -0.06 | 0.94 | 3.96 | TRUE |
| KAT8 | DLPFC | 0.34 | 0.09 | 1.09 | 3.96 | FALSE |
| SOD3 | HP | 0.36 | -0.03 | 0.97 | 3.95 | TRUE |
| SPTBN4 | sACC | 0.5 | 0.06 | 1.06 | 3.95 | FALSE |

| | | | | | | |
|---|---|---|---|---|---|---|
| XXbac-BPG154L12.4 | CN | 0.41 | 0.05 | 1.06 | 3.95 | TRUE |
| TMEM121 | sACC | 0.34 | 0.05 | 1.06 | 3.95 | FALSE |
| GLYCTK | sACC | 0.17 | -0.08 | 0.93 | 3.93 | TRUE |
| SEMA3F | Amygdala | 0.01 | -0.05 | 0.95 | 3.93 | FALSE |
| CHIC1 | Amygdala | 0.51 | -0.03 | 0.97 | 3.93 | FALSE |
| RP5-874C20.7 | Amygdala | 0.05 | 0.04 | 1.05 | 3.92 | TRUE |
| NUP88 | Amygdala | 0.21 | -0.07 | 0.93 | 3.91 | FALSE |
| ADCY2 | sACC | 1.3 | -0.03 | 0.97 | 3.91 | TRUE |
| SLC4A2 | dACC | 1 | 0.03 | 1.03 | 3.91 | TRUE |
| APOPT1 | CN | 0.36 | 0.05 | 1.05 | 3.91 | TRUE |
| CPE | dACC | 0.37 | 0.03 | 1.03 | 3.89 | TRUE |
| ACP1 | Amygdala | 0.52 | -0.03 | 0.97 | 3.89 | TRUE |
| SUGT1 | Amygdala | 0.06 | -0.03 | 0.97 | 3.89 | TRUE |
| ZNF602P | dACC | 0.27 | -0.04 | 0.96 | 3.89 | TRUE |
| AC083843.4 | Amygdala | 0.01 | -0.07 | 0.93 | 3.89 | TRUE |
| LINC00461 | dACC | 0.43 | 0.03 | 1.03 | 3.88 | TRUE |
| ALMS1P1 | HP | 0.13 | -0.05 | 0.95 | 3.86 | TRUE |
| POLR2B | sACC | 0.53 | -0.03 | 0.97 | 3.86 | TRUE |
| FARSA | sACC | 0.38 | 0.07 | 1.07 | 3.86 | FALSE |
| SRPK3 | dACC | 0.41 | 0.03 | 1.03 | 3.84 | TRUE |
| GFOD1 | Amygdala | 1 | 0.03 | 1.03 | 3.83 | TRUE |
| PSORS1C1 | sACC | 0.12 | -0.04 | 0.96 | 3.83 | FALSE |
| TMEM126A | Amygdala | 2.2 | -0.04 | 0.96 | 3.81 | TRUE |
| CYP21A1P | Amygdala | 2.03 | -0.05 | 0.95 | 3.81 | TRUE |
| PSORS1C1 | HP | 0.02 | -0.04 | 0.96 | 3.8 | TRUE |
| MICB | Amygdala | 0.1 | 0.05 | 1.05 | 3.77 | TRUE |
| FRRS1L | DLPFC | 0.22 | 0.03 | 1.03 | 3.77 | TRUE |
| CTXN3 | HP | 0.51 | -0.03 | 0.97 | 3.76 | TRUE |
| CHRNA5 | sACC | 0.4 | -0.08 | 0.92 | 3.75 | FALSE |
| CYP21A2 | DLPFC | 0.08 | 0.05 | 1.05 | 3.75 | TRUE |
| ZSCAN16 | DLPFC | 0.15 | -0.04 | 0.96 | 3.74 | TRUE |
| NIPSNAP3B | Amygdala | 1 | -0.03 | 0.97 | 3.74 | TRUE |
| RP11-350N15.5 | dACC | 0 | 0.04 | 1.04 | 3.73 | TRUE |
| HCP5 | DLPFC | 0.25 | -0.05 | 0.95 | 3.73 | TRUE |
| SEPT1 | Amygdala | 0.17 | -0.09 | 0.92 | 3.73 | FALSE |
| FAM83H-AS1 | sACC | 0.05 | -0.05 | 0.95 | 3.73 | TRUE |
| RP11-333E1.1 | dACC | 0.02 | 0.07 | 1.07 | 3.73 | TRUE |
| MED30 | Amygdala | 0.09 | -0.03 | 0.97 | 3.71 | TRUE |
| NCAM2 | sACC | 0.28 | -0.03 | 0.97 | 3.7 | TRUE |
| C11orf95 | HP | 1 | -0.06 | 0.95 | 3.69 | FALSE |
| CELSR3-AS1 | DLPFC | 0 | 0.05 | 1.05 | 3.69 | TRUE |
| TMEM44 | dACC | 0.68 | -0.03 | 0.97 | 3.68 | TRUE |
| CSPG4P11 | DLPFC | 0.02 | -0.09 | 0.91 | 3.67 | TRUE |
| RP11-133K1.12 | HP | 0 | 0.06 | 1.06 | 3.67 | TRUE |
| ZFP36 | sACC | 0.36 | 0.06 | 1.06 | 3.67 | TRUE |
| FEM1C | sACC | 1.31 | 0.03 | 1.03 | 3.67 | TRUE |
| HAMP | sACC | 0.38 | 0.06 | 1.06 | 3.67 | FALSE |
| DNAH10OS | CN | 0.29 | -0.06 | 0.95 | 3.67 | TRUE |
| STAG2 | CN | 0.22 | -0.03 | 0.97 | 3.65 | FALSE |
| JPH1 | HP | 0.41 | 0.03 | 1.03 | 3.65 | TRUE |
| SNX19 | sACC | 1.31 | -0.06 | 0.94 | 3.64 | FALSE |
| C5orf63 | sACC | 0.06 | 0.03 | 1.03 | 3.64 | TRUE |

| | | | | | | |
|---|---|---|---|---|---|---|
| CYP17A1-AS1 | HP | 0.01 | 0.05 | 1.05 | 3.64 | TRUE |
| DUSP22 | CN | 0.34 | -0.03 | 0.97 | 3.64 | TRUE |
| RP11-694I15.7 | HP | 0.1 | -0.05 | 0.95 | 3.64 | FALSE |
| ZNF141 | Amygdala | 0.36 | 0.03 | 1.03 | 3.64 | TRUE |
| INPP4B | sACC | 0.26 | -0.03 | 0.97 | 3.62 | TRUE |
| PLEKHG2 | sACC | 0.08 | 0.06 | 1.06 | 3.6 | TRUE |
| RP11-731C17.2 | DLPFC | 0.45 | 0.06 | 1.06 | 3.59 | FALSE |
| SRSF7 | HP | 0.84 | -0.04 | 0.96 | 3.59 | TRUE |
| CCDC36 | DLPFC | 0.04 | 0.05 | 1.05 | 3.59 | FALSE |
| C10orf10 | sACC | 0.32 | -0.03 | 0.97 | 3.59 | TRUE |
| IMPA1 | Amygdala | 0.9 | -0.03 | 0.97 | 3.58 | TRUE |
| PPM1M | DLPFC | 0.64 | -0.06 | 0.95 | 3.58 | TRUE |
| LINC00342 | HP | 0.52 | -0.03 | 0.97 | 3.58 | TRUE |
| AC083843.4 | sACC | 0.01 | -0.05 | 0.95 | 3.58 | TRUE |
| SNX19 | DLPFC | 0.45 | -0.06 | 0.94 | 3.53 | TRUE |
| UBE2Q2P11 | sACC | 0.22 | -0.08 | 0.92 | 3.53 | TRUE |
| CCDC144CP | sACC | 0.55 | -0.07 | 0.94 | 3.53 | FALSE |
| MAT2B | HP | 0.32 | -0.03 | 0.97 | 3.51 | TRUE |
| PRPSAP2 | dACC | 3.11 | -0.07 | 0.94 | 3.51 | TRUE |
| ODC1 | dACC | 0.49 | 0.03 | 1.03 | 3.5 | TRUE |
| GNG12 | dACC | 0.38 | 0.03 | 1.03 | 3.5 | TRUE |
| ZNF622 | sACC | 0.08 | -0.03 | 0.97 | 3.49 | TRUE |
| RP5-1157M23.2 | HP | 0 | -0.05 | 0.95 | 3.49 | TRUE |
| BORCS7 | DLPFC | 0.31 | -0.04 | 0.96 | 3.48 | FALSE |
| RP11-890B15.3 | Amygdala | 0.1 | -0.06 | 0.94 | 3.48 | FALSE |
| CCDC189 | DLPFC | 0.75 | -0.09 | 0.92 | 3.48 | FALSE |
| ZNF204P | Amygdala | 0.41 | -0.04 | 0.96 | 3.48 | TRUE |
| AC110781.3 | HP | 0 | -0.05 | 0.95 | 3.45 | TRUE |
| DNM1P51 | HP | 0.01 | -0.07 | 0.93 | 3.45 | FALSE |
| LAMP5 | DLPFC | 0.3 | -0.03 | 0.97 | 3.45 | TRUE |
| GNL3 | DLPFC | 0.27 | -0.06 | 0.95 | 3.45 | FALSE |
| MRPS18AP1 | DLPFC | 0.07 | 0.05 | 1.05 | 3.44 | FALSE |
| TFDP1 | sACC | 3.86 | 0.03 | 1.03 | 3.44 | TRUE |
| PLCB3 | HP | 0.42 | -0.05 | 0.95 | 3.44 | FALSE |
| MLKL | sACC | 0.53 | 0.05 | 1.05 | 3.43 | FALSE |
| ATF6B | sACC | 0.46 | -0.05 | 0.95 | 3.43 | TRUE |
| RP11-731C17.2 | CN | 0.01 | 0.06 | 1.06 | 3.43 | FALSE |
| CYP21A1P | CN | 0.04 | -0.05 | 0.95 | 3.42 | TRUE |
| ITIH4 | dACC | 2.58 | 0.05 | 1.05 | 3.42 | TRUE |
| RP11-890B15.3 | DLPFC | 0.21 | -0.06 | 0.94 | 3.42 | TRUE |
| RP11-73M18.7 | HP | 0.08 | -0.05 | 0.95 | 3.41 | TRUE |
| TPP2 | CN | 0.26 | 0.03 | 1.03 | 3.39 | TRUE |
| PFN1P11 | DLPFC | 0.02 | 0.05 | 1.05 | 3.39 | TRUE |
| HLA-K | sACC | 0.71 | -0.05 | 0.96 | 3.38 | FALSE |
| DNM1P51 | DLPFC | 0.26 | 0.08 | 1.09 | 3.38 | TRUE |
| NKAPL | sACC | 0.05 | -0.04 | 0.96 | 3.37 | TRUE |
| RHBDL1 | sACC | 0.25 | -0.05 | 0.95 | 3.36 | FALSE |
| TBC1D9B | sACC | 0.46 | -0.06 | 0.94 | 3.35 | FALSE |
| IL15RA | sACC | 0.07 | -0.03 | 0.97 | 3.35 | FALSE |
| BTBD1 | Amygdala | 0.15 | -0.08 | 0.92 | 3.34 | TRUE |
| HLA-DMB | sACC | 0.14 | 0.05 | 1.05 | 3.34 | FALSE |
| C4B | HP | 1 | -0.05 | 0.95 | 3.33 | TRUE |
| HCG4P3 | DLPFC | 0.13 | 0.04 | 1.04 | 3.32 | TRUE |

| | | | | | | |
|---|---|---|---|---|---|---|
| ATOH8 | sACC | 0.06 | -0.03 | 0.97 | 3.31 | TRUE |
| HLA-C | sACC | 0.08 | 0.05 | 1.05 | 3.31 | TRUE |
| C4A | CN | 0.08 | -0.04 | 0.96 | 3.31 | TRUE |
| MB21D1 | sACC | 0.54 | -0.03 | 0.97 | 3.31 | TRUE |
| CSPG4P11 | sACC | 0.18 | -0.07 | 0.93 | 3.29 | TRUE |
| C6orf120 | Amygdala | 0.29 | -0.03 | 0.97 | 3.29 | TRUE |
| COX8C | Amygdala | 0 | -0.05 | 0.95 | 3.29 | FALSE |
| ZNRD1ASP | dACC | 0.23 | -0.05 | 0.95 | 3.29 | TRUE |
| HLA-K | HP | 0.37 | -0.05 | 0.95 | 3.29 | TRUE |
| SLC47A1P1 | Amygdala | 0.01 | -0.06 | 0.94 | 3.29 | FALSE |
| RP11-182J1.18 | DLPFC | 0.59 | 0.08 | 1.08 | 3.29 | FALSE |
| CRABP1 | DLPFC | 0.08 | 0.07 | 1.07 | 3.28 | FALSE |
| C1orf53 | dACC | 1 | 0.03 | 1.03 | 3.27 | TRUE |
| RN7SL605P | DLPFC | 0.11 | 0.05 | 1.05 | 3.27 | TRUE |
| RP11-795F19.5 | CN | 0.07 | -0.03 | 0.97 | 3.27 | TRUE |
| GLIS1 | sACC | 0.92 | -0.04 | 0.96 | 3.27 | TRUE |
| ENPEP | dACC | 1 | 0.03 | 1.03 | 3.25 | TRUE |
| MICB | CN | 0.04 | 0.04 | 1.04 | 3.25 | TRUE |
| ZMAT4 | sACC | 0.4 | -0.03 | 0.97 | 3.25 | TRUE |
| STRC | dACC | 0 | 0.07 | 1.07 | 3.24 | TRUE |
| CYB5R4 | sACC | 1.78 | -0.03 | 0.97 | 3.23 | TRUE |
| HCG23 | sACC | 0.15 | 0.05 | 1.05 | 3.23 | TRUE |
| LINC00336 | Amygdala | 0.02 | 0.04 | 1.04 | 3.23 | FALSE |
| GOLGA6L9 | CN | 1.38 | 0.07 | 1.08 | 3.22 | FALSE |
| RP11-101E13.5 | sACC | 0.22 | -0.03 | 0.97 | 3.21 | TRUE |
| CTD-2104P17.1 | DLPFC | 0.03 | 0.06 | 1.07 | 3.2 | FALSE |
| RP11-95O2.1 | DLPFC | 0.56 | -0.03 | 0.97 | 3.19 | FALSE |
| AAED1 | sACC | 1 | 0.04 | 1.04 | 3.18 | FALSE |
| MICB | HP | 0.02 | 0.05 | 1.05 | 3.17 | TRUE |
| CIDEA | DLPFC | 1.85 | 0.03 | 1.03 | 3.17 | TRUE |
| ALMS1P1 | DLPFC | 0.27 | -0.05 | 0.96 | 3.17 | TRUE |
| BTN3A2 | CN | 0.56 | -0.04 | 0.96 | 3.17 | TRUE |
| EGFL6 | CN | 0.1 | -0.03 | 0.97 | 3.16 | TRUE |
| ZNF232 | dACC | 0.35 | -0.07 | 0.93 | 3.16 | FALSE |
| RP11-798M19.6 | dACC | 0.33 | 0.03 | 1.03 | 3.16 | TRUE |
| RP11-209D14.2 | CN | 0 | 0.07 | 1.07 | 3.15 | FALSE |
| BTNL2 | HP | 6.74 | -0.05 | 0.95 | 3.15 | TRUE |
| ELAC2 | CN | 0.04 | -0.06 | 0.94 | 3.15 | TRUE |
| INPP5B | sACC | 0.17 | 0.05 | 1.06 | 3.15 | FALSE |
| DDAH2 | DLPFC | 0.36 | -0.05 | 0.95 | 3.15 | TRUE |
| HLA-F | sACC | 0.79 | -0.04 | 0.96 | 3.15 | TRUE |
| INCA1 | sACC | 0.02 | -0.07 | 0.93 | 3.14 | FALSE |
| C4A | HP | 0.05 | -0.05 | 0.96 | 3.13 | TRUE |
| DMD | dACC | 0.26 | 0.03 | 1.03 | 3.13 | TRUE |
| RP11-477N3.1 | Amygdala | 0.13 | -0.03 | 0.97 | 3.12 | TRUE |
| ZNF689 | dACC | 0.6 | 0.07 | 1.08 | 3.12 | TRUE |
| KAT8 | dACC | 0.24 | 0.08 | 1.08 | 3.11 | FALSE |
| GOLGA2P7 | dACC | 0.02 | 0.07 | 1.07 | 3.1 | FALSE |
| CPT1C | Amygdala | 0.01 | -0.05 | 0.95 | 3.09 | TRUE |
| P4HTM | sACC | 1.08 | 0.05 | 1.06 | 3.09 | FALSE |
| STRC | DLPFC | 0.05 | 0.05 | 1.06 | 3.09 | TRUE |
| ZNF792 | Amygdala | 1 | -0.06 | 0.95 | 3.09 | FALSE |
| CRISPLD2 | sACC | 0.45 | 0.05 | 1.05 | 3.09 | FALSE |

| | | | | | | |
|---|---|---|---|---|---|---|
| Y_RNA | DLPFC | 0.05 | -0.05 | 0.95 | 3.09 | FALSE |
| SERP1 | dACC | 0.22 | 0.03 | 1.03 | 3.09 | TRUE |
| CXorf40A | CN | 0.77 | -0.04 | 0.97 | 3.09 | TRUE |
| HCG4P3 | sACC | 0.06 | 0.04 | 1.04 | 3.09 | TRUE |
| ZKSCAN3 | DLPFC | 0.24 | -0.04 | 0.96 | 3.09 | TRUE |
| CYP17A1-AS1 | sACC | 0.08 | 0.04 | 1.04 | 3.09 | FALSE |
| HCG17 | HP | 0.01 | -0.05 | 0.95 | 3.08 | TRUE |
| RN7SL605P | Amygdala | 0.11 | 0.05 | 1.05 | 3.08 | FALSE |
| NUMBL | HP | 0.23 | -0.06 | 0.94 | 3.08 | FALSE |
| NDUFS4 | dACC | 0.34 | -0.03 | 0.97 | 3.06 | TRUE |
| LINC00882 | Amygdala | 1 | -0.03 | 0.97 | 3.06 | TRUE |
| FBRS | dACC | 0.19 | 0.08 | 1.09 | 3.05 | FALSE |
| CDC25A | Amygdala | 0.01 | 0.05 | 1.05 | 3.05 | FALSE |
| ABCB7 | CN | 0.26 | -0.03 | 0.97 | 3.05 | TRUE |
| RAB23 | DLPFC | 0.3 | 0.03 | 1.03 | 3.05 | TRUE |
| CDC20 | sACC | 0.02 | -0.05 | 0.95 | 3.05 | TRUE |
| MRPL28 | CN | 0.4 | 0.05 | 1.05 | 3.05 | FALSE |
| PTP4A1 | DLPFC | 0.51 | 0.03 | 1.03 | 3.05 | TRUE |
| PYDC1 | dACC | 0.01 | 0.08 | 1.08 | 3.05 | TRUE |
| NMI | sACC | 0.31 | -0.03 | 0.97 | 3.03 | TRUE |
| HIGD1A | HP | 0.03 | 0.06 | 1.06 | 3.03 | TRUE |
| KLF8 | dACC | 1.16 | 0.03 | 1.03 | 3.02 | TRUE |
| KAT8 | CN | 0.61 | 0.07 | 1.08 | 3.02 | FALSE |
| ADAMTS19 | sACC | 0.53 | 0.03 | 1.03 | 3.02 | TRUE |
| ZNF33A | CN | 0.45 | -0.03 | 0.97 | 3 | TRUE |
| MSN | HP | 0.22 | -0.03 | 0.97 | 3 | FALSE |
| XK | DLPFC | 0.27 | 0.03 | 1.03 | 2.99 | TRUE |
| SLC35G2 | Amygdala | 0.01 | 0.05 | 1.06 | 2.99 | TRUE |
| MST1R | CN | 0 | 0.05 | 1.05 | 2.99 | FALSE |
| GMPR | dACC | 0.39 | 0.03 | 1.03 | 2.99 | TRUE |
| THBD | sACC | 0.34 | 0.04 | 1.04 | 2.99 | TRUE |
| TMEM139 | sACC | 0.65 | 0.06 | 1.06 | 2.98 | FALSE |
| NOTCH4 | Amygdala | 0.19 | -0.04 | 0.96 | 2.98 | FALSE |
| ZSCAN29 | Amygdala | 0.03 | 0.05 | 1.06 | 2.98 | TRUE |
| TBC1D10B | Amygdala | 0.21 | 0.07 | 1.08 | 2.98 | FALSE |
| RP11-473C18.3 | sACC | 0.03 | -0.06 | 0.94 | 2.98 | TRUE |
| DTD1 | dACC | 0.9 | -0.03 | 0.97 | 2.97 | TRUE |
| C5AR1 | sACC | 0.07 | 0.05 | 1.05 | 2.96 | TRUE |
| DBI | dACC | 0.45 | 0.03 | 1.03 | 2.96 | TRUE |
| GLDC | sACC | 7.12 | 0.03 | 1.03 | 2.96 | TRUE |
| HLA-V | DLPFC | 0.12 | -0.04 | 0.96 | 2.96 | TRUE |
| NUP88 | dACC | 1.08 | -0.06 | 0.94 | 2.95 | FALSE |
| KCND1 | dACC | 0.42 | -0.03 | 0.97 | 2.94 | TRUE |
| SLC35E2B | Amygdala | 0.04 | -0.05 | 0.95 | 2.94 | FALSE |
| RP1-265C24.8 | CN | 0.09 | 0.04 | 1.04 | 2.94 | TRUE |
| ETV6 | sACC | 0.43 | 0.05 | 1.05 | 2.93 | TRUE |
| ZSCAN31 | sACC | 0.08 | 0.04 | 1.04 | 2.92 | TRUE |
| ZNF668 | dACC | 0 | -0.07 | 0.93 | 2.92 | FALSE |
| AC009961.3 | sACC | 0.01 | -0.03 | 0.97 | 2.91 | FALSE |
| ANKRD27 | HP | 0 | 0.05 | 1.05 | 2.91 | FALSE |
| RNF219 | sACC | 0.29 | -0.03 | 0.97 | 2.91 | TRUE |
| HLA-F-AS1 | dACC | 0.04 | -0.05 | 0.95 | 2.91 | TRUE |
| NOS2P3 | Amygdala | 0.07 | 0.07 | 1.07 | 2.9 | TRUE |

| | | | | | | |
|---|---|---|---|---|---|---|
| RAB9A | DLPFC | 0.31 | -0.03 | 0.97 | 2.9 | FALSE |
| ARID3A | sACC | 0.54 | -0.06 | 0.94 | 2.9 | FALSE |
| HLA-K | dACC | 0.53 | -0.04 | 0.96 | 2.9 | FALSE |
| RP11-158J3.2 | DLPFC | 1 | 0.03 | 1.03 | 2.89 | TRUE |
| RP11-311F12.1 | HP | 0.02 | 0.06 | 1.06 | 2.89 | FALSE |
| AP3B2 | HP | 0.07 | 0.07 | 1.07 | 2.89 | FALSE |
| SNCAIP | dACC | 0.84 | 0.03 | 1.03 | 2.88 | TRUE |
| CDK16 | CN | 0.39 | 0.03 | 1.03 | 2.88 | TRUE |
| SULT1A2 | Amygdala | 0.18 | -0.08 | 0.93 | 2.88 | FALSE |
| TIMM10 | Amygdala | 0.29 | 0.05 | 1.05 | 2.87 | FALSE |
| GUSBP5 | dACC | 0.99 | -0.05 | 0.95 | 2.87 | TRUE |
| PLXNB3 | HP | 0.5 | 0.03 | 1.03 | 2.86 | FALSE |
| ZNRD1ASP | HP | 0.1 | -0.04 | 0.96 | 2.86 | TRUE |
| BSN-AS2 | sACC | 0 | -0.05 | 0.95 | 2.86 | TRUE |
| AC110781.3 | DLPFC | 0.03 | -0.04 | 0.96 | 2.86 | TRUE |
| FBXW12 | DLPFC | 0 | 0.05 | 1.05 | 2.85 | FALSE |
| SERPINB8 | sACC | 1 | -0.03 | 0.97 | 2.85 | TRUE |
| CCDC144CP | CN | 0.26 | -0.06 | 0.95 | 2.85 | FALSE |
| RP11-380L11.4 | HP | 0.01 | -0.05 | 0.95 | 2.85 | TRUE |
| RNF7 | DLPFC | 0.2 | -0.03 | 0.97 | 2.84 | TRUE |
| KNTC1 | CN | 0.03 | -0.05 | 0.95 | 2.84 | TRUE |
| RP11-127L20.3 | Amygdala | 0.15 | -0.04 | 0.96 | 2.84 | TRUE |
| SERPINC1 | Amygdala | 0.05 | 0.05 | 1.05 | 2.84 | TRUE |
| SLC43A1 | DLPFC | 0.1 | 0.05 | 1.05 | 2.84 | FALSE |
| Y_RNA | CN | 0.01 | -0.05 | 0.95 | 2.84 | FALSE |
| HCG4P3 | dACC | 0 | 0.04 | 1.04 | 2.84 | FALSE |
| S100A11 | sACC | 0.39 | 0.04 | 1.05 | 2.84 | FALSE |
| APOPT1 | DLPFC | 0.38 | 0.05 | 1.05 | 2.83 | FALSE |
| AC004448.2 | HP | 0 | 0.06 | 1.06 | 2.82 | FALSE |
| RP11-1072A3.4 | DLPFC | 0.01 | 0.07 | 1.07 | 2.82 | FALSE |
| WDR1 | HP | 0.08 | -0.03 | 0.97 | 2.82 | TRUE |
| RP11-43A14.2 | DLPFC | 0.64 | -0.03 | 0.97 | 2.82 | TRUE |
| YJEFN3 | DLPFC | 0.76 | 0.06 | 1.06 | 2.81 | FALSE |
| HCG17 | CN | 0.03 | -0.04 | 0.96 | 2.81 | TRUE |
| ZNF589 | Amygdala | 0.38 | 0.05 | 1.05 | 2.81 | TRUE |
| FANCG | sACC | 0.46 | 0.04 | 1.04 | 2.81 | TRUE |
| AC074141.3 | sACC | 0.01 | 0.05 | 1.05 | 2.81 | FALSE |
| ZSCAN31 | dACC | 0.12 | 0.04 | 1.04 | 2.8 | TRUE |
| KRT17P6 | DLPFC | 0.04 | 0.07 | 1.07 | 2.8 | TRUE |
| JAM2 | DLPFC | 0.59 | 0.03 | 1.03 | 2.8 | TRUE |
| MAMDC2 | dACC | 0 | 0.03 | 1.03 | 2.8 | TRUE |
| VTN | sACC | 10.22 | 0.07 | 1.07 | 2.79 | FALSE |
| UBL3 | sACC | 0.62 | -0.03 | 0.97 | 2.79 | TRUE |
| ZSWIM1 | sACC | 0.24 | 0.05 | 1.06 | 2.79 | TRUE |
| MICB | DLPFC | 0.15 | 0.05 | 1.05 | 2.79 | TRUE |
| ENSG00000282772 | CN | 0 | 0.04 | 1.04 | 2.79 | FALSE |
| BRD2 | DLPFC | 0.15 | 0.05 | 1.05 | 2.79 | TRUE |
| ZNF589 | DLPFC | 0.72 | 0.05 | 1.05 | 2.78 | FALSE |
| GOLGA6L9 | DLPFC | 0.26 | 0.07 | 1.07 | 2.78 | FALSE |
| ZSCAN31 | DLPFC | 0.42 | 0.04 | 1.04 | 2.78 | TRUE |
| PSMB7 | DLPFC | 0.66 | 0.06 | 1.06 | 2.78 | FALSE |
| DLG2 | DLPFC | 0.62 | 0.03 | 1.03 | 2.78 | TRUE |
| SNX29P2 | sACC | 0.02 | 0.08 | 1.08 | 2.78 | TRUE |

| | | | | | | |
|---|---|---|---|---|---|---|
| INO80E | dACC | 0.39 | -0.08 | 0.92 | 2.77 | FALSE |
| IKBKG | DLPFC | 0.78 | 0.03 | 1.03 | 2.77 | TRUE |
| AF064858.8 | Amygdala | 0.4 | -0.04 | 0.96 | 2.77 | TRUE |
| HCG4B | CN | 0.21 | 0.05 | 1.05 | 2.77 | TRUE |
| NDUFA1 | Amygdala | 0.3 | -0.03 | 0.98 | 2.74 | TRUE |
| ZFPL1 | dACC | 0.28 | 0.05 | 1.05 | 2.74 | FALSE |
| CDC25A | DLPFC | 0.03 | 0.05 | 1.05 | 2.74 | TRUE |
| NCKIPSD | sACC | 0.37 | 0.05 | 1.05 | 2.74 | FALSE |
| HLA-K | CN | 0.47 | -0.04 | 0.96 | 2.73 | TRUE |
| USP6 | DLPFC | 0.02 | -0.06 | 0.94 | 2.73 | FALSE |
| MARCKS | dACC | 0.63 | 0.03 | 1.03 | 2.73 | TRUE |
| FCF1P2 | sACC | 0.02 | -0.05 | 0.96 | 2.72 | TRUE |
| TRNT1 | sACC | 1 | -0.03 | 0.98 | 2.72 | TRUE |
| CYP21A2 | sACC | 0.08 | 0.05 | 1.05 | 2.72 | TRUE |
| UBXN2B | sACC | 0.35 | -0.02 | 0.98 | 2.72 | TRUE |
| GUSBP5 | CN | 0.36 | -0.05 | 0.95 | 2.71 | TRUE |
| LINC00606 | DLPFC | 0 | -0.03 | 0.97 | 2.71 | TRUE |
| RP13-578N3.3 | Amygdala | 0.09 | -0.04 | 0.96 | 2.7 | TRUE |
| HLA-K | Amygdala | 0.46 | -0.04 | 0.96 | 2.7 | FALSE |
| SLC47A1 | CN | 0.53 | -0.06 | 0.95 | 2.7 | TRUE |
| ZNF764 | DLPFC | 0.82 | 0.07 | 1.07 | 2.69 | FALSE |
| CFAP36 | Amygdala | 0.13 | -0.03 | 0.97 | 2.69 | TRUE |
| CYP21A2 | HP | 0.02 | 0.05 | 1.05 | 2.68 | TRUE |
| RP11-53O19.3 | HP | 0.1 | -0.03 | 0.97 | 2.68 | TRUE |
| AC009506.1 | CN | 0.01 | 0.04 | 1.04 | 2.68 | TRUE |
| RP11-154D17.1 | CN | 0.03 | -0.02 | 0.98 | 2.68 | FALSE |
| AL022393.9 | sACC | 0.05 | -0.04 | 0.96 | 2.68 | TRUE |
| RP11-209D14.2 | dACC | 0.04 | 0.05 | 1.06 | 2.68 | FALSE |
| CXorf36 | DLPFC | 1 | -0.03 | 0.97 | 2.68 | TRUE |
| HCG23 | HP | 0.11 | 0.04 | 1.04 | 2.68 | FALSE |
| CKMT1A | sACC | 2.41 | 0.06 | 1.06 | 2.68 | FALSE |
| SETD5 | sACC | 1 | -0.03 | 0.97 | 2.68 | FALSE |
| GUSBP5 | sACC | 1.37 | -0.05 | 0.96 | 2.67 | TRUE |
| RN7SL605P | dACC | 0.09 | 0.05 | 1.05 | 2.67 | FALSE |
| GBAS | Amygdala | 0.09 | 0.02 | 1.03 | 2.67 | TRUE |
| GABRA2 | Amygdala | 0.52 | 0.03 | 1.03 | 2.67 | TRUE |
| EIF4G1 | Amygdala | 0.5 | 0.04 | 1.04 | 2.66 | TRUE |
| RP11-434D2.2 | HP | 0.01 | 0.06 | 1.06 | 2.66 | FALSE |
| RP11-115H18.1 | CN | 0.01 | 0.03 | 1.03 | 2.66 | TRUE |
| STK19 | sACC | 0.39 | -0.04 | 0.96 | 2.66 | FALSE |
| HCG4 | sACC | 0.62 | -0.04 | 0.96 | 2.66 | TRUE |
| BSN-AS2 | Amygdala | 0.01 | 0.05 | 1.05 | 2.66 | TRUE |
| SNORD3B-2 | CN | 0.05 | -0.06 | 0.94 | 2.66 | TRUE |
| SNX19 | CN | 0.3 | -0.05 | 0.95 | 2.65 | TRUE |
| GOLGA6L10 | dACC | 0.18 | -0.06 | 0.94 | 2.65 | TRUE |
| RP11-807H7.2 | sACC | 0 | -0.03 | 0.97 | 2.65 | TRUE |
| ZNF518B | DLPFC | 0.77 | 0.03 | 1.03 | 2.65 | TRUE |
| NOP58 | DLPFC | 0.46 | -0.03 | 0.97 | 2.65 | TRUE |
| RP11-644A7.2 | DLPFC | 0.49 | -0.03 | 0.97 | 2.64 | FALSE |
| NCK1-AS1 | DLPFC | 0.26 | -0.06 | 0.94 | 2.64 | FALSE |
| ARMCX4 | dACC | 0.27 | 0.03 | 1.03 | 2.63 | TRUE |
| FAM26F | HP | 2.44 | -0.03 | 0.97 | 2.63 | TRUE |
| AF064858.7 | CN | 0.02 | -0.04 | 0.96 | 2.63 | TRUE |

| | | | | | | |
|---|---|---|---|---|---|---|
| TRIM16L | dACC | 0.56 | 0.06 | 1.06 | 2.63 | FALSE |
| SNX20 | CN | 0.38 | 0.04 | 1.04 | 2.63 | FALSE |
| TTC39C | dACC | 1 | -0.02 | 0.98 | 2.62 | TRUE |
| SPINK8 | sACC | 0 | -0.05 | 0.96 | 2.62 | TRUE |
| WDR55 | HP | 0.15 | -0.03 | 0.97 | 2.62 | TRUE |
| SLC16A9 | dACC | 1.6 | 0.03 | 1.03 | 2.62 | TRUE |
| FAM60A | HP | 0.21 | -0.02 | 0.98 | 2.62 | TRUE |
| CDK2AP1 | Amygdala | 0.2 | 0.05 | 1.05 | 2.61 | FALSE |
| HLA-DQA1 | DLPFC | 0.07 | -0.04 | 0.96 | 2.61 | TRUE |
| ELAC2 | sACC | 0.91 | -0.07 | 0.94 | 2.6 | TRUE |
| STON1 | Amygdala | 0.15 | 0.02 | 1.02 | 2.6 | TRUE |
| ZNF649 | CN | 0.07 | 0.05 | 1.06 | 2.59 | FALSE |
| RP5-874C20.7 | CN | 0.05 | 0.04 | 1.04 | 2.59 | FALSE |
| RRN3P2 | dACC | 0.04 | -0.08 | 0.93 | 2.59 | FALSE |
| CYFIP2 | dACC | 0.32 | -0.03 | 0.97 | 2.58 | TRUE |
| RP11-368I7.4 | HP | 0 | -0.05 | 0.95 | 2.58 | TRUE |
| RP11-673E1.4 | dACC | 0.01 | -0.04 | 0.96 | 2.58 | TRUE |
| HLA-C | dACC | 0.27 | 0.04 | 1.04 | 2.58 | TRUE |
| RN7SL605P | CN | 0.02 | 0.05 | 1.05 | 2.58 | TRUE |
| RP5-874C20.6 | dACC | 0.01 | -0.04 | 0.97 | 2.57 | TRUE |
| ANKRD45 | CN | 0.07 | -0.04 | 0.96 | 2.57 | TRUE |
| ACP1 | HP | 0.42 | -0.03 | 0.97 | 2.57 | TRUE |
| ITPKC | sACC | 0.08 | 0.05 | 1.05 | 2.57 | FALSE |
| MCF2 | sACC | 0.68 | -0.03 | 0.97 | 2.57 | TRUE |
| NFX1 | DLPFC | 0.27 | 0.03 | 1.03 | 2.56 | TRUE |
| APLNR | HP | 0.26 | -0.05 | 0.96 | 2.56 | TRUE |
| ZNF460 | sACC | 0.56 | 0.05 | 1.05 | 2.56 | FALSE |
| APOPT1 | sACC | 2.58 | 0.05 | 1.05 | 2.56 | FALSE |
| HHIP-AS1 | sACC | 0.07 | -0.04 | 0.96 | 2.56 | FALSE |
| SAT1 | sACC | 0.27 | -0.02 | 0.98 | 2.55 | FALSE |
| NOS2P3 | DLPFC | 0.29 | 0.06 | 1.06 | 2.55 | TRUE |
| ZNF192P2 | DLPFC | 0.02 | -0.05 | 0.96 | 2.55 | TRUE |
| CCNG1 | CN | 0.18 | -0.02 | 0.98 | 2.55 | TRUE |
| TYMSOS | HP | 0.62 | 0.02 | 1.02 | 2.54 | TRUE |
| SMC1A | CN | 0.7 | 0.03 | 1.03 | 2.54 | FALSE |
| TAP2 | DLPFC | 0.53 | 0.03 | 1.04 | 2.53 | TRUE |
| UBE2Q2P2 | CN | 0.22 | 0.07 | 1.07 | 2.53 | FALSE |
| ASF1A | dACC | 0.27 | 0.02 | 1.02 | 2.53 | FALSE |
| PNISR | Amygdala | 0.29 | 0.02 | 1.02 | 2.53 | TRUE |
| C15orf40 | CN | 0.14 | -0.07 | 0.93 | 2.52 | FALSE |
| APOE | DLPFC | 0.08 | -0.05 | 0.95 | 2.52 | FALSE |
| WDR6 | CN | 0.51 | -0.04 | 0.96 | 2.52 | TRUE |
| RP11-532F6.3 | sACC | 0.07 | 0.04 | 1.04 | 2.52 | FALSE |
| TNNC1 | sACC | 0 | 0.05 | 1.05 | 2.51 | FALSE |
| EEF1A2 | DLPFC | 0.38 | -0.05 | 0.95 | 2.5 | FALSE |
| ZNRD1ASP | CN | 0.08 | -0.04 | 0.96 | 2.5 | TRUE |
| SMPD4 | sACC | 0.34 | 0.03 | 1.03 | 2.5 | TRUE |
| RP13-578N3.3 | DLPFC | 0.22 | -0.04 | 0.96 | 2.5 | TRUE |
| RP11-426C22.6 | CN | 0.07 | -0.07 | 0.94 | 2.5 | FALSE |
| TM2D2 | Amygdala | 0.53 | 0.03 | 1.03 | 2.5 | TRUE |
| PCCB | dACC | 0.7 | 0.06 | 1.07 | 2.5 | TRUE |
| SLC9C2 | Amygdala | 0.13 | 0.04 | 1.04 | 2.5 | TRUE |
| AC110781.3 | dACC | 0 | -0.04 | 0.96 | 2.5 | TRUE |

| | | | | | | |
|---|---|---|---|---|---|---|
| ZSCAN26 | sACC | 0.04 | -0.03 | 0.97 | 2.49 | FALSE |
| GYG2 | HP | 0.5 | 0.02 | 1.02 | 2.49 | TRUE |
| HLA-F-AS1 | HP | 0.01 | -0.04 | 0.96 | 2.49 | TRUE |
| HDX | DLPFC | 0.33 | -0.03 | 0.97 | 2.49 | TRUE |
| PROM1 | CN | 0.57 | 0.03 | 1.03 | 2.49 | TRUE |
| LRP4 | HP | 0.49 | 0.03 | 1.03 | 2.49 | TRUE |
| UBE3C | DLPFC | 0.45 | 0.03 | 1.03 | 2.48 | TRUE |
| RP11-127L20.3 | CN | 0.43 | -0.04 | 0.96 | 2.48 | TRUE |
| CCDC36 | CN | 0 | 0.04 | 1.04 | 2.47 | TRUE |
| XXbac-BPG154L12.4 | dACC | 0.03 | -0.04 | 0.96 | 2.47 | FALSE |
| AP5Z1 | sACC | 0.14 | 0.04 | 1.04 | 2.47 | TRUE |
| DMRTC1B | dACC | 0.66 | -0.03 | 0.97 | 2.46 | TRUE |
| MICE | HP | 0.01 | -0.04 | 0.96 | 2.46 | TRUE |
| PYROXD2 | sACC | 0.33 | 0.04 | 1.04 | 2.46 | FALSE |
| GPS2 | sACC | 0.42 | -0.05 | 0.95 | 2.45 | FALSE |
| RFX1 | sACC | 0.3 | -0.05 | 0.95 | 2.45 | FALSE |
| HLA-DQA1 | sACC | 0.2 | -0.03 | 0.97 | 2.44 | TRUE |
| CD47 | sACC | 0.19 | -0.03 | 0.97 | 2.44 | TRUE |
| BNIP3 | Amygdala | 0.27 | -0.02 | 0.98 | 2.44 | TRUE |
| DUTP6 | HP | 0 | -0.04 | 0.96 | 2.44 | TRUE |
| HLA-K | DLPFC | 0.5 | -0.04 | 0.96 | 2.44 | FALSE |
| CCDC144CP | dACC | 0.03 | -0.06 | 0.94 | 2.44 | TRUE |
| RP11-259K15.2 | dACC | 0.51 | -0.03 | 0.97 | 2.43 | TRUE |
| RRN3P2 | CN | 0.08 | -0.06 | 0.94 | 2.43 | FALSE |
| PABPC5 | Amygdala | 0.65 | 0.03 | 1.03 | 2.43 | TRUE |
| NUP88 | DLPFC | 0.39 | -0.05 | 0.95 | 2.43 | FALSE |
| CCDC36 | dACC | 0.01 | 0.05 | 1.05 | 2.42 | FALSE |
| IFITM4P | sACC | 0.01 | -0.04 | 0.96 | 2.42 | TRUE |
| MANEA-AS1 | dACC | 1.27 | 0.03 | 1.03 | 2.42 | FALSE |
| HYAL3 | sACC | 0.3 | -0.04 | 0.96 | 2.42 | FALSE |
| SLC9C2 | CN | 0.34 | 0.04 | 1.04 | 2.41 | TRUE |
| SLC9C2 | HP | 0.02 | 0.04 | 1.04 | 2.41 | TRUE |
| LAMA5 | sACC | 0.21 | 0.05 | 1.05 | 2.41 | FALSE |
| MAP7 | DLPFC | 0.04 | 0.02 | 1.02 | 2.41 | TRUE |
| ZNF668 | DLPFC | 0.63 | -0.07 | 0.94 | 2.41 | FALSE |
| TVP23B | CN | 0.87 | -0.05 | 0.95 | 2.41 | TRUE |
| MICE | DLPFC | 0.04 | -0.04 | 0.96 | 2.41 | TRUE |
| PTPA | Amygdala | 0.1 | 0.06 | 1.06 | 2.41 | FALSE |
| AC091878.1 | HP | 0.54 | 0.03 | 1.03 | 2.41 | TRUE |
| CHODL | CN | 1 | 0.02 | 1.02 | 2.41 | TRUE |
| BRCC3 | DLPFC | 0.53 | 0.03 | 1.03 | 2.41 | TRUE |
| CBX3P9 | dACC | 0.51 | 0.02 | 1.02 | 2.4 | FALSE |
| DDAH2 | HP | 0.07 | -0.04 | 0.96 | 2.4 | FALSE |
| ZNF275 | dACC | 0.27 | -0.03 | 0.97 | 2.4 | TRUE |
| RP11-51O6.1 | sACC | 0.76 | -0.02 | 0.98 | 2.4 | FALSE |
| RP13-578N3.3 | CN | 0.16 | -0.05 | 0.95 | 2.39 | TRUE |
| FXYD3 | sACC | 0.53 | 0.05 | 1.05 | 2.39 | TRUE |
| KIAA1211 | HP | 0.34 | -0.03 | 0.97 | 2.39 | TRUE |
| RP11-365P13.5 | sACC | 0.03 | 0.04 | 1.04 | 2.39 | TRUE |
| OSMR | sACC | 0.34 | -0.02 | 0.98 | 2.39 | TRUE |
| HCG9 | CN | 0.06 | 0.04 | 1.04 | 2.39 | TRUE |
| ELOVL4 | sACC | 0.77 | -0.03 | 0.98 | 2.39 | TRUE |

| | | | | | | |
|---|---|---|---|---|---|---|
| KIF13B | dACC | 0.51 | 0.03 | 1.03 | 2.39 | TRUE |
| HCG17 | DLPFC | 0.04 | -0.05 | 0.96 | 2.39 | FALSE |
| DYNC1H1 | DLPFC | 0.32 | 0.04 | 1.04 | 2.38 | FALSE |
| NPIPB11 | Amygdala | 2.93 | 0.08 | 1.08 | 2.38 | FALSE |
| PRIMPOL | Amygdala | 0.59 | 0.04 | 1.04 | 2.38 | FALSE |
| ATF6B | HP | 0.02 | -0.04 | 0.96 | 2.38 | TRUE |
| HCG4 | HP | 0.24 | -0.04 | 0.96 | 2.38 | FALSE |
| ACOT8 | sACC | 0.18 | 0.05 | 1.05 | 2.38 | FALSE |
| NCKIPSD | DLPFC | 0.14 | 0.05 | 1.05 | 2.38 | TRUE |
| VARS2 | dACC | 0.44 | -0.03 | 0.97 | 2.37 | TRUE |
| BAG4 | Amygdala | 0.29 | -0.03 | 0.97 | 2.37 | TRUE |
| CCDC144CP | DLPFC | 0.02 | -0.05 | 0.95 | 2.37 | TRUE |
| VWC2 | dACC | 0.24 | 0.02 | 1.02 | 2.37 | TRUE |
| AF064858.7 | sACC | 0.03 | -0.04 | 0.96 | 2.36 | TRUE |
| VARS2 | CN | 0.22 | -0.03 | 0.97 | 2.36 | FALSE |
| FPR1 | sACC | 0.11 | 0.05 | 1.05 | 2.36 | FALSE |
| UBE2Q2L | dACC | 0.14 | -0.07 | 0.93 | 2.36 | TRUE |
| CSNK2B | dACC | 1 | 0.03 | 1.04 | 2.36 | FALSE |
| YBX3 | sACC | 0.06 | 0.04 | 1.04 | 2.36 | FALSE |
| HCG9 | DLPFC | 0.03 | 0.04 | 1.04 | 2.36 | TRUE |
| ZNF25 | Amygdala | 0.48 | -0.03 | 0.97 | 2.36 | TRUE |
| BRD2 | dACC | 0.2 | 0.04 | 1.04 | 2.36 | TRUE |
| SOD2 | sACC | 0.14 | -0.03 | 0.97 | 2.35 | TRUE |
| TGFB1 | sACC | 0.07 | 0.05 | 1.05 | 2.35 | FALSE |
| SH3TC2 | dACC | 0.51 | 0.03 | 1.03 | 2.35 | TRUE |
| RP11-196G11.2 | HP | 0 | 0.07 | 1.07 | 2.35 | FALSE |
| HCG20 | Amygdala | 0.08 | -0.04 | 0.96 | 2.35 | FALSE |
| TRIM16L | DLPFC | 0.31 | 0.05 | 1.05 | 2.35 | FALSE |
| MAML2 | Amygdala | 0.2 | 0.03 | 1.03 | 2.35 | TRUE |
| HLA-C | DLPFC | 0.07 | 0.04 | 1.04 | 2.34 | TRUE |
| FBXL19 | dACC | 0.23 | 0.07 | 1.07 | 2.34 | TRUE |
| ITPRIP | HP | 0.01 | 0.04 | 1.04 | 2.34 | TRUE |
| SYT1 | DLPFC | 0.63 | 0.03 | 1.03 | 2.34 | TRUE |
| RP11-426C22.5 | sACC | 0.02 | -0.06 | 0.94 | 2.34 | TRUE |
| LRRTM1 | HP | 0.28 | 0.02 | 1.02 | 2.34 | TRUE |
| RP11-127L20.3 | dACC | 0.05 | -0.04 | 0.96 | 2.33 | TRUE |
| RP1-153G14.4 | dACC | 0.01 | 0.04 | 1.04 | 2.33 | TRUE |
| MICE | dACC | 0 | -0.04 | 0.96 | 2.33 | TRUE |
| CYP17A1-AS1 | CN | 0.01 | -0.04 | 0.96 | 2.33 | TRUE |
| APOPT1 | Amygdala | 2.17 | 0.05 | 1.05 | 2.33 | FALSE |
| C14orf2 | CN | 0.26 | 0.04 | 1.04 | 2.33 | FALSE |
| ZNRD1ASP | sACC | 0.13 | -0.04 | 0.96 | 2.33 | TRUE |
| RN7SL605P | HP | 0.07 | 0.04 | 1.04 | 2.33 | FALSE |
| CHI3L2 | sACC | 0.27 | -0.03 | 0.97 | 2.33 | TRUE |
| MMADHC | CN | 0.26 | -0.02 | 0.98 | 2.33 | TRUE |
| SOCS3 | sACC | 0.06 | 0.05 | 1.05 | 2.33 | FALSE |
| BSN-AS2 | dACC | 0.01 | -0.04 | 0.96 | 2.33 | TRUE |
| NCK1-AS1 | Amygdala | 0.01 | 0.04 | 1.04 | 2.33 | FALSE |
| DMXL1 | CN | 0.16 | -0.02 | 0.98 | 2.32 | TRUE |
| PROS1 | sACC | 0.18 | -0.02 | 0.98 | 2.32 | FALSE |
| AC012066.1 | dACC | 0.48 | -0.03 | 0.97 | 2.32 | FALSE |
| RP11-320H14.1 | sACC | 0.48 | -0.02 | 0.98 | 2.32 | FALSE |
| RN7SL605P | sACC | 0.21 | 0.04 | 1.04 | 2.32 | FALSE |

| | | | | | | |
|---|---|---|---|---|---|---|
| IFT74 | sACC | 0.22 | 0.02 | 1.02 | 2.32 | TRUE |
| IFITM4P | Amygdala | 0.02 | -0.04 | 0.96 | 2.32 | TRUE |
| RP11-127L20.3 | sACC | 0.1 | -0.04 | 0.96 | 2.32 | TRUE |
| TSN | HP | 0.39 | -0.03 | 0.97 | 2.32 | TRUE |
| TSHZ1 | dACC | 0.27 | 0.03 | 1.03 | 2.32 | TRUE |
| RP11-121C2.2 | sACC | 0.51 | -0.02 | 0.98 | 2.31 | TRUE |
| TP53I11 | dACC | 0.44 | -0.03 | 0.97 | 2.31 | TRUE |
| PGBD1 | HP | 0.02 | 0.03 | 1.04 | 2.31 | FALSE |
| CEP85L | DLPFC | 0.37 | 0.02 | 1.02 | 2.31 | TRUE |
| RP11-454H19.2 | HP | 0.25 | 0.02 | 1.02 | 2.31 | TRUE |
| SUSD6 | sACC | 0.37 | 0.04 | 1.04 | 2.31 | TRUE |
| CREBZF | dACC | 0.94 | -0.03 | 0.97 | 2.31 | FALSE |
| RP11-269M20.3 | DLPFC | 0 | -0.03 | 0.98 | 2.31 | FALSE |
| RP11-80H5.9 | CN | 0.04 | -0.04 | 0.96 | 2.31 | FALSE |
| TDP2 | sACC | 0.45 | -0.02 | 0.98 | 2.3 | FALSE |
| APOO | DLPFC | 0.27 | 0.02 | 1.02 | 2.3 | TRUE |
| RP11-557H15.4 | DLPFC | 0.48 | -0.04 | 0.96 | 2.3 | FALSE |
| C15orf40 | DLPFC | 0.74 | -0.07 | 0.94 | 2.3 | FALSE |
| NDUFAF5 | HP | 0.34 | -0.03 | 0.97 | 2.3 | TRUE |
| IRF7 | sACC | 0.54 | 0.05 | 1.05 | 2.3 | TRUE |
| HLA-V | sACC | 0.04 | -0.04 | 0.96 | 2.29 | TRUE |
| PCDHA5 | dACC | 0.28 | 0.03 | 1.03 | 2.29 | TRUE |
| RP11-73M18.8 | sACC | 0.75 | -0.04 | 0.96 | 2.29 | TRUE |
| GPLD1 | HP | 0.25 | -0.02 | 0.98 | 2.29 | TRUE |
| FNIP1 | sACC | 0.22 | -0.03 | 0.97 | 2.28 | TRUE |
| KCTD18 | HP | 0.29 | 0.03 | 1.03 | 2.28 | TRUE |
| BRD9 | sACC | 0.54 | -0.02 | 0.98 | 2.28 | TRUE |
| GNE | Amygdala | 0.81 | 0.03 | 1.03 | 2.28 | TRUE |
| RP11-677M14.7 | HP | 0.01 | 0.04 | 1.04 | 2.27 | TRUE |
| TATDN3 | Amygdala | 0.08 | 0.04 | 1.05 | 2.27 | FALSE |
| RP11-98I9.4 | dACC | 0.44 | 0.03 | 1.03 | 2.27 | TRUE |
| PRIMPOL | HP | 0.02 | -0.02 | 0.98 | 2.27 | FALSE |
| C4A | dACC | 1.02 | -0.04 | 0.96 | 2.27 | FALSE |
| SUMO1 | CN | 0.5 | -0.03 | 0.98 | 2.26 | TRUE |
| C4A | Amygdala | 0.26 | -0.04 | 0.96 | 2.26 | TRUE |
| MRPL28 | DLPFC | 0.31 | 0.04 | 1.05 | 2.26 | FALSE |
| REEP4 | dACC | 1.05 | -0.03 | 0.97 | 2.26 | TRUE |
| ZNF768 | HP | 0.03 | 0.06 | 1.06 | 2.26 | FALSE |
| HLA-B | CN | 0 | 0.04 | 1.04 | 2.26 | TRUE |
| HCG4P3 | CN | 0.05 | 0.04 | 1.04 | 2.26 | TRUE |
| ZDHHC15 | dACC | 0.45 | 0.02 | 1.02 | 2.26 | TRUE |
| TTC14 | HP | 0.03 | -0.02 | 0.98 | 2.26 | TRUE |
| ZNF668 | sACC | 0.21 | -0.06 | 0.94 | 2.26 | FALSE |
| C4B | dACC | 0.08 | 0.04 | 1.04 | 2.26 | TRUE |
| MICE | sACC | 0.04 | -0.05 | 0.95 | 2.26 | TRUE |
| CMTM6 | DLPFC | 0.34 | 0.02 | 1.02 | 2.25 | TRUE |
| XXbac-BPG154L12.5 | dACC | 0.03 | 0.04 | 1.04 | 2.25 | TRUE |
| DDX49 | HP | 0.03 | -0.05 | 0.95 | 2.25 | TRUE |
| HLA-F-AS1 | Amygdala | 0.04 | -0.04 | 0.96 | 2.25 | TRUE |
| SYCP2L | HP | 1 | -0.03 | 0.97 | 2.25 | TRUE |
| CCDC173 | sACC | 0.36 | 0.03 | 1.04 | 2.24 | FALSE |
| ZNF646 | HP | 0.38 | 0.06 | 1.06 | 2.24 | FALSE |

| | | | | | | |
|---|---|---|---|---|---|---|
| STAG1-DT | HP | 0.01 | 0.05 | 1.05 | 2.24 | TRUE |
| ACTR5 | sACC | 0.01 | -0.03 | 0.97 | 2.24 | TRUE |
| HIST1H2BE | DLPFC | 0.65 | -0.04 | 0.97 | 2.23 | TRUE |
| TNFAIP8L3 | sACC | 3.52 | -0.03 | 0.97 | 2.23 | TRUE |
| GDPD3 | sACC | 0.26 | -0.06 | 0.94 | 2.23 | FALSE |
| TFG | DLPFC | 0.31 | -0.02 | 0.98 | 2.23 | FALSE |
| RP11-165J3.6 | Amygdala | 0.14 | -0.03 | 0.97 | 2.23 | TRUE |
| CTD-2195M15.1 | dACC | 0.32 | -0.02 | 0.98 | 2.23 | TRUE |
| VARS2 | DLPFC | 0.35 | -0.03 | 0.97 | 2.22 | TRUE |
| CTC-471J1.11 | sACC | 2.18 | 0.04 | 1.05 | 2.22 | FALSE |
| RP11-365P13.5 | Amygdala | 0.03 | 0.04 | 1.04 | 2.22 | TRUE |
| BAG6 | CN | 0.25 | -0.04 | 0.97 | 2.22 | TRUE |
| CTB-33G10.1 | sACC | 0 | -0.05 | 0.95 | 2.22 | TRUE |
| AEBP1 | HP | 0.42 | -0.04 | 0.96 | 2.22 | FALSE |
| NME6 | dACC | 0.1 | -0.04 | 0.96 | 2.22 | FALSE |
| MIR4482 | CN | 0.03 | 0.03 | 1.03 | 2.22 | TRUE |
| HLA-V | dACC | 0.02 | -0.04 | 0.96 | 2.22 | TRUE |
| SEC14L1P1 | sACC | 1 | 0.03 | 1.03 | 2.22 | TRUE |
| HCP5 | CN | 0 | -0.03 | 0.97 | 2.21 | TRUE |
| GSTO2 | dACC | 0.32 | -0.04 | 0.96 | 2.21 | TRUE |
| MXI1 | CN | 0.11 | -0.02 | 0.98 | 2.21 | TRUE |
| HLA-DMA | HP | 0.2 | 0.03 | 1.03 | 2.21 | FALSE |
| MAEL | sACC | 0.63 | -0.03 | 0.97 | 2.21 | TRUE |
| RP11-73K9.2 | sACC | 1.58 | -0.02 | 0.98 | 2.21 | FALSE |
| ENPP4 | sACC | 0.67 | -0.03 | 0.97 | 2.21 | TRUE |
| OGT | HP | 0.35 | 0.02 | 1.02 | 2.21 | TRUE |
| MID1 | dACC | 0.62 | 0.02 | 1.02 | 2.21 | TRUE |
| BAG6 | sACC | 0.08 | -0.04 | 0.96 | 2.21 | FALSE |
| LONP1 | sACC | 0.28 | 0.05 | 1.05 | 2.2 | TRUE |
| MED16 | Amygdala | 0.07 | 0.05 | 1.05 | 2.2 | TRUE |
| LINC01351 | DLPFC | 0.29 | 0.02 | 1.02 | 2.2 | TRUE |
| DGKD | sACC | 1.55 | -0.06 | 0.95 | 2.19 | FALSE |
| TRIM26 | CN | 0.24 | 0.03 | 1.03 | 2.19 | FALSE |
| ZNF649 | DLPFC | 0.22 | 0.05 | 1.05 | 2.19 | FALSE |
| AKAP13 | dACC | 0.25 | 0.06 | 1.06 | 2.19 | FALSE |
| VARS2 | Amygdala | 0.28 | -0.04 | 0.97 | 2.18 | TRUE |
| FCHSD1 | HP | 0.62 | -0.03 | 0.97 | 2.18 | TRUE |
| ANKRD27 | CN | 0.12 | 0.04 | 1.04 | 2.18 | FALSE |
| NDUFB3 | CN | 0.44 | -0.02 | 0.98 | 2.18 | FALSE |
| FGL2 | dACC | 0.32 | -0.03 | 0.97 | 2.18 | TRUE |
| MCF2 | DLPFC | 0.01 | -0.02 | 0.98 | 2.17 | TRUE |
| SCAND2P | CN | 0.17 | -0.06 | 0.94 | 2.17 | FALSE |
| CRAT | Amygdala | 0.07 | 0.04 | 1.04 | 2.17 | FALSE |
| ELAC2 | dACC | 1.06 | -0.05 | 0.95 | 2.17 | FALSE |
| ACP1 | sACC | 0.54 | -0.02 | 0.98 | 2.17 | FALSE |
| PLPP5 | DLPFC | 3.64 | 0.03 | 1.03 | 2.17 | FALSE |
| DPYSL5 | dACC | 0.42 | -0.03 | 0.97 | 2.17 | TRUE |
| USP32P3 | DLPFC | 0.62 | -0.05 | 0.95 | 2.17 | TRUE |
| RP11-429J17.7 | sACC | 0.01 | -0.04 | 0.96 | 2.17 | TRUE |
| GPIHBP1 | Amygdala | 0.07 | 0.05 | 1.05 | 2.17 | FALSE |
| XPNPEP1 | DLPFC | 0.95 | -0.02 | 0.98 | 2.16 | FALSE |
| AC011330.13 | DLPFC | 0.01 | -0.04 | 0.96 | 2.16 | FALSE |
| P4HTM | DLPFC | 0.15 | 0.04 | 1.04 | 2.16 | FALSE |

| | | | | | | |
|---|---|---|---|---|---|---|
| PDZD4 | sACC | 0.24 | -0.02 | 0.98 | 2.16 | TRUE |
| FAM46A | DLPFC | 0.24 | -0.03 | 0.97 | 2.16 | TRUE |
| GMPR | sACC | 0.83 | -0.03 | 0.97 | 2.16 | TRUE |
| AKIRIN2 | DLPFC | 0.23 | 0.02 | 1.02 | 2.16 | TRUE |
| MTCP1 | dACC | 0.5 | -0.02 | 0.98 | 2.16 | FALSE |
| C4A | sACC | 0.66 | -0.03 | 0.97 | 2.16 | TRUE |
| PNPT1 | sACC | 0.28 | -0.02 | 0.98 | 2.16 | TRUE |
| HLA-V | CN | 0.32 | 0.05 | 1.05 | 2.16 | FALSE |
| NCKIPSD | dACC | 0.47 | 0.05 | 1.05 | 2.16 | FALSE |
| SYNPO | sACC | 0.62 | -0.03 | 0.97 | 2.16 | TRUE |
| ADAMTS6 | Amygdala | 1 | -0.02 | 0.98 | 2.15 | TRUE |
| WDR6 | DLPFC | 0.17 | -0.04 | 0.96 | 2.15 | TRUE |
| INIP | sACC | 2.67 | -0.03 | 0.97 | 2.15 | FALSE |
| CTC-524C5.2 | HP | 0.07 | 0.05 | 1.06 | 2.15 | FALSE |
| HLA-F-AS1 | sACC | 0.07 | -0.05 | 0.96 | 2.15 | TRUE |
| RPP30 | DLPFC | 0.65 | 0.03 | 1.03 | 2.15 | FALSE |
| RP1-97D16.1 | DLPFC | 0.1 | -0.03 | 0.97 | 2.14 | TRUE |
| RAB30-AS1 | sACC | 0.26 | 0.02 | 1.02 | 2.14 | FALSE |
| PNMA3 | HP | 0.2 | -0.02 | 0.98 | 2.14 | TRUE |
| ZNF75D | DLPFC | 0.44 | -0.02 | 0.98 | 2.14 | TRUE |
| RP11-127L20.3 | DLPFC | 0.05 | -0.04 | 0.96 | 2.14 | TRUE |
| BTN3A1 | dACC | 0.33 | -0.03 | 0.97 | 2.14 | FALSE |
| AZI2 | DLPFC | 0.14 | 0.03 | 1.03 | 2.14 | TRUE |
| NHS | HP | 0.27 | -0.03 | 0.97 | 2.14 | TRUE |
| FPGT | CN | 0.28 | -0.02 | 0.98 | 2.14 | FALSE |
| XXbac-BPG154L12.5 | DLPFC | 0.03 | 0.03 | 1.03 | 2.14 | TRUE |
| ZMYM3 | HP | 1 | -0.02 | 0.98 | 2.13 | TRUE |
| RP11-438B23.2 | sACC | 0.02 | -0.02 | 0.98 | 2.13 | TRUE |
| QPRT | Amygdala | 0.08 | 0.06 | 1.06 | 2.12 | FALSE |
| ACTR5 | Amygdala | 0 | 0.03 | 1.03 | 2.12 | TRUE |
| C1S | sACC | 0.25 | 0.04 | 1.04 | 2.12 | TRUE |
| RP1-97D16.1 | sACC | 0.1 | -0.03 | 0.97 | 2.12 | TRUE |
| PSORS1C1 | CN | 0.15 | -0.03 | 0.97 | 2.12 | FALSE |
| VDR | sACC | 0.79 | -0.03 | 0.97 | 2.12 | TRUE |
| RP11-365P13.5 | DLPFC | 0.04 | 0.04 | 1.04 | 2.11 | TRUE |
| SGO2 | dACC | 1 | -0.02 | 0.98 | 2.11 | FALSE |
| TROVE2 | Amygdala | 0.19 | -0.02 | 0.98 | 2.11 | FALSE |
| CENPL | dACC | 1 | -0.03 | 0.97 | 2.11 | TRUE |
| ASTN2 | sACC | 1 | -0.03 | 0.97 | 2.1 | TRUE |
| RP11-333E1.1 | sACC | 0.31 | 0.05 | 1.05 | 2.1 | FALSE |
| SRCAP | dACC | 0.2 | 0.06 | 1.07 | 2.1 | TRUE |
| TMEM106C | DLPFC | 48.48 | 0.03 | 1.03 | 2.1 | TRUE |
| RP11-128M1.1 | DLPFC | 0 | 0.03 | 1.03 | 2.1 | TRUE |
| MID1 | Amygdala | 0.51 | -0.02 | 0.98 | 2.1 | TRUE |
| HLA-F-AS1 | DLPFC | 0.09 | -0.04 | 0.96 | 2.1 | TRUE |
| NAT6 | HP | 0 | -0.04 | 0.96 | 2.1 | FALSE |
| TPMT | DLPFC | 0.69 | 0.02 | 1.03 | 2.09 | TRUE |
| NFIL3 | sACC | 0.34 | -0.03 | 0.97 | 2.09 | TRUE |
| CYB5RL | CN | 1 | 0.03 | 1.03 | 2.09 | TRUE |
| OGFOD2 | dACC | 0.04 | 0.04 | 1.04 | 2.09 | FALSE |
| FAM122B | Amygdala | 0.4 | -0.02 | 0.98 | 2.09 | FALSE |
| FOXJ2 | sACC | 0.5 | 0.04 | 1.04 | 2.09 | TRUE |

| | | | | | | |
|---|---|---|---|---|---|---|
| UGGT1 | CN | 0.56 | 0.02 | 1.02 | 2.09 | TRUE |
| RP13-942N8.1 | CN | 0.07 | 0.04 | 1.04 | 2.08 | FALSE |
| ZNF582-AS1 | sACC | 0.05 | 0.04 | 1.04 | 2.08 | FALSE |
| LTBR | sACC | 0.13 | 0.04 | 1.04 | 2.08 | FALSE |
| LIMS1 | dACC | 0.35 | 0.03 | 1.03 | 2.08 | TRUE |
| RP1-178F10.1 | dACC | 0.01 | 0.05 | 1.05 | 2.08 | TRUE |
| RP11-209D14.2 | sACC | 0.05 | 0.05 | 1.05 | 2.08 | FALSE |
| MOB4 | HP | 0.35 | -0.03 | 0.97 | 2.08 | TRUE |
| ZNF391 | Amygdala | 0.02 | 0.03 | 1.03 | 2.08 | TRUE |
| ADAMTS7 | sACC | 0.02 | -0.07 | 0.94 | 2.08 | TRUE |
| IL1RAPL1 | DLPFC | 0.44 | -0.02 | 0.98 | 2.08 | FALSE |
| NDUFAF5 | dACC | 0.54 | 0.02 | 1.02 | 2.08 | TRUE |
| PCCB | CN | 0.26 | 0.04 | 1.04 | 2.08 | TRUE |
| LINC01470 | dACC | 0 | -0.03 | 0.97 | 2.08 | FALSE |
| BOLA2 | HP | 0.01 | -0.07 | 0.94 | 2.08 | FALSE |
| ZNF682 | CN | 0.02 | 0.06 | 1.07 | 2.08 | FALSE |
| RP11-809C18.3 | dACC | 0.35 | -0.02 | 0.98 | 2.08 | TRUE |
| RPL17P43 | CN | 0.09 | -0.05 | 0.95 | 2.08 | TRUE |
| COL12A1 | sACC | 3.93 | 0.02 | 1.02 | 2.08 | TRUE |
| IGBP1 | DLPFC | 0.19 | -0.02 | 0.98 | 2.08 | FALSE |
| TRAPPC13 | DLPFC | 0.2 | -0.02 | 0.98 | 2.08 | FALSE |
| INSIG2 | dACC | 1 | 0.02 | 1.02 | 2.08 | TRUE |
| HIST1H1C | HP | 0.32 | 0.03 | 1.03 | 2.08 | FALSE |
| POLR2J4 | dACC | 0.02 | -0.04 | 0.96 | 2.08 | TRUE |
| AF064858.11 | sACC | 0.01 | -0.04 | 0.97 | 2.08 | FALSE |
| HLA-B | dACC | 0.11 | 0.03 | 1.03 | 2.07 | FALSE |
| FAM104B | Amygdala | 0.35 | 0.02 | 1.02 | 2.07 | TRUE |
| PQBP1 | dACC | 0.29 | -0.02 | 0.98 | 2.07 | TRUE |
| MARCH9 | CN | 0.51 | -0.03 | 0.97 | 2.07 | FALSE |
| HIST1H2BJ | DLPFC | 0.36 | 0.03 | 1.03 | 2.07 | TRUE |
| RP11-645C24.5 | CN | 0.07 | 0.04 | 1.04 | 2.07 | FALSE |
| BAG6 | Amygdala | 0.21 | -0.03 | 0.97 | 2.07 | TRUE |
| TDRD9 | Amygdala | 0.1 | 0.05 | 1.05 | 2.07 | FALSE |
| FLOT1 | DLPFC | 0.76 | 0.03 | 1.03 | 2.07 | TRUE |
| ZC3H12B | HP | 0.36 | 0.02 | 1.02 | 2.07 | TRUE |
| NOX4 | sACC | 0.67 | 0.03 | 1.03 | 2.07 | TRUE |
| LINC01415 | sACC | 0 | -0.03 | 0.97 | 2.06 | FALSE |
| ELF4 | sACC | 1 | -0.02 | 0.98 | 2.06 | FALSE |
| CKMT1A | Amygdala | 0.95 | 0.04 | 1.05 | 2.06 | FALSE |
| AF064858.8 | sACC | 0.24 | -0.03 | 0.97 | 2.06 | TRUE |
| PSMG1 | Amygdala | 5.53 | -0.04 | 0.97 | 2.06 | FALSE |
| C15orf40 | sACC | 0.24 | -0.06 | 0.95 | 2.06 | FALSE |
| DARS2 | sACC | 0.41 | -0.03 | 0.97 | 2.05 | TRUE |
| CHSY3 | dACC | 0.7 | -0.02 | 0.98 | 2.05 | TRUE |
| UBA6 | dACC | 0.98 | 0.02 | 1.02 | 2.05 | TRUE |
| DPYSL3 | CN | 0.26 | 0.03 | 1.03 | 2.05 | TRUE |
| AC093838.4 | dACC | 1.05 | -0.02 | 0.98 | 2.05 | FALSE |
| PRR3 | DLPFC | 0.39 | 0.03 | 1.03 | 2.05 | FALSE |
| ADD3-AS1 | sACC | 0.55 | -0.02 | 0.98 | 2.05 | FALSE |
| CEBPD | sACC | 0.28 | -0.02 | 0.98 | 2.05 | FALSE |
| SNTG2 | dACC | 0.08 | -0.02 | 0.98 | 2.05 | FALSE |
| CDKN2A | dACC | 0.83 | 0.03 | 1.03 | 2.05 | TRUE |
| RP11-5O17.1 | sACC | 0 | -0.04 | 0.96 | 2.05 | TRUE |

| | | | | | | |
|---|---|---|---|---|---|---|
| ATP11C | dACC | 1.24 | 0.02 | 1.02 | 2.04 | FALSE |
| ZNF649 | HP | 0.39 | 0.04 | 1.04 | 2.04 | FALSE |
| SNX13 | HP | 0.13 | 0.02 | 1.02 | 2.04 | TRUE |
| AP3B2 | Amygdala | 0.12 | 0.07 | 1.07 | 2.04 | FALSE |
| ABT1 | sACC | 2.85 | 0.03 | 1.03 | 2.04 | FALSE |
| SMIM13 | HP | 0.44 | -0.03 | 0.97 | 2.04 | TRUE |
| KCNC4 | HP | 0.44 | 0.03 | 1.03 | 2.04 | TRUE |
| TNFAIP6 | CN | 0.04 | -0.02 | 0.98 | 2.04 | FALSE |
| PYDC1 | sACC | 0.86 | 0.06 | 1.06 | 2.04 | FALSE |
| NUP88 | HP | 1.16 | -0.05 | 0.95 | 2.04 | FALSE |
| HLA-DRB5 | DLPFC | 0.22 | -0.04 | 0.96 | 2.04 | TRUE |
| C20orf27 | dACC | 0.2 | -0.03 | 0.97 | 2.04 | TRUE |
| TBC1D30 | Amygdala | 0.7 | 0.02 | 1.03 | 2.04 | FALSE |
| CRABP1 | sACC | 0.41 | 0.06 | 1.06 | 2.04 | FALSE |
| HLA-DQB1 | DLPFC | 0.55 | 0.04 | 1.04 | 2.04 | TRUE |
| CHIC2 | sACC | 1.2 | -0.03 | 0.97 | 2.03 | TRUE |
| WDR73 | HP | 0.72 | -0.06 | 0.95 | 2.03 | TRUE |
| HCG4 | Amygdala | 0.39 | -0.04 | 0.96 | 2.03 | TRUE |
| SETD6 | Amygdala | 0.01 | -0.03 | 0.97 | 2.03 | TRUE |
| DNAH10 | DLPFC | 0.34 | -0.04 | 0.96 | 2.03 | FALSE |
| BDNF-AS | dACC | 0.98 | -0.02 | 0.98 | 2.03 | TRUE |
| VBP1 | DLPFC | 1 | -0.02 | 0.98 | 2.03 | TRUE |
| HSPE1 | DLPFC | 0.22 | 0.02 | 1.02 | 2.03 | TRUE |
| ZNF671 | dACC | 1 | -0.04 | 0.96 | 2.03 | FALSE |
| TRIM5 | sACC | 0.65 | 0.04 | 1.04 | 2.03 | TRUE |
| MMS22L | DLPFC | 1 | -0.02 | 0.98 | 2.03 | FALSE |
| L1CAM | sACC | 1 | 0.02 | 1.02 | 2.03 | TRUE |
| GNL3LP1 | DLPFC | 0.02 | -0.03 | 0.98 | 2.03 | TRUE |
| HCG9 | Amygdala | 0.05 | 0.04 | 1.04 | 2.03 | TRUE |
| B4GALT1 | sACC | 0.36 | -0.03 | 0.97 | 2.02 | TRUE |
| RBMX2 | HP | 1 | 0.02 | 1.02 | 2.02 | TRUE |
| CSNK2B | DLPFC | 0.13 | 0.04 | 1.04 | 2.02 | FALSE |
| ABHD17AP6 | Amygdala | 0 | 0.05 | 1.05 | 2.02 | FALSE |
| G6PD | CN | 1 | -0.02 | 0.98 | 2.02 | FALSE |
| CCDC144A | DLPFC | 0.54 | 0.05 | 1.05 | 2.02 | TRUE |
| PRRC1 | sACC | 0.85 | 0.02 | 1.02 | 2.02 | TRUE |
| PRIMPOL | dACC | 0.07 | 0.04 | 1.04 | 2.02 | FALSE |
| MXI1 | dACC | 0.34 | 0.02 | 1.02 | 2.02 | TRUE |
| SLC9A6 | sACC | 0.29 | -0.02 | 0.98 | 2.02 | FALSE |
| CCSER2 | sACC | 0.23 | -0.02 | 0.98 | 2.02 | FALSE |
| PIP5K1C | CN | 0.51 | 0.05 | 1.05 | 2.02 | FALSE |
| SMS | sACC | 0.5 | -0.02 | 0.98 | 2.02 | TRUE |
| PURA | dACC | 0.37 | -0.03 | 0.97 | 2.02 | FALSE |
| POU5F1 | DLPFC | 0.05 | 0.03 | 1.03 | 2.02 | TRUE |
| TDRD9 | DLPFC | 0.08 | 0.04 | 1.04 | 2.01 | FALSE |
| EPB42 | DLPFC | 0 | -0.05 | 0.95 | 2.01 | FALSE |
| XXbac-BPG154L12.5 | CN | 0.02 | 0.04 | 1.04 | 2.01 | TRUE |
| GOPC | DLPFC | 0.85 | -0.02 | 0.98 | 2.01 | FALSE |
| WDR6 | Amygdala | 0.36 | -0.04 | 0.96 | 2.01 | FALSE |
| RUSC2 | CN | 0.14 | -0.03 | 0.97 | 2.01 | TRUE |
| PAK3 | DLPFC | 0.34 | 0.02 | 1.02 | 2.01 | FALSE |
| POU5F1 | Amygdala | 0.04 | 0.04 | 1.04 | 2.01 | TRUE |

| | | | | | | |
|---|---|---|---|---|---|---|
| COL27A1 | dACC | 0.5 | -0.03 | 0.98 | 2.01 | FALSE |
| POU3F4 | DLPFC | 0.39 | -0.02 | 0.98 | 2 | TRUE |
| RP11-420A6.2 | dACC | 0.02 | -0.05 | 0.95 | 2 | TRUE |
| SREBF1 | sACC | 0.23 | -0.05 | 0.95 | 2 | FALSE |
| CHRDL1 | DLPFC | 1 | 0.02 | 1.02 | 2 | TRUE |
| AGER | Amygdala | 0.44 | 0.03 | 1.03 | 2 | TRUE |
| RP11-73M18.8 | HP | 0.06 | -0.04 | 0.96 | 2 | TRUE |

***Table S7. coTWAS-significant genes with corresponding effect estimates and cross-validation replicability.*** *List of genes showing significant coTWAS associations (FDR < 0.01) across brain regions. Columns report: tissue (brain region of association); GVAR, the pooled effective-sample-size–weighted variance of genetically predicted expression across PGC3 cohorts, reflecting prediction stability; Adjusted β (meta-analytic regression coefficient); Adjusted OR (odds ratio derived from β); log p (−$log_{10}$ conditional $p_{(FDR)}$); and CV Replicated, indicating whether the association was confirmed in the leave-site-out cross-validation analysis (empirical one-tailed p < 0.05).*

| Region | N° genes |
|---|---|
| Amygdala | 27.287 |
| CN | 26.660 |
| dACC | 27.241 |
| DLPFC | 25.593 |
| HP | 26.993 |
| sACC | 27.254 |

***Table S8.*** *Number of genes in each brain region after processing of gene expression in the LIBD dataset.*
